# Discovery and characterization of coding and non-coding driver mutations in more than 2,500 whole cancer genomes

**DOI:** 10.1101/237313

**Authors:** Esther Rheinbay, Morten Muhlig Nielsen, Federico Abascal, Grace Tiao, Henrik Hornshøj, Julian M. Hess, Randi Istrup Pedersen, Lars Feuerbach, Radhakrishnan Sabarinathan, Tobias Madsen, Jaegil Kim, Loris Mularoni, Shimin Shuai, Andrés Lanzós, Carl Herrmann, Yosef E. Maruvka, Ciyue Shen, Samirkumar B. Amin, Johanna Bertl, Priyanka Dhingra, Klev Diamanti, Abel Gonzalez-Perez, Qianyun Guo, Nicholas J. Haradhvala, Keren Isaev, Malene Juul, Jan Komorowski, Sushant Kumar, Donghoon Lee, Lucas Lochovsky, Eric Minwei Liu, Oriol Pich, David Tamborero, Husen M. Umer, Liis Uusküla-Reimand, Claes Wadelius, Lina Wadi, Jing Zhang, Keith A. Boroevich, Joana Carlevaro-Fita, Dimple Chakravarty, Calvin W.Y. Chan, Nuno A. Fonseca, Mark P. Hamilton, Chen Hong, Andre Kahles, Youngwook Kim, Kjong-Van Lehmann, Todd A. Johnson, Abdullah Kahraman, Keunchil Park, Gordon Saksena, Lina Sieverling, Nicholas A. Sinnott-Armstrong, Peter J. Campbell, Asger Hobolth, Manolis Kellis, Michael S. Lawrence, Ben Raphael, Mark A. Rubin, Chris Sander, Lincoln Stein, Josh Stuart, Tatsuhiko Tsunoda, David A. Wheeler, Rory Johnson, Jüri Reimand, Mark B. Gerstein, Ekta Khurana, Núria López-Bigas, Iñigo Martincorena, Jakob Skou Pedersen, Gad Getz

## Abstract

Discovery of cancer drivers has traditionally focused on the identification of protein-coding genes. Here we present a comprehensive analysis of putative cancer driver mutations in both protein-coding and non-coding genomic regions across >2,500 whole cancer genomes from the Pan-Cancer Analysis of Whole Genomes (PCAWG) Consortium. We developed a statistically rigorous strategy for combining significance levels from multiple driver discovery methods and demonstrate that the integrated results overcome limitations of individual methods. We combined this strategy with careful filtering and applied it to protein-coding genes, promoters, untranslated regions (UTRs), distal enhancers and non-coding RNAs. These analyses redefine the landscape of non-coding driver mutations in cancer genomes, confirming a few previously reported elements and raising doubts about others, while identifying novel candidate elements across 27 cancer types. Novel recurrent events were found in the promoters or 5’UTRs of *TP53, RFTN1, RNF34,* and *MTG2,* in the 3’UTRs of *NFKBIZ* and *TOB1,* and in the non-coding RNA *RMRP.* We provide evidence that the previously reported non-coding RNAs *NEAT1* and *MALAT1* may be subject to a localized mutational process. Perhaps the most striking finding is the relative paucity of point mutations driving cancer in non-coding genes and regulatory elements. Though we have limited power to discover infrequent non-coding drivers in individual cohorts, combined analysis of promoters of known cancer genes show little excess of mutations beyond *TERT*.

## Introduction

Discovery of cancer drivers has traditionally focused on the identification of recurrently mutated protein-coding genes. Large-scale projects such as The Cancer Genome Atlas (TCGA) and the International Cancer Genome Consortium (ICGC) have profiled the genomes of a number of different cancer types to date, leading to the discovery of many putative cancer genes. Whole-genome sequencing (WGS) has made it possible to systematically survey non-coding regions for potential driver events. Recent studies of single-nucleotide variants (SNVs) and insertions/deletions (indels) detected from WGS data from smaller cohorts of patients have revealed putative candidates for regulatory driver events^1–8^. However, to date, only a few events have been functionally validated as regulatory drivers, affecting expression of one or more target genes by changing their regulation, RNA stability or disruption of normal genome topology^6,9–12^. Non-coding RNAs play diverse regulatory roles and are enzymatically involved in key steps of transcription and protein synthesis. Though functional evidence for non-coding RNAs in cancer is accumulating^13, 14^, only few examples have been shown to be the target of recurrent mutation^15, 16^. In contrast to protein-coding regions, the lower coverage and complexity of DNA sequence in these noncoding regions pose additional challenges for high-quality mutation calling, mutation rate estimation and identification of driver events. Adequate statistical methods that address these issues are needed to identify non-coding drivers from these data.

The ICGC and TCGA Pan-Cancer Analysis of Whole Genomes (PCAWG) effort, which has collected and systematically analyzed >2,700 cancer genome sequences from >2,500 patients representing a variety of cancer types^17^, offers an unprecedented opportunity to perform a comprehensive analysis of putative coding and non-coding driver events. Here, we describe results from multiple methods for detecting such drivers based on somatic SNVs and indels and a framework for integrating them. Using this approach, we identify significantly mutated genomic elements of various types in individual cancer types and across cancer types. We then use additional sources of information to remove from the initial list of significant elements those likely to result from mapping issues caused by repetitive regions or by inaccurate estimation of the local background mutation rate, and classify recurrently mutated elements as putative cancer-drivers or as likely false positives. Finally, we estimate our power to discover novel drivers in the PCAWG data set and evaluate the overall density of non-coding regulatory driver mutations around known cancer genes.

### Overview of recurrent mutations across cancer types

Many protein-coding driver mutations and the two well-studied *TERT* promoter mutations occur in frequently mutated single base genomic “hotspots”. In order to obtain an initial view of these in the PCAWG data set, we generated a genome-wide list of highly mutated hotspots by ranking all sites in the genome based on the number of cancers that harbor somatic mutations in them. Only 12 genomic sites were recurrently somatically mutated in more than 1% (26) of cancers and 106 in more than 0.5%, while the vast majority (93.19%) of somatic mutations were private to a single patient’s cancer (**Methods; Extended Data Fig. 1**). Interestingly, although protein-coding regions span only ~1% of the genome, 15 (30%) of the 50 most frequently mutated sites were known amino acid altering hotspots in known cancer genes (in *KRAS, BRAF, PIK3CA, TP53,* and *IDH1)* (**Fig. 1a**). Also among the top hits were the two *TERT* promoter hotspots. The top 50 list further contained a high proportion of mutations almost exclusively contributed by lymphoid malignancies (Lymph-BNHL and Lymph-CLL) (18 hotspots) and melanomas (6). The melanoma-specific hotspots were located in regions occupied by transcription factors (5/6 in promoters), a phenomenon that has recently been attributed to reduced nucleotide excision repair (NER) of UV-induced DNA damage^17–19^. Indeed, signature analysis of the melanoma hotspot mutations attributed them to UV radiation (**Fig. 1b**). The hotspots in lymphoid malignancies reside in the *IGH* locus on chromosome 14, a known phenomenon in B-cell-derived cancers where *IGH* has undergone somatic hypermutation by the activation-induced cytosine deaminase (AID), an observation confirmed by mutational signature analysis (**Fig. 1b**). Hotspots in an intron of *GPR126* and in the promoter of *PLEKHS1*^3, 20^ were located inside palindromic DNA that may fold into hairpin structures and expose single-stranded DNA loops to APOBEC enzymes^3^; accordingly, these mutations had high APOBEC signature probabilities. The six remaining non-coding hotspots could be attributed to a combination of mutational processes and presumed technical artifacts (**Supplementary Note**). In contrast to these non-coding and cancer-specific sites, mutations in protein-coding hotspots were attributed to the aging signatures, or to a mixture of signatures. Our observation that, with the exception of the *TERT* promoter hotspots, all of the top non-coding hotspots are generated by highly localized, cancer-specific mutational processes suggests that these may be passengers.

**Figure 1:**
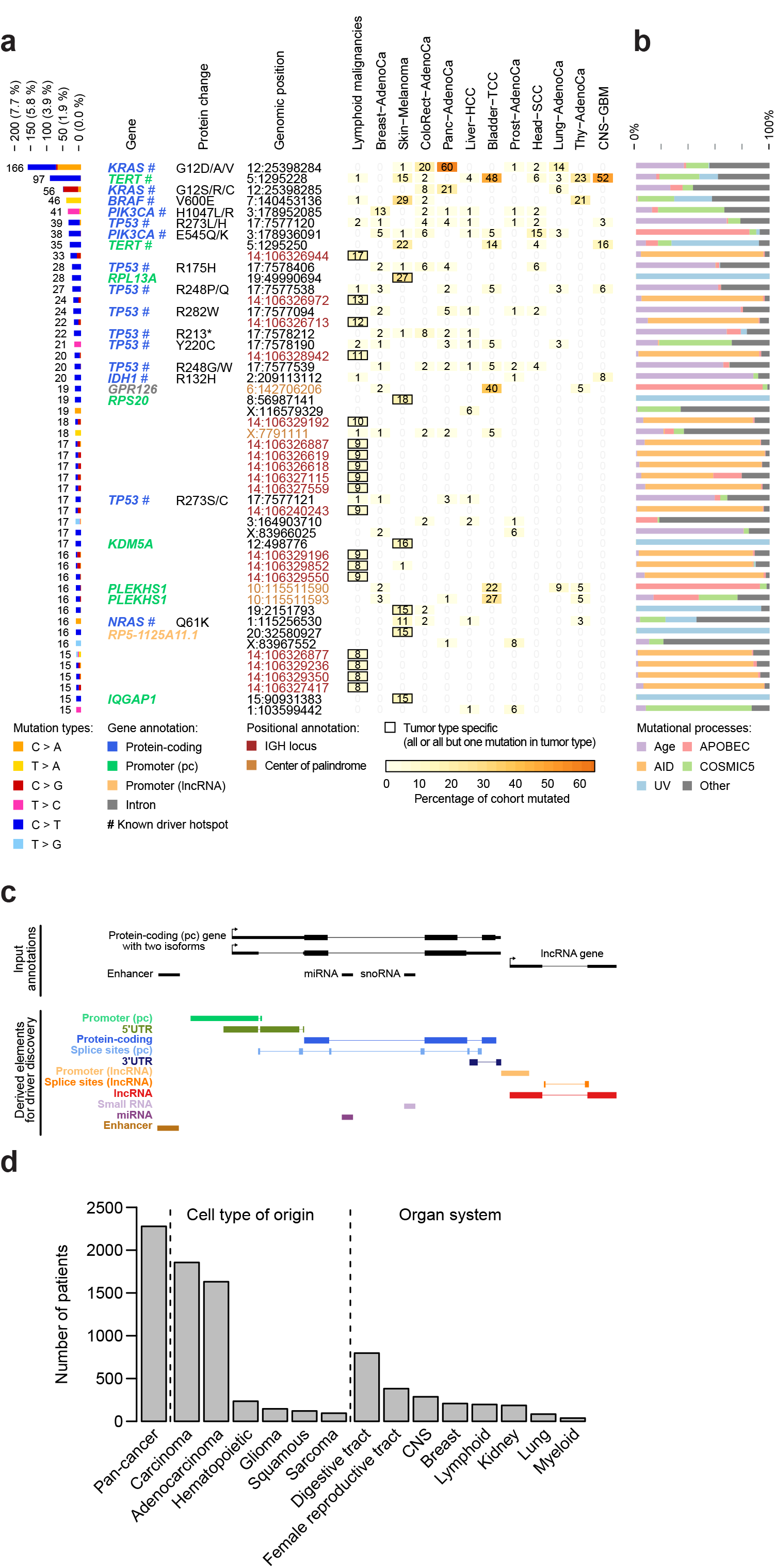
Mutational hotspots and overview of functional elements and meta-cohorts. **a**, Characteristics of top 50 single-site hotspots in PCAWG SNV data. The stacked bar chart (left) shows the total number of patients mutated across the PCAWG cohorts colored by mutation type. Gene names are given when hotspots overlap functional elements (colour-coded). Known somatic driver sites are marked with a hashtag. Amino acid change is indicated for protein-coding genes. The genomic location (chr:position) is colored by overlap with center of palindromes (orange) or immunoglobulin loci (brown). The table (right) shows the distribution of hotspot SNVs in the PCAWG cohorts sorted by number of hotspots. Tumor-type specific hotspots are indicated with a box. This only includes cohorts with at least 20 patients, and at least 10 patients or 10% of patients with a SNV. The Lymph-BNHL and Lymph-CLL cohorts are shown together as Lymphoid malignancies. See **Extended Data Fig. 1** for a complete set of individual and meta cohorts. **b**, Mutational signature attributions for mutations in each hotspot site. **c**, Schematic describing definition of functional element types from GENCODE and ENCODE annotation resources. Functional elements (black) are defined based on transcript annotations from various databases. Elements arising from multiple transcripts with the same gene ID are collapsed, as seen here for the protein-coding isoforms. Promoter elements are defined as 200 bases upstream and downstream of the transcription start sites of a gene’s transcripts (marked with red). Splice site elements extend 6 and 20 bases from 3’ and 5’ exonic ends into intronic regions respectively, as indicated. Regions overlapping protein-coding bases and protein-coding splice sites are subtracted from other regions, as indicated for the lncRNA promoter element in gray. **d**, Organization of meta-cohorts defined by tissue of origin and organ system (Methods). Pan-cancer contains all cancers excluding Skin-Melanoma and lymphoid malignancies.

Next, we sought to systematically identify genomic elements that drive cancer by comprehensively analyzing SNVs and indels in transcribed and regulatory genomic elements, including protein-coding genes, promoters, 5’UTRs, 3’UTRs, splice sites, distal regulatory elements/enhancers, lncRNA genes, short ncRNAs, and miRNAs (in total ~4% of the genome) (**Methods; Fig. 1c; Extended Data Fig. 2; Supplementary Table 1**). Drivers can be common across many cancer types (such as *TP53, PIK3CA, PTEN,* and *TERT promoter*) or highly specific (e.g. *CIC* and *FUBP1* in oligodendroglioma, *BCR-ABL* in chronic myelogenous leukemia, and *FOXA1* in breast and prostate cancers). Analysis of large mixed tumor cohorts increases the power to discover common drivers but dilutes the signal from cancer-specific ones. In order to maximize our ability to detect common and cancer-specific drivers, we performed analyses of individual tumor types, tumors grouped by their tissue of origin or organ site (“meta-cohorts”), as well as a pan-cancer set (**Fig. 1d; Methods**). This aggregation of tumors by tissue or organ site further allowed us to take advantage of patients from cohorts too small to analyze individually (<20 patients). Overall, we analyzed 2,583 unique patient samples in 27 individual tumor types and 15 meta-cohorts.

### Discovery of driver elements

In order to identify *bona fide* drivers in each cohort, we collected and integrated results from multiple driver discovery methods. These methods identify statistically significant elements based on SNVs and indels and evidence from one or more of three criteria: (i) mutational burden, (ii) functional impact of mutation changes, and (iii) clustering of mutation sites in hotspots (**Fig. 2a; Methods; Supplementary Table 2**). Generally, mutational burden and clustering of events are independent of the type of genomic element that is tested for significance. The ability to use measures of functional impact, however, may vary greatly depending on the element type. In protein-coding genes, functional impact is assessed through the amino acid change introduced by a given mutation. For some non-coding elements, the impact of mutations on transcription factor binding sites, miRNA binding sites, or functional RNA structure may be predicted. However, in general, the exact consequences of somatic mutations are harder to assess in non-coding regions than in protein-coding regions, and may even vary substantially across cell types.

**Figure 2:**
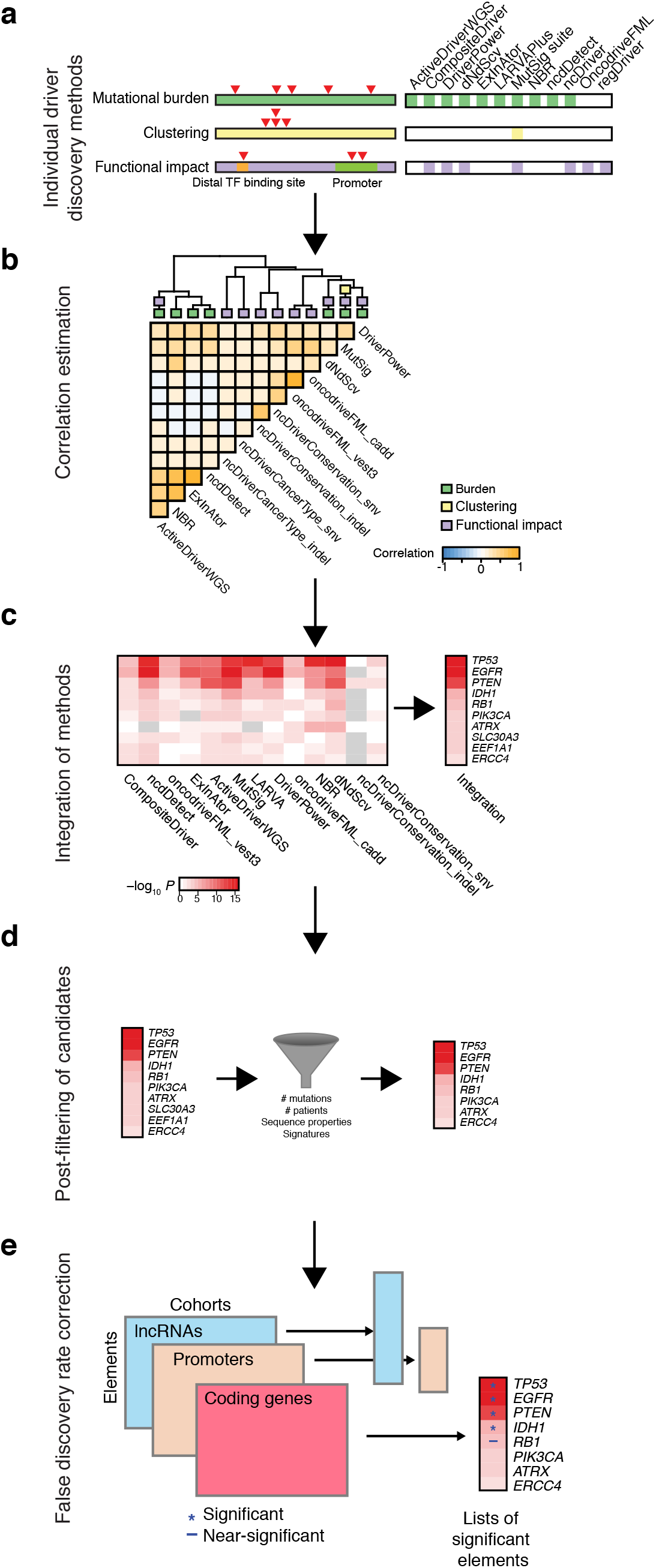
Filtering and integration process for nomination of driver candidates. **a**, Overview of driver discovery method and their lines of evidence to evaluate candidate gene drivers. Methods employing each feature are marked with blue boxes next to the appropriate track. **b**, Spearman’s correlation of p-values across the different driver discovery algorithms based on simulated (null model) mutational data. Dendrogram illustrates relatedness of method p-values and algorithm approaches marked by colored boxes on dendrogram leaves. **c**, P-values are combined with Brown’s method based on the correlation structure calculated in (b). Individual method (left) and integrated (right) log-transformed p-values are shown in a heatmap (grey: missing data). **d**, Post-filtering used several criteria to identify likely suspicious candidates (shaded grey rectangle). **e**, Significant driver candidates were identified after controlling for multiple hypothesis testing based on an FDR q-value threshold of 0.1 (blue asterisk). Candidates with q-values below 0.25 (blue dash) were also considered of interest.

### Integration of results from multiple driver discovery methods

Most cancer genomic studies to date have employed a single method to identify putative cancer driver elements. This can introduce biases because the number and type of drivers detected by individual methods depends on their underlying assumptions and statistical model. To reach a comprehensive consensus list of candidate drivers, we collected the p-values calculated by up to 16 driver discovery methods for each genomic element (**Supplementary Table 2, 3**) and derived a general strategy to integrate them (**Fig. 2a**). Our strategy accounts for the correlation of p-values among driver discovery methods that use similar approaches (burden, clustering, functional impact) when applied to the same data, while accumulating the independent evidence provided by different methods (**Fig. 2b**). To integrate potentially correlated statistical results, we used Brown’s method for combining p-values^21^, an extension of the popular Fisher’s method^22^ (**Fig. 2c**). We first used simulated data sets that preserve local mutation rates and signatures but do not contain drivers, to assess the specificity of each method and the correlation structure among different methods when no genomic element is expected to have a significant excess or pattern of mutations (**Methods; Extended Data Fig. 3**). As anticipated, p-values were correlated among methods that share similar approaches. We then used the resulting correlation structure to integrate p-values from the observed (real) mutation data from different methods into a single p-value for each genomic element. Since simulated data sets are based on assumptions, we tested whether we can directly estimate the correlation structure from p-values calculated on the observed data. As this simpler approach yielded analogous results, it was used throughout this study (**Methods, Extended Data Fig. 3**). After additional filtering steps (see below), we conservatively controlled for multiple hypothesis testing using the Benjamini-Hochberg procedure (BH)^23^ across the 27 individual and 15 meta-cohorts, for each element type separately (**Fig. 2e**). Cohort-element combinations (hits) with q<0.1 (10% false discovery rate [FDR]) were considered significant. Overall, the union of all hits from the individual driver discovery methods contained 3,048 cohort-element pairs involving 1,694 unique elements, whereas the integrated results contained 1,406 significant hits with 635 unique elements (**Supplementary Table 4, 5**).

### Flagging and removal of potential false positive hits

Even after careful variant calling, false positive identification of driver loci can arise from residual inaccuracies in background models, residual sequencing and mapping artifacts or from local increases in the burden of mutations generated by as-yet unmodeled mutational processes. To minimize technical issues, we carefully reviewed and filtered all candidate driver elements based on low-confidence mappability regions and site-specific noise in a panel of normal samples from PCAWG (**Fig. 2d; Fig. 3a,b; Methods**). For example, the lncRNA *RN7SK* is a small nuclear RNA with many homologous regions in the genome, making it prone to read mapping errors (**Fig. 3a**), while a fraction of *LEPROTL1* mutations fell in positions flagged by the site-specific noise filter (**Fig. 3b**).

**Figure 3:**
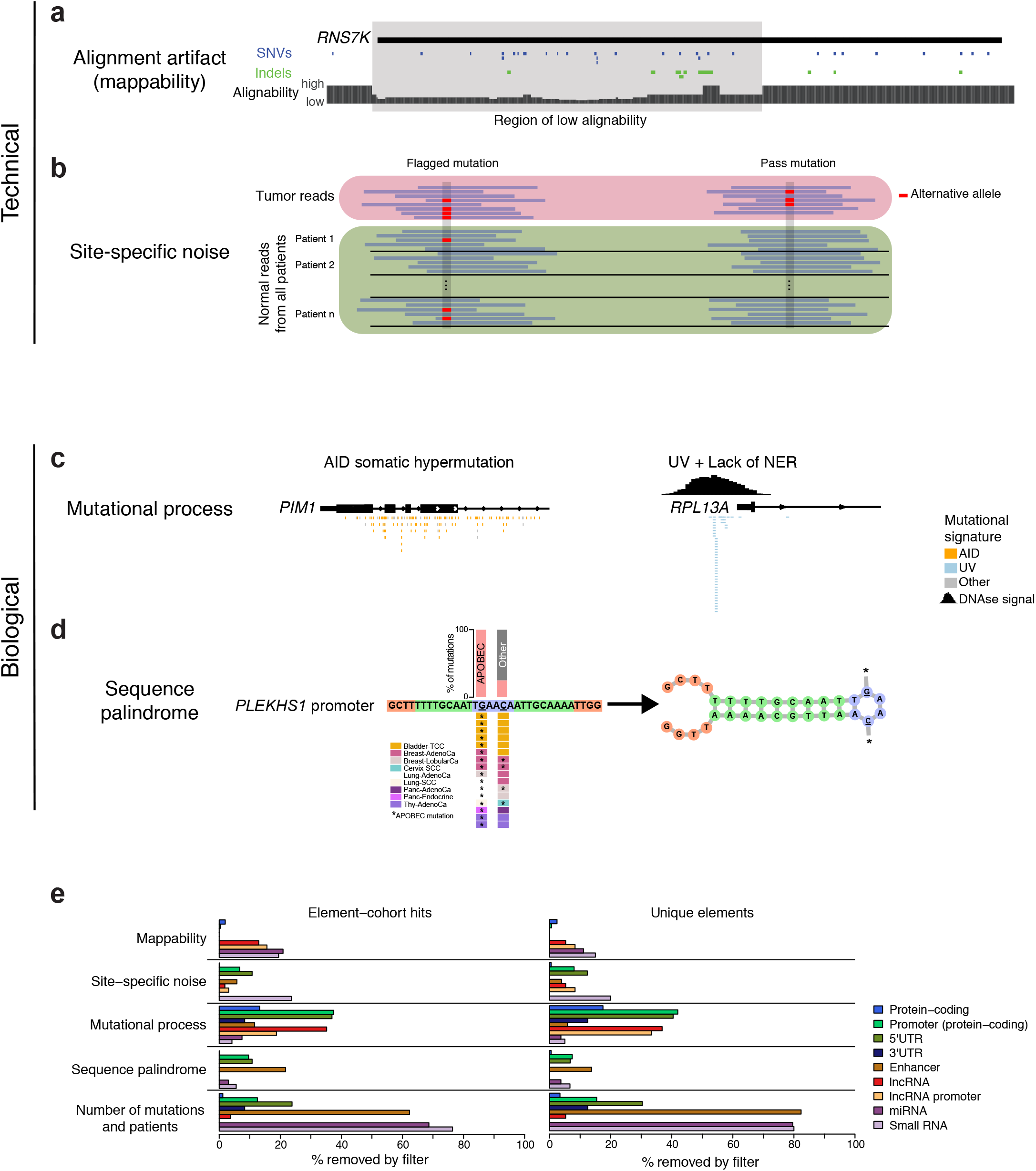
Technical and biological confounders used in candidate filtering. **a**, Read mappability based on alignability, uniqueness and blacklisted regions. Example shown for *RN7SK,* which was removed due to low alignability. **b**, Site-specific noise evaluated in normal samples from PCAWG. **c**, Increased local mutation density caused by AID and UV mutational processes in lymphoma and melanoma, respectively. **d**, Palindromic DNA can expose bases to APOBEC enzymes. **f**, Number of significant unique hits (left) and unique elements (right) removed by each filter.

In addition, mutational processes not properly captured by current background models can cause local accumulation of mutations. As shown in the hotspot analysis above, events in the promoters of *PIM1* and *RPL13A* are strongly associated with the AID and UV mutational processes, respectively, as previously noted^18, 19, 24^ (**Fig. 3c**). Moreover, DNA palindromes may form hairpin structures that expose loop bases to APOBEC enzymes, as has been proposed for *PLEKHS1, GPR126, TBC1D12* and *LEPROTL1^2, 3^* (**Fig. 3d**). Non-coding RNAs that form secondary structures often contain palindromic sequences, which could be preferential targets for APOBEC enzymes. However, only a small fraction of the mutations in structural RNAs appear to be APOBEC derived overall (4.8%) (**Supplementary Note**). In total, 569 out of 1,406 hits (element-cohort combinations with q<0.1) and 383 out of 635 unique elements that were initially significant, were filtered out, resulting in 837 hits (**Fig. 3e; Supplementary Tables 4, 5**). Finally, multiple hypothesis correction was repeated after assigning a non-significant p-value (*P* = 1) to the filtered out hits, resulting in further removal of 91 candidate element-cohort combinations.

### Performance of the integration method

To evaluate the sensitivity and validity of the integration approach, we compared the performance of individual methods and of the integration on protein-coding genes using, as a gold-standard, a list of 603 known recurrently-mutated cancer genes in the Cancer Gene Census^25^ (CGC v80). These analyses revealed that, typically, the integration of different methods outperformed individual methods, both in terms of sensitivity and specificity, yielding lists of significant genes that were longer and more enriched in high-confidence cancer genes (**Extended Data Fig. 4**).

### Discovery of recurrently mutated elements

Our conservative integration and filtering strategy yielded 746 total hits, which include 646 hits in 157 protein-coding genes, 30 hits in seven protein-coding promoters, 26 hits in eight long non-coding RNAs, 18 hits in six non-coding RNA promoters, ten hits in six 3’UTRs, six hits in four 5’UTRs, six hits in one enhancer, three hits in one microRNA and one hit in a small RNA candidate (**Fig 4a; Supplementary Table 5**). The number of significant elements varied from just one in clear-cell renal cancer to 78 in the Carcinoma meta-cohort (**Fig. 4b**) and depended strongly on cohort size (r = 0.84; *P* = 1.9x10^−8^; also see power analysis below), reflecting that the landscape of driver candidates, particularly in small tumor cohorts, is still incomplete.

**Figure 4:**
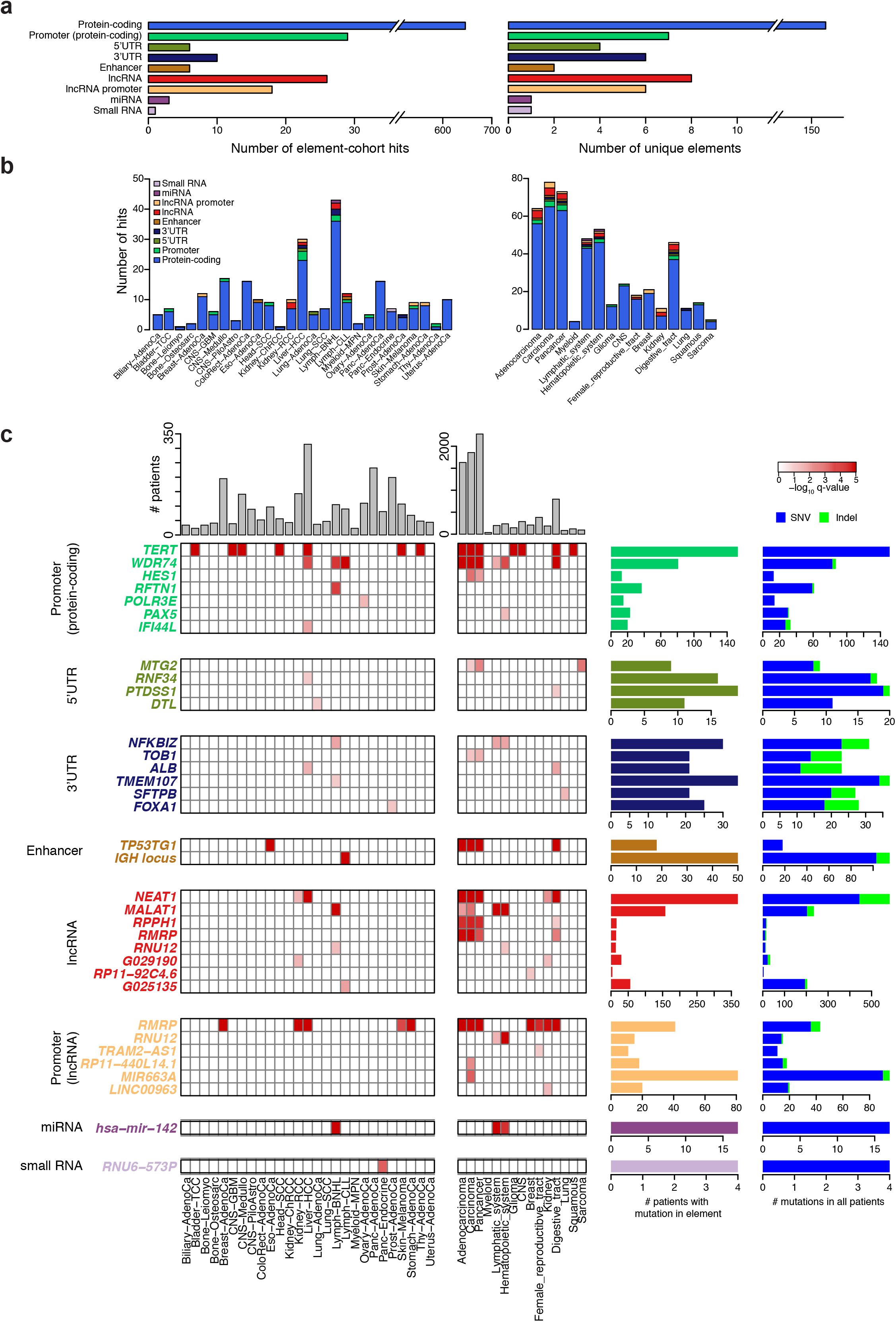
Protein-coding and non-coding driver elements identified in PCAWG cohorts. **a**, Number of total hits (elements significant in a certain patient cohort; left) and number of unique significant elements (right). **b**, Number of significant elements by type and by cohort. **c**, Significant non-coding elements identified in this study. Element types are indicated with colors as in **Fig. 1c**.

Comparison of the p-values obtained from individual tumor types and meta-cohorts revealed that although most candidate drivers gained significance in larger meta-cohorts, the tumor-type specific genes *DAXX* (Panc-Endocrine), *BRAF* (Skin-Melanoma), *NRAS* (Skin-Melanoma), *SPOP* (Prost-AdenoCa) and *hsa-mir-142* (Lymph-BNHL) scored higher in their respective tumor types (**Extended Data Fig. 5**). These results emphasize the need for careful tradeoff between tumor-type specificity and cohort size when searching for drivers.

#### Protein-coding mutations

The ability to discover known protein-coding cancer genes from WGS data provided the opportunity to evaluate our strategy and to put non-coding driver hits in context (**Extended Data Fig. 6**). Overall, candidate coding drivers were concordant with previous results: of the 157 genes significant in at least one patient cohort, 64% are listed in the CGC and 87% are in a more comprehensive list of cancer genes (https://doi.org/10.1101/190330). In contrast to prior studies based on large exome sequencing data sets, the moderate number of patients per cancer type in this data set provided sufficient power to detect only the genes with the strongest signal. Indeed, when we relaxed the significance threshold (q<0.25) to detect hits “near significance”, we found 93 additional hits in 62 unique genes. Half (31) of these genes were already discovered as significant (q<0.1) in at least one cohort in this study, and now became significant in additional cohorts. Of the 31 genes not previously identified by our analyses, 19% were in the CGC and 32% in the more comprehensive list of drivers. These results confirm that many cancer genes were just beyond our significance threshold and would likely have been discovered with larger cohorts (**Supplementary Table 4**).

### Non-coding driver candidates

In contrast to protein-coding elements, there was far less agreement among significance methods when analyzing non-coding elements, and most hits were strongly supported by only a few methods (**Extended Data Fig. 7**). The limited agreement between driver discovery methods in non-coding elements is likely due to a weaker signal of selection and different underlying assumptions that imperfectly model background mutation frequencies in non-coding regions. Thus, in order to nominate a significant element as a candidate driver, we carefully reviewed the supporting evidence from the genomic data and sought additional evidence from PCAWG (chromosomal breakpoints, copy-number, loss-of-heterozygosity and expression data), cancer gene databases and the literature (**Supplementary Table 6**).

#### Promoters and 5’UTRs

Due to the overlap between the downstream regions of promoters and 5’UTRs, we reviewed the 11 significant elements from our promoter and 5’UTR analyses together, yielding several interesting candidates. Our integrated results confirmed that the *TERT* promoter is the most significantly mutated promoter in cancer, being significant in 8 individual tumor types and 11 meta-cohorts (**Fig. 4c; Supplementary Table 5**). As previously reported, we observed significantly higher *TERT* transcript expression levels in mutated compared to non-mutated cases (**Extended Data Fig. 8**).

Among the remaining candidates, we found recurrent mutations in the promoters of *PAX5* and *RFTN1* in lymphoma. *PAX5* is a known off-target of AID hypermutation^24^ and, indeed, the percentage of its mutations attributable to AID activity (46%) was just below our filtering threshold. Consistently, mutations in its promoter were not associated with gene expression changes, suggesting that these mutations may be passengers (**Extended Data Fig. 8**). In contrast, mutations in the promoter of *RFTN1* were weakly associated with an increase in *RFTN1* expression levels (P = 0.03; mutant/wild type fold difference (FD) = 1.2; *P* = 0.02, after excluding 8/21 mutations attributed to the AID signature) (**Fig. 5a**). The protein encoded by *RFTN1,* Raftlin, associates with B-cell receptor complexes^26^ but its function in lymphoma is unclear.

**Figure 5:**
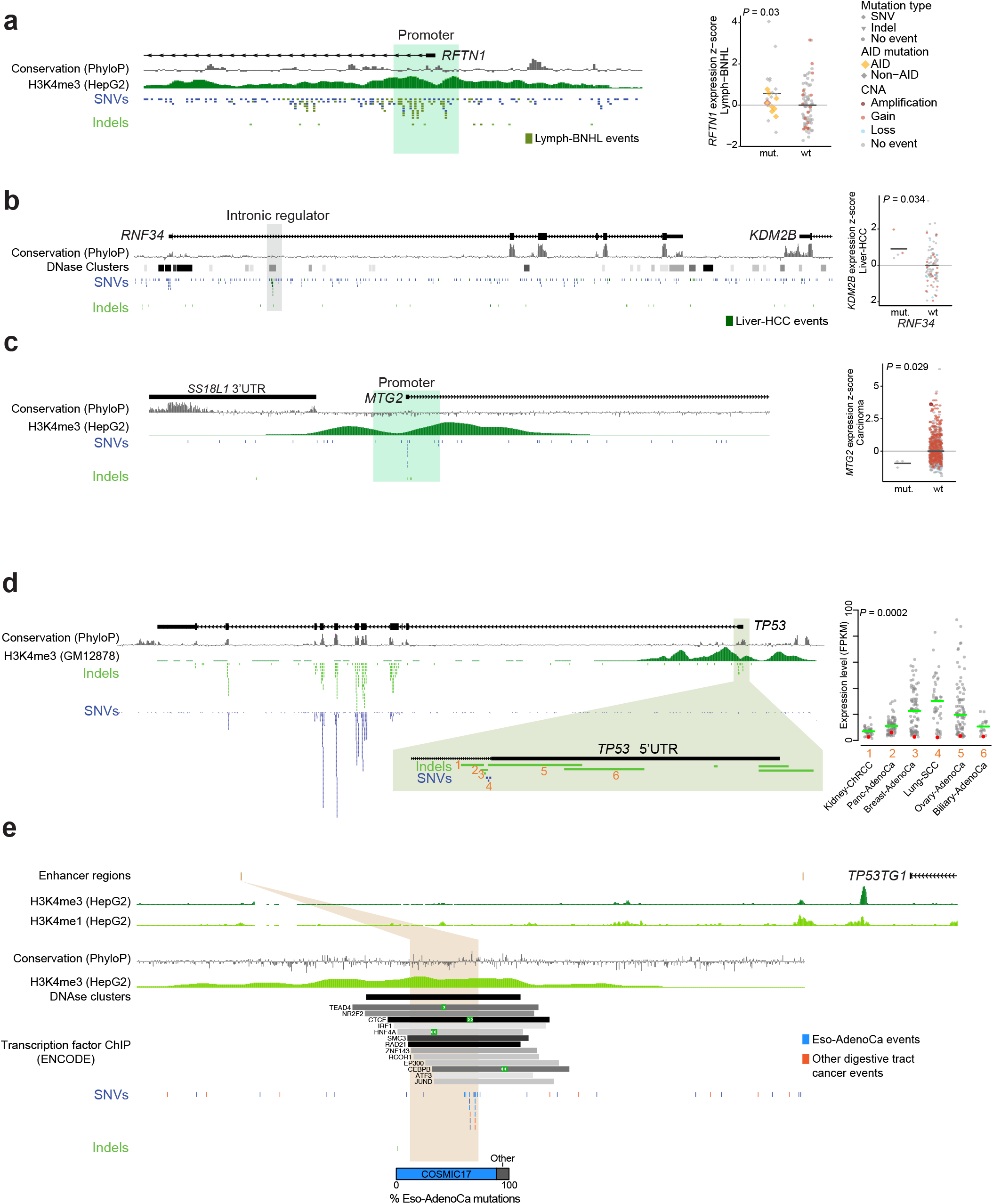
Novel non-coding promoter and enhancer driver candidates. **a**, *RFTN1* promoter locus (left) and gene expression for mutant (mut.) and wild-type (wt) (right) in Lymph-BNHL tumors. Expression values are colored by copy number status; AID mutations are highlighted in yellow. **b**, A mutational hotspot inside *RNF34* overlapping a DNAse I site (left) correlates with gene expression changes of *KDM2B* in Liver-HCC (right). Coloring of mutations as in a. **c**, *MTG2* promoter locus (left) and associated gene expression changes in Carcinoma tumors (right). Coloring of mutations as in a. **d**, An enhancer associated with *TP53TG1* (Methods) (left) contains mutations mostly attributed to an esophageal cancer-associated signature.

Mutations called in the 5’UTR of *RNF34* in Liver-HCC overlap an intronic DNAse I hypersensitive region in multiple cell types^27^, suggesting that events may affect expression of *RNF34* or a neighboring gene through an intragenic enhancer. Indeed, expression of the histone demethylase *KDM2B* located downstream of *RNF34* shows weak correlation with these mutations (P = 0.03; FD = 1.3; **Fig 5b**) with no effect on *RNF34* mRNA (P = 0.41). Mutations in the 5’ region of *MTG2* were concentrated in a hotspot and showed marginally significant decreased expression in the Pan-cancer (P = 0.036; FD = 0.8) and Carcinoma (P = 0.029; FD = 0.8) meta-cohorts (**Fig. 5c; Extended Data Fig. 8**). *MTG2* encodes a little-studied GTPase that associates with the mitochondrial ribosome^28^. *HES1* promoter mutations were significant in Carcinoma and Pan-cancer, but showed no association with gene expression (**Extended Data Fig. 8**). *HES1* is a NOTCH signalling target^29^, and is focally amplified in gastric cancers (**Extended Data Fig. 9**).

*PTDSS1, DTL* (and *INTS7,* which shares a bidirectional promoter), *IFI44L* and *POLR3E* showed trends towards increased or decreased expression in mutated tumors, although the small number of samples prevented us from drawing definitive conclusions (**Extended Data Fig. 8**). None of these genes were present in significantly amplified or deleted focal peaks (**Methods**). Validation of these hits with additional data in further studies will be needed to evaluate whether they are genuine drivers.

As previously reported in a number of studies^1, 3, 7^, we also found recurrent mutations in the promoter of *WDR74,* which was significant in multiple cohorts. The mutations concentrate in a small region overlapping an evolutionarily conserved spliceosomal U2 snRNA and therefore potentially affect splice patterns. However, no significant associations with either *WDR74* or transcriptome-wide splicing were seen (**Supplementary Note; Methods**). We did, however, find that the repetitive U2 sequence was frequently affected by recurrent artifacts in SNP databases, raising concerns about potential mapping errors for this repetitive RNA type (**Extended Data Fig. 10**).

Finally, restricted hypothesis testing of the promoters of CGC genes (n = 603) revealed significant recurrence of mutations in the promoter region of *TP53* (11 patients in the pancancer cohort; q = 0.044, NBR method). Most of these mutations were substitutions and deletions affecting either the TSS or the donor splice site of the first non-coding exon of *TP53.* For the 6 samples with expression data, these mutations were associated with a dramatic reduction of expression (**Fig. 5d**). In 8 of the 11 mutant cases, the mutation occurred in combination with copy-neutral loss of heterozygosity. This is the first report of a relatively infrequent, but impactful, form of *TP53* inactivation by non-coding mutations.

#### Enhancers

Two enhancer regions were found significant in our integration analysis; one at the *IGH* locus in Lymph-CLL and the other near *TP53TG1* in multiple cohorts. However, many of the recurrent mutations in the enhancer overlapping the *IGH* locus are due to characteristic somatic hypermutation; 48% of mutations in this enhancer element matched the AID signature, which is just below our filtering threshold. Nine of 18 mutations in the enhancer near *TP53TG1* were contributed by esophageal cancers. However, 89% of mutations in these esophageal tumors were attributed to a mutational signature common in this cancer type (mostly T>G substitutions in NTT context), raising the concern that these reflect a localized mutational process rather than a driver event^30^ (PCAWG7 publication). These mutations were concentrated around a conserved region (Fig. 5e) and overlapped sites bound by NFIC and ZBTB7 transcription factors in HepG2 cells^27^. *TP53TG1* is a noncoding RNA suggested as a tumor suppressor involved in p53 response to DNA damage and has been reported to be epigenetically silenced in cancer^31^.

#### 3’UTRs

Recurrent somatic events were identified in the 3’UTRs of six genes: *FOXA1* in prostate cancer, *ALB* in liver cancer, *SFTPB* in lung adenocarcinoma, *NFKBIZ* and *TMEM107* in lymphomas, and *TOB1* in Carcinoma, most of which contained a large number of indels (**Fig. 4c**). High rates of indels in the 3’UTR of *ALB* in liver cancer and *SFTPB* in lung cancer have been recently reported^8^ but it is unclear whether they are cancer drivers or the result of a poorly-understood mechanism of localized indel hypermutation. To determine whether indels affect function and accumulate specifically in the 3’UTRs, we compared indel rates in 3’UTRs with other regions around these genes. Consistent with local hypermutation, rather than selection, we observed similar indel rates in 3’UTR and 1 kb downstream of the polyadenylation site in *ALB, SFTPB* and *FOXA1* (**Fig. 6a**). This, together with additional observations presented below, strongly suggest that indel recurrence at these loci is caused by a novel localized hypermutation process.

**Figure 6:**
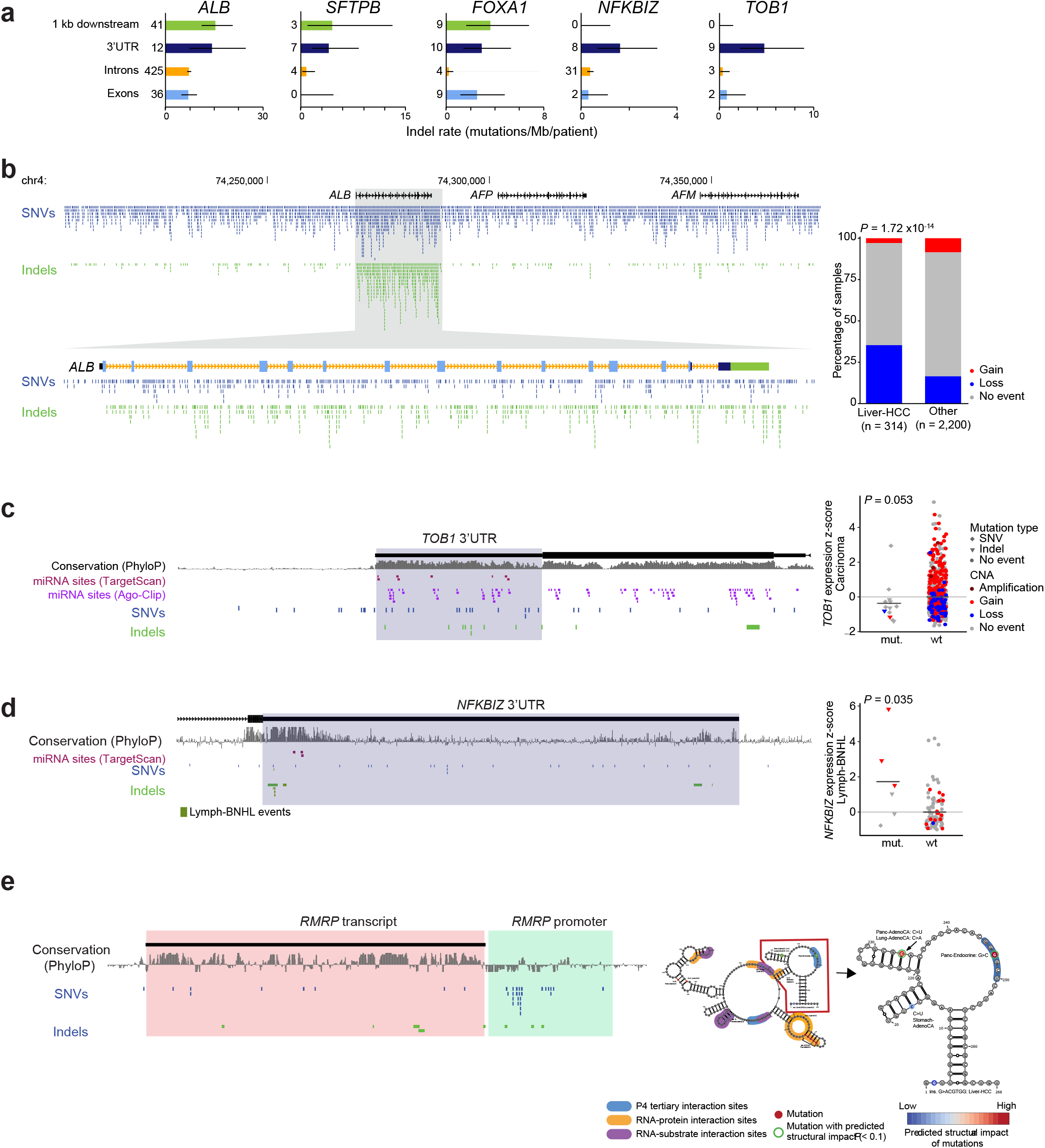
Novel 3’UTR and non-coding RNA driver candidates. **a**, Quantification of indel rates for functional elements associated with significant 3’UTRs. **b**, Overview of *ALB* genomic locus and PCAWG variants illustrates local density of indels. Coloring of enlarged *ALB* gene elements corresponds to functional elements in barplots in a. Copy number changes from 314 Liver-HCC cases are compared to other samples from PCAWG. **c**, Genomic locus of *TOB1* 3’UTR. Note extreme conservation (PhyloP) for the 3’UTR. **d**, Genomic locus of *NFKBIZ* 3’UTR (left) and associated gene expression changes in Lymph-BNHL (right). **e**, Genomic locus of the *RMRP* transcript and promoter region (left), and its RNA secondary structure, tertiary structure interactions, protein and substrate interactions, and mutations with their predicted structural impact (right; **Extended Data Figure 9d**); lymphoma and melanoma are excluded.

On the other hand, although *ALB* is subject to localized hypermutation, protein-coding SNVs in *ALB* are enriched in missense and truncating events relative to synonymous mutations in liver cancer (*P* = 1.5x10^−7^; **Fig. 6b**). Furthermore, the *ALB* locus is lost in a fraction of liver cases (**Fig. 6b**), with copy losses having a tendency to occur in samples without somatic mutations (Fisher’s exact test, *P* = 0.0099). These findings, suggest that loss of *ALB* may be a genuine driver event in liver cancer. Similarly, the well-known driver role of *FOXA1* in prostate tumors is supported by the functional impact of coding mutations (*P* = 4.4x10^−7^) and focal amplifications, raising the possibility that the mutation enrichment observed in the 3’UTR might also be functional.

In contrast to the genes discussed above, the number of indels in the 3’UTRs of *NFKBIZ* and *TOB1* was significantly higher than in other parts of these genes, suggesting that indels in the 3’UTRs could be under functional selection (**Fig. 6a**). *TOB1* encodes an anti-proliferation regulator that associates with ERBB2, and also affects migration and invasion in gastric cancer^32^. As part of the CCR4-NOT complex, TOB1 regulates other mRNAs through binding to their 3’UTR and promoting deadenylation^33^. Tumors with 3’UTR mutations in *TOB1* showed a trend towards decreased expression (P = 0.053; FD = 0.7). Mutations did not concentrate in known miRNA binding sites, possibly due to incomplete annotation (**Fig. 6c**). However, the extreme conservation of this region (5th most conserved among all tested 3’UTRs with an average PhyloP score of 5.6) indicates that these events likely disrupt functionally important sites (**Fig. 6c**). Interestingly, *TOB1* and its neighboring gene *WFIKKN2* are focally amplified in breast cancer and pan-cancer, suggesting a complex role in cancer (**Extended Data Fig. 9**). *NFKBIZ* is a transcription factor that is mutated in recurrent/relapsed DLBCL^34^ and amplified in primary lymphomas^34, 35^. 3’UTR mutations accumulated in a hotspot proximal to the stop codon and upstream of conserved miRNA binding sites which might be affected by these mutations (**Fig. 6d**). Only two events in these regions are SNVs, suggesting that mutations in the *NFKBIZ* 3’UTR are not the consequence of AID off-target activity. Lymphomas with 3’UTR mutations showed a trend towards increased mRNA levels of *NFKBIZ (P* = 0.035; FD = 3.2; the trend remains after correction for copy number, *P* = 0.03; **Fig. 6d**). Functional studies will be required to understand the exact function of *3’UTR* mutations on transcript regulation of *TOB1* and *NFKBIZ.*

Mutations in the significant 3’UTR of *TMEM107* in lymphoma overlap the U8 small RNA *SNORD118* (**Extended Data Fig. 10**). Similar to *WDR74*, overlap with a repetitive RNA element suggests that these mutations might be caused by mapping artifacts.

#### Non-coding RNAs

ncRNAs are often part of large gene families with multiple copies in the genome. Sequence similarity with other genomic regions may therefore complicate their analysis. Similarly to the *U2* RNA upstream of *WDR74,* we observed high levels of germline polymorphisms in normal samples in some of the significant ncRNAs and their promoters (**Extended Data Fig. 10**). Though interesting candidates, caution should thus be exercised in interpreting these hits.

The non-coding RNA *RMRP* is significantly mutated in multiple cancer types, in both its gene body and promoter (**Fig 4c; Fig 6e; Supplementary Table 5**). *RMRP* is the RNA component of the endoribonuclease RNase MRP, an enzymatically active ribonucleoprotein^36, 37, 38^. Its catalytic function depends on the RNA secondary and tertiary structure and its interactions with proteins^39^. Germline mutations in *RMRP* cause cartilage-hair hypoplasia, and somatic promoter mutations have been reported to be functional^9^. In addition, the *RMRP* locus is focally amplified in several tumor types, including epithelial cancer (**Extended Data Fig. 9c**). SNVs in the gene body (7 in pan-cancer) are significantly biased towards secondary structure impact (*P* = 0.011, permutation test)^40, 41^. Three of these are individually significant (each with *P* < 0.1, sample level permutation tests), with two affecting the same position and one located in a tertiary interaction site (**Fig. 6e**). Of the four gene-body indels, three are located in or near protein-binding sites (*P* = *0.08*), including a deletion that is predicted to affect the secondary structure (**Extended Data Fig. 9d**). Given *RMRPs* role in replication of the mitochodrial genome^36^, we tested whether mutations in this locus were associated with altered mitochondrial genome copy number. Indeed, mutated samples showed a trend towards higher mitochondrial copy number (two-sided rank-sum test, *P* = 0.1). However, *RMRP* also appears to be a target of potential artifacts (**Extended Data Fig. 10**) and so its relevance to cancer requires further scrutiny.

The microRNA precursor (pre-miRNA) for *miR-142* was found significant in Lymph-BNHL, Lymphatic system and Hematopoietic system (**Fig. 4c; Supplementary Table 5**). The locus is a known AID off-target in lymphoma^5, 42^(**Extended Data Fig. 10**). However, five of seven mutations (71%) in the dominantly expressed mature miRNA *mir-142-p3*, where the largest functional impact is expected, were not assigned to AID, raising the possibility that they may be under selection^5^.

Eleven additional ncRNA-related elements *(RNU6-573P, RPPH1, RNU12, TRAM2-AS1, G025135, G029190,RP11-92C4.6* and promoters of *RNU12, MIR663A, RP11-440L14.1*, and *LINC00963*) passed our stringent post-filtering, but had limited supporting evidence after manual inspection. These are discussed in a Supplementary Note. We further checked whether any of our non-coding candidates were associated with known germline cancer risk variants, and found no significant associations, with the exception of *TERT* (**Methods**).

*NEAT1* and *MALAT1,* two neighboring lncRNAs previously reported as recurrently mutated in liver^43^ and breast cancer^3^, were found significant in multiple cohorts (**Fig. 4c**), mutated in a large number of patients (369 for *NEAT1* and 158 for *MALAT1;* **Fig 7a**). In addition, these genes are significantly associated with nearby chromosomal breakpoints *(NEAT1: P* = 5.3x10^−9^; *MALAT1: P* = 0.0012; **Methods; Extended Data Fig. 11**). The pattern of breakpoints, indels and SNVs scattered throughout their sequence would suggest a possible tumor suppressor role. To evaluate this hypothesis, we tested whether *MALAT1* and *NEAT1* mutations are associated to loss of heterozygosity (LOH). Neither gene showed biallelic loss, in contrast to many known canonical tumor suppressors (**Fig. 7b; Extended Data Fig. 11**). Mutations in *MALAT1* and *NEAT1* also do not exhibit higher cancer allele fractions compared to mutations in flanking regions, a feature of early driver mutations (**Fig 7c; Extended Data Fig. 11**). Furthermore, *NEAT1* and *MALAT1* mutations are not associated with altered expression levels, suggesting that they do not affect post-transcriptional stability nor show increased expression like many mutated oncogenes (**Fig 7d; Extended Data Fig. 11**).

**Figure 7:**
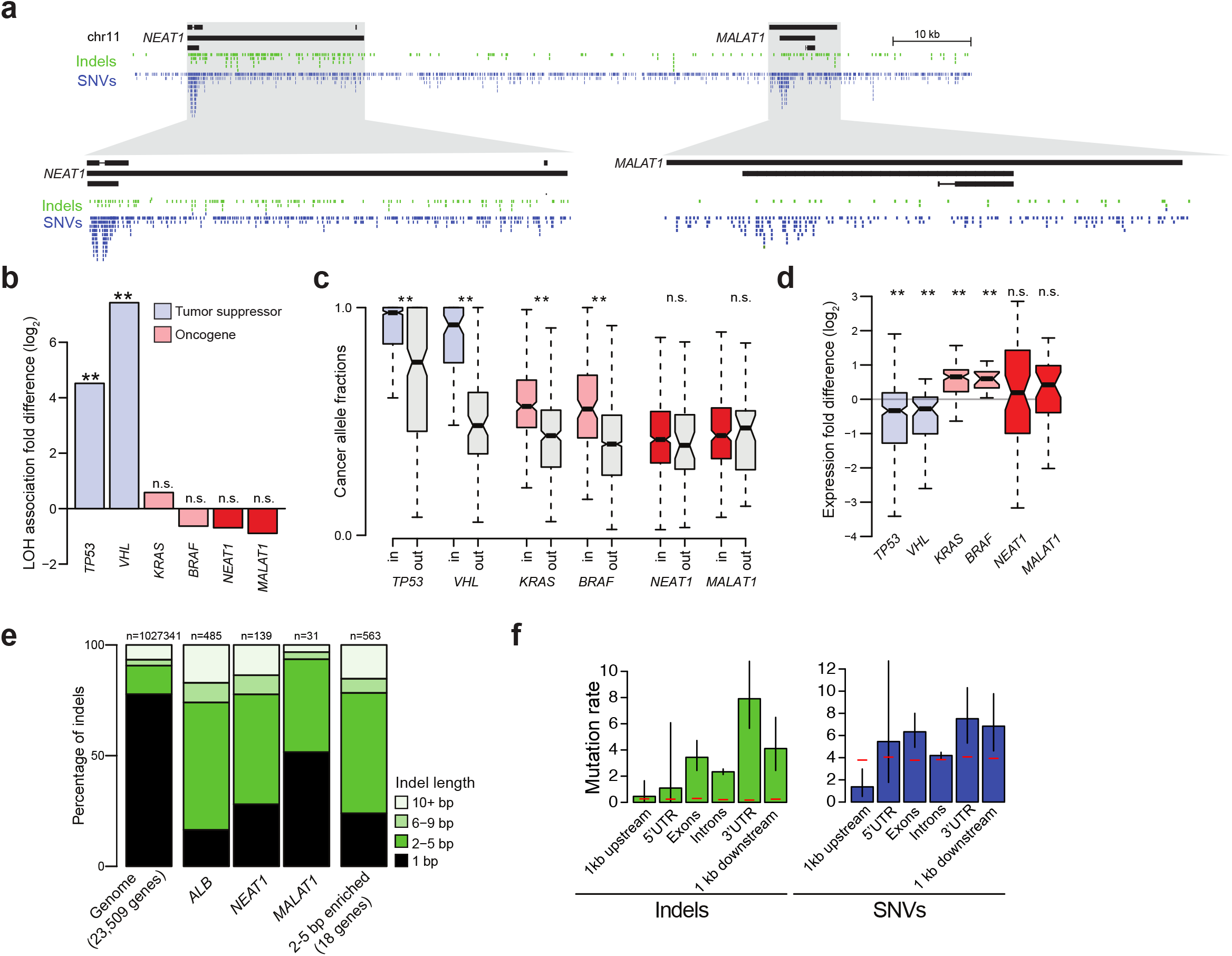
Characterization of a mutational process in *NEAT1* and *MALAT1*. **a**, Genomic locus overview showing the distribution of indels and SNVs in *NEAT1* and *MALAT1.* **b**, Analysis of enrichment in LOH associated with mutation (“double-hit”) for canonical tumor suppressors (TSG), oncogenes (OG), and *NEAT1/MALAT1*. **c**, Comparison of cancer allelic fractions within those same genes and within flanking regions (including 2 kbp upstream, 2 kbp downstream and introns). **d**, Difference in expression between mutated and wild-type alleles of these genes. **e**, Percentages of different groups of indel sizes for all protein-coding and lncRNA genes, *ALB, NEAT1, MALAT1* and the set of genes enriched in 2-5 bp indels. **f**, SNV and indel rates (total events/bp) in different functional regions of 18 protein-coding genes enriched in 2-5 bps indels (without *ALB,* which contributed 47% of indels). Red lines indicate background indel and SNV rates estimated from all protein-coding genes.

The high enrichment of indels throughout the gene body of *NEAT1* and *MALAT1,* which have very high expression levels across many cancer types, resembles the phenomenon described above for *ALB* and *SFTPB.* If these indels are generated by a specific mutational process, we might expect distinct features from indels found elsewhere in the genome. Indeed, we find that indels in *NEAT1, MALAT1, ALB* and *SFTPB* are strongly enriched in events longer than 1 bp and particularly in indels of length 2-5 bp, compared to the genomic background (least significant Fisher’s *P* = 6.8x10^−5^, for *MALAT1;* **Fig. 7e**). The association of these indels with highly expressed genes suggests a transcription-coupled mutagenic mechanism, possibly transcription-replication collisions^44–46^. A systematic search of genes with increased rates of 2-5 bp indels reveals that this yet-unknown mutational process affects other highly expressed genes (**Extended Data Fig. 11**), some of which were reported in a recent study^8^, indicating that these indel sizes are a feature of the reported mutational process. Interestingly, SNVs also occur at higher frequencies in these genes, suggesting that they may be generated by the same mechanism, although less frequently (**Fig. 7f**).

Overall, the discovery of a localized mutational signature and the lack of association of the mutations in *MALAT1* and *NEAT1* with loss of heterozygosity or higher allele frequencies suggests that indels in these genes are most likely passenger events. The previously reported oncogenic phenotypes associated with both ncRNAs^43, 47, 48^ may thus be related to other types of alterations.

### Localized lack of power to call mutations

Certain genomic regions, especially those with high GC content, are subject to systematic low coverage in next-generation sequencing^49^. We used power analysis to quantify the number of possibly missed mutations, especially in non-coding regions. These calculations attempt to account for variations in local sequencing depth, purity and overall ploidy of individual tumor samples, background mutation rates across cohorts and elements, and the size of patient cohorts^9,50–52^. We found that mutation detection sensitivity was high across the element types, (median d.s. = 0.98) but slightly lower for the typically GC-rich promoters (median d.s. = 0.96) and 5’UTR elements (0.96; **Fig. 8a; Supplementary Table 4,5**). However, for some individual genomic regions the detection sensitivity is dramatically reduced and it is therefore important to evaluate it for elements of interest. For example, only 43% of tested cases have at least 50% average detection sensitivity across the third exon of *TCF7L2* and 20% in the 5’UTR of *AKT1* (**Extended Data Fig. 12a**). Positional detection sensitivities for the two canonical *TERT* promoter hotspot sites were highly variable among patients and cohorts, ranging from 4% of sufficiently powered (≥90%) patients in CNS-PiloAstro to 100% of patients in Thy-AdenoCa (**Fig. 8b; Extended Data Fig. 12b**). This means that we have nearly no information about possible *TERT* promoter mutations in CNS-PiloAstro and incomplete information in other cohorts. Using the detection sensitivity and observed mutation count at the *TERT* hotspot sites, we inferred the expected total number of events in each tumor type (**Fig 8c**). This analysis revealed that ~216 (CI_95_ = [188,245]) *TERT* hotspot mutations were likely missed in this study due to lack of power. Likewise, about four additional mutations would be expected at the recurrent *FOXA1* promoter hotspot shown to be mutated in hormone-receptor positive breast cancers (**Extended Data Fig. 12**)^9^.

**Figure 8:**
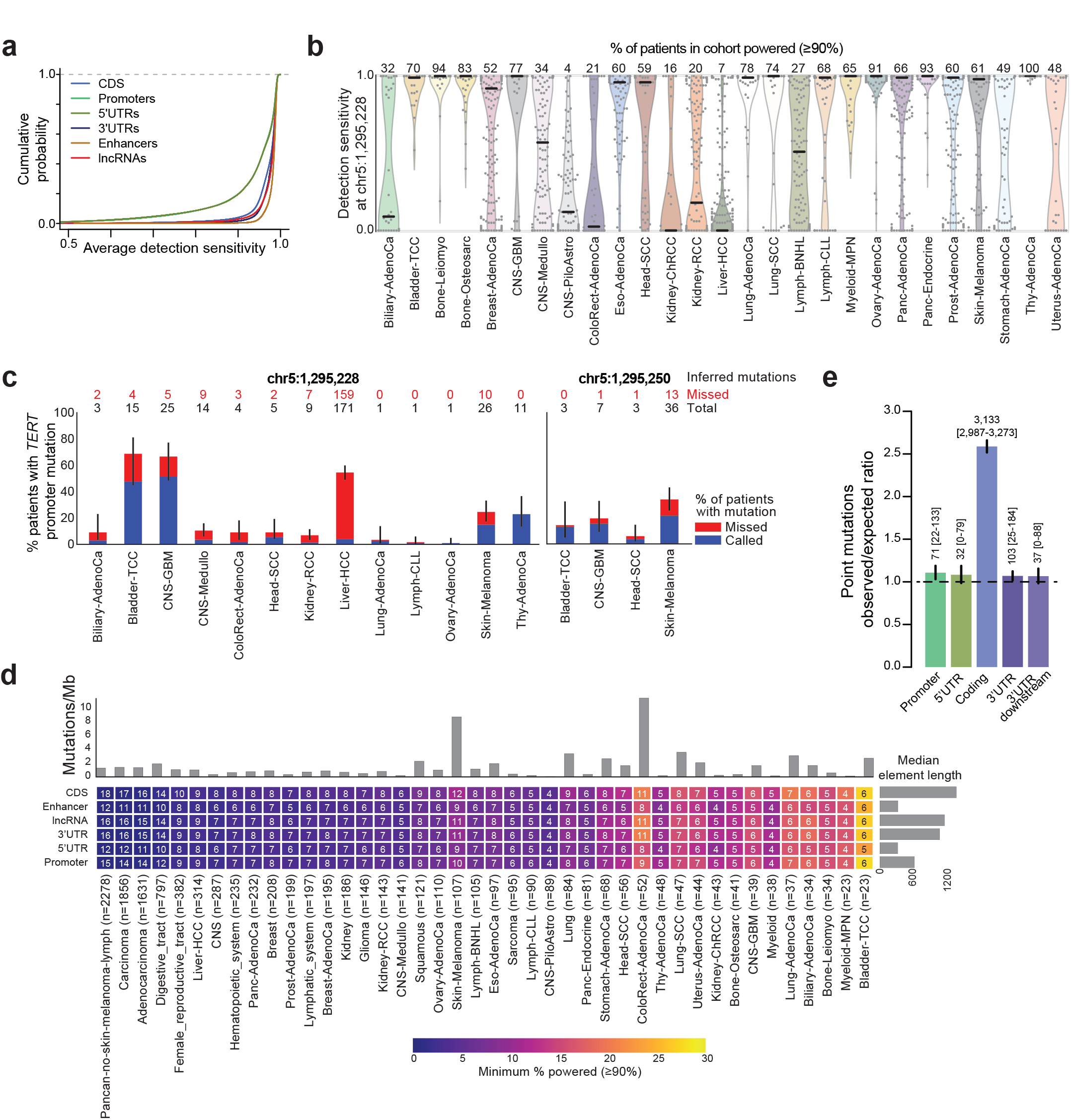
Missed mutations and paucity of non-coding drivers. **a**, Cumulative distribution of average detection sensitivity (d.s.) from 60 PCAWG pilot samples in different functional element types. **b**, Distribution of *TERT* promoter hotspot (chr5:1295228; hg19) detection sensitivity for each each patient, by cohort. Grey dots indicate values for individual patients inside estimated distribution (areas colored by cohort). Horizontal black bars mark the medians. Numbers above distributions indicate the percentage of patients powered (d.s. >90%) in each cohort. See **Extended Data Fig. 12b** for the second *TERT* hotspot. **c**, Percentage of patients with observed (blue) and inferred missed (red) mutations at the chr5:1,295,228 and chr5:1,295,250 *TERT* promoter hotspot sites. Error bars indicate 95% confidence interval. Numbers above bars show the total inferred number of *TERT* promoter mutations for each site in this cohort. Red numbers indicate the absolute number of inferred missed mutations (due to lack of read coverage). **d**, Heatmap shows minimal frequency of a driver element in a cohort that is powered (≥90%) to be discovered. Cell numbers indicate the number of patients in a given cohort that would need to harbor a mutation in a given element. For example, the pan-cancer cohort is powered to discover a driver gene (CDS) present in <1% or 18 patients, while the Bladder-TCC cohort is only powered to discover drivers present in at least 27% or 6 patients. Power to discover driver elements is dependent on the background mutation frequency (shown above the heatmap) and element length (shown to the right). **e**, The ratio between observed numbers of mutations (SNVs and indels) in regulatory and coding regions of 142 protein-coding cancer genes. The absolute number of driver mutations predicted, with CI_95_% in brackets, is shown above each bar.

These calculations suggest that a considerable number of potentially impactful somatic mutations were not detected due to systematic low sequencing coverage, and that specific positions can be unpowered in elements with overall sufficient coverage.

We further evaluated the discovery power for recurrent events identified in mutational burden tests in the tumor cohorts analyzed in this study^9, 50^. As expected, discovery power was highest in cohorts with many patients and low background mutation densities allowing discovery of typical-sized driver elements mutated in fewer than 1% of patients in the Carcinoma, Adenocarcinoma and pan-cancer meta-cohorts with 90% power (**Fig. 8d**). In contrast, the small Bladder-TCC cohort (n=23) with relatively high background mutation density (~2.7 mutations/Mb) is only powered to discover drivers that occur in at least 25% of patients (**Fig. 8d**). Overall, power differences between element types for individual tumor cohorts were small, suggesting that lack of power alone cannot explain the observed paucity of regulatory driver elements.

### Relative paucity of non-coding driver point mutations

To further analyze the relative paucity of driver mutations in non-coding elements, we sought to estimate the overall number of driver mutations, in coding and non-coding regions of known cancer genes. Given the limited statistical power to detect individually significant elements, we combined the signal from multiple elements, as recently described^53^. The difference between the total number of mutations observed across a combined set of elements and the number of passenger mutations that is expected by chance, *i.e.* the excess of mutations, approximates the total number of driver mutations in the set of elements. To estimate the expected number of background mutations, we fit the NBR model to presumed passenger genes, controlling for sequence composition, element size and regional mutation densities (**Methods; Supplementary Table 7; Supplementary Note**)^53^. We focused on 142 known cancer genes that include the significantly mutated cancer genes in this study and frequently amplified or deleted cancer genes (**Methods**). We specifically excluded *TERT* from this analysis because of its high frequency of promoter driver mutations and the incomplete detection sensitivity reported above.

Overall, this approach predicted an excess of 3,133 driver mutations (CI_95_% [2,987-3,273]; 2,258 SNVs and 875 indels) in the protein-coding sequences of these genes across the pancancer meta-cohort (**Fig. 8e**). In contrast to coding regions, the observed number of mutations across the combined set of non-coding elements associated with these genes is very close to the expected number of passenger mutations,, with an excess of 71 (CI_95%_:22-133) mutations in promoters, 32 (0-79) in 5’UTRs, and 103 (25-184) in 3’UTRs. These results indicate that coding genes contribute the vast majority (>90%) of driver point mutations in these 142 cancer genes. Importantly, these estimates are conservative, since the estimation of the background number of mutations from putative passenger genes may include yet undetected driver mutations. In addition, non-coding mutations in promoters of cancer genes were also not generally associated with LOH nor with altered expression, as one would expect if they were enriched with driver mutations (Supplementary Note). Altogether, our results suggest that mutations in protein-coding regions dominate the landscape of driver point mutations in known cancer genes.

## Discussion

If we are to fulfill the ambitions of precision medicine, we need a detailed understanding of the genetic changes that drive each person’s cancer, including those in non-coding regions (https://doi.org/10.1101/190330). Most cancer genomic studies have focused on protein-coding genes, leaving non-coding regulatory regions and non-coding RNA genes largely unexplored. The unprecedented availability of high-quality mutation calls from whole-genomes of >2,500 patients across 27 cancer types has enabled us to comprehensively search for functional elements with cancer-driver mutations across the genome. To obtain reliable results and avoid common pitfalls in detecting drivers^54^, we have benchmarked a large number of methods and developed a novel and rigorous statistical strategy for integrating their results.

Among the most interesting candidate non-coding driver elements identified in this analysis, we have uncovered promoter or 5’UTR mutations in *TP53, RFTN1* and *MTG2,* a putative intragenic regulatory element in *RNF34* that possibly regulates *KDM2B;* 3’UTR mutations in *NFKBIZ* and *TOB1;* and recurrent mutations in the non-coding RNA *RMRP.* We have also found evidence suggesting that a number of previously reported and frequently mutated non-coding elements may not be genuine cancer drivers. Particularly, the non-coding RNAs *NEAT1* and *MALAT1* are subject to a high density of passenger indels, seemingly due to a transcription-associated mutational process that targets some of the most highly expressed genes.

Our study has yielded an unexpectedly low number of non-coding driver candidates. The results from four analyses - genomic hotspot recurrence, driver element discovery, discovery power and mutational excess - suggest that the regulatory elements studied here contribute a much smaller number of recurrent cancer-driving mutations than protein-coding genes. This contrasts the distribution of germline polymorphisms associated with heritability of complex traits, which are most frequently located outside of protein-coding genes ^55^.

Technical shortcomings, such as severe localized lack of sequence coverage in GC-rich promoters (as in the case of the *TERT* promoter), may lead to considerable underestimation of true drivers in certain regions. The yield of driver events will thus benefit from technical improvements, including less variable sequence coverage (*e.g.* PCR-free library preparation), longer sequencing reads and advanced methods for read mapping and variant calling that can overcome common artifacts. Moreover, studying larger tumor cohorts will provide increased power to discover infrequently mutated driver elements.

Our analyses have focused on currently-annotated non-coding regulatory elements and noncoding genes, comprising 4% of the genome, and exclusively on SNVs and indels. Although our genome-wide hotspot analysis did not detect novel highly recurrent non-coding mutation sites, it is possible that additional non-coding drivers reside outside of the regions tested here. Improvements in the functional annotation of non-coding elements, our understanding of their tissue-specificity, and the impact of mutations in them, will refine current driver discovery models and enable more comprehensive screens.

A challenge in the identification of non-coding drivers is distinguishing real novel driver events from yet-unidentified mutational mechanisms targeting certain genomic regions, such as the recently described hypermutation of transcription factor binding sites in melanoma^56, 57^. Better understanding of mutational processes and their activity along the genome will be critical to improve the sensitivity and specificity of statistical methods for driver discovery.

One potential explanation for the relative paucity of non-coding drivers is the smaller functional territory size of many regulatory elements, and hence smaller chance of being mutated, compared to protein-coding genes (**Extended Data Fig.1; Fig. 8d**). The presence of *TERT* promoter mutations at just two sites suggests that non-coding driver mutations may be confined to specific positions, similar to the small number of impactful sites in the oncogenes *BRAF, KRAS* and *IDH1.*

SNVs and small indels may not easily alter the function of non-coding regulatory elements. Directly mutating protein-coding sequences or altering expression levels by copy number changes, epigenetic changes or repurposing of distal enhancers through genomic rearrangements^58, 59^ may be more likely to provide large phenotypic effects. While protein-truncating or frameshift mutations can be created by a single nucleotide change, silencing regulatory regions of tumor suppressors may depend on large deletions to achieve a similar effect. Although the very high frequency of *TERT* promoter mutations in cancer demonstrates that some promoter point mutations can be potent driver events, our analyses suggest that the number of regulatory sites where a point mutation can lead to major phenotypic effects may be smaller than anticipated across the genome. This highlights *TERT* as an unusual example, perhaps because even a modest increase in the expression level of *TERT* may be sufficient for circumventing normal telomere shortening in cancer cells.

Comprehensive and reliable discovery of non-coding driver mutations in cancer genomes will be an integral part of cancer research and precision medicine in the coming years. We anticipate that the approaches developed here will provide a solid foundation for the incipient era of driver discovery from ever-larger numbers of cancer whole-genome sequences.

## Acknowledgements

We thank Kirsten Kübler for assistance with meta cohort generation. We are grateful to Matthew Meyerson, Eric S. Lander and the PCAWG steering committee for helpful feedback on the manuscript.

## Author contributions

This work was carried out by the Discovery of Driver Elements Subgroup of the PCAWG Drivers and Functional Interpretation Group based on data from the ICGC/TCGA Pan-Cancer Analysis of Whole Genomes Network. Contributions spanned a range of categories, as defined in **Supplementary Note**. The *Driver discovery* category includes method development, driver significance evaluation and results integration. The *Candidate vetting and filtering* category covers removal of potential false positive hits. The *Case-based analysis* category includes driver potential and performed follow-up studies using additional evidence. The *Power analysis and driver mutations at known cancer genes* category includes power evaluations and estimation of the number of true drivers. The *Leadership and organisational work* category includes organization and workgroup leadership. E.R., M.M.N., F.A., G.T., H.H., J.M.H., R.I.P., I.M., J.S.P. and G.G. wrote the manuscript. All authors have had access to read and comment on the manuscript.

---

**Extended Data Fig. 1:**
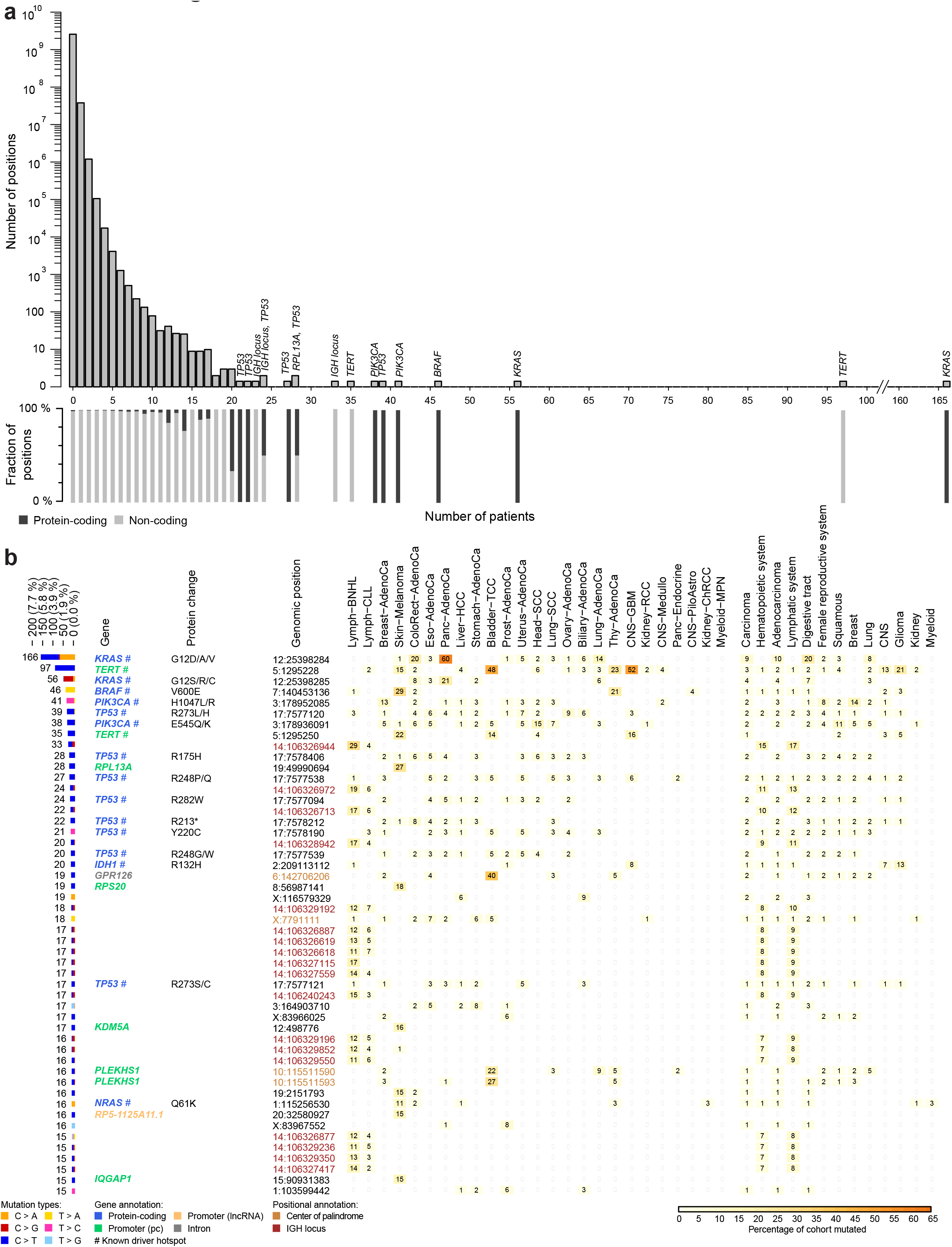
Mutational hotspots in additional tumor types. **a**, Barplot of positions vs. patients. The stacked barcharts under the barplot show the proportion of protein-coding (dark grey) and non-coding (light grey) positions, respectively, in each of the bars in the barplot. **b**, Distribution of SNVs in top 50 single-site hotspots across all analyzed individual cohorts and meta-cohorts.

**Extended Data Fig. 2:**
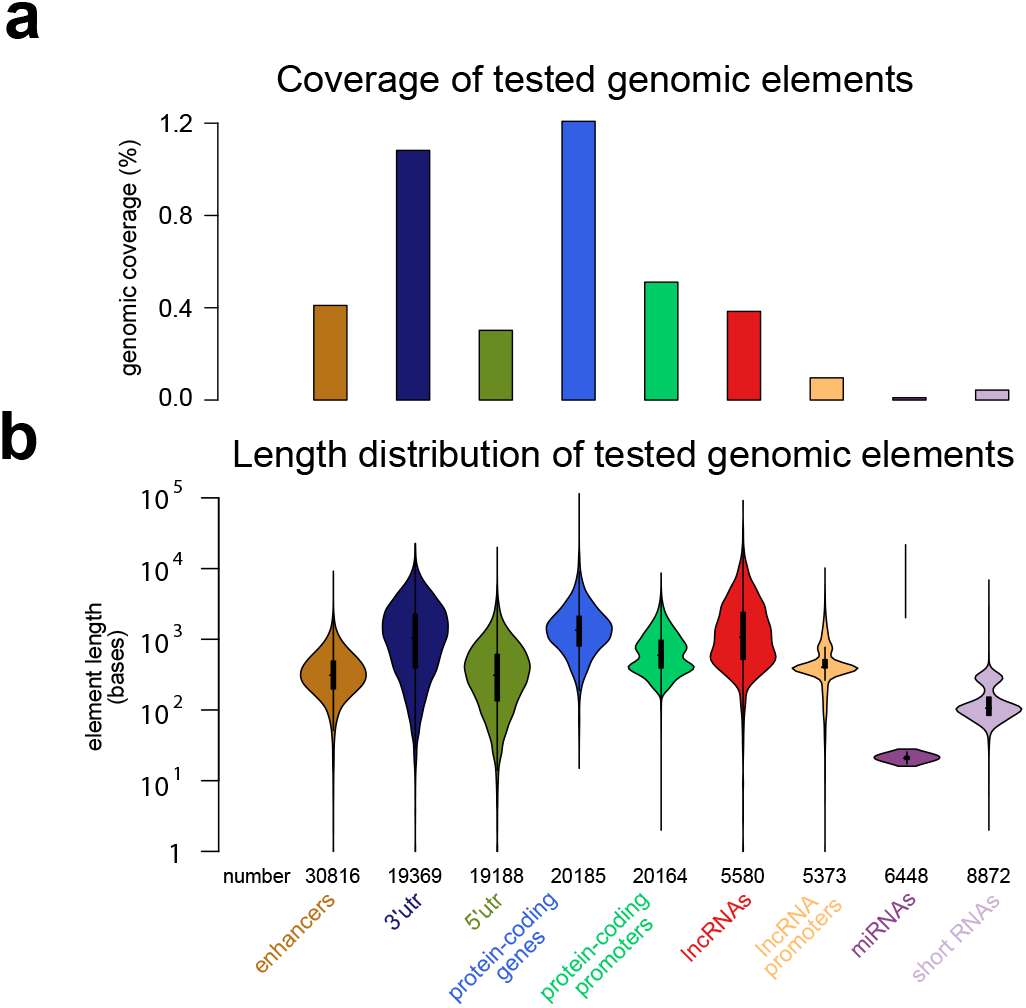
Genomic element statistics. **a**, Percentage of genomic coverage for each element type. **b**, Distribution of element lengths for each element type.

**Extended Data Fig. 3:**
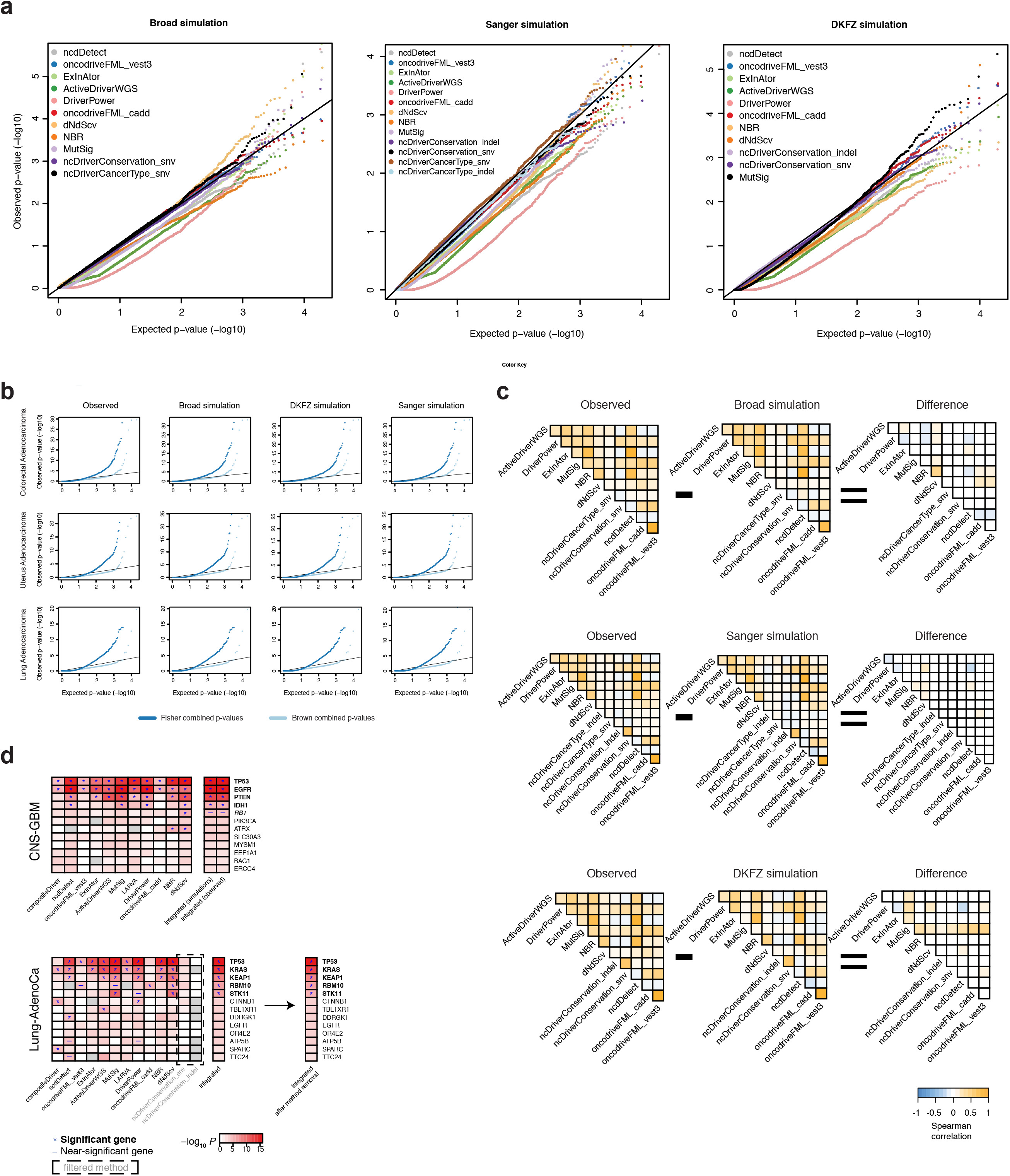
Integration details. **a** Quantile-quantile plots of p-values reported by various driver detection algorithms on the three simulated data sets (Broad, DKFZ, and Sanger; shown for coding regions in the meta-carcinoma cohort) showed no major enrichment of mutations above the background rate. Results generally followed the expected null (uniform) distribution, and the p-values reported on simulated data were subsequently used to assess the covariance of method results. **b**, Quantile-quantile plots of integrated p-values using the Brown and Fisher methods for combining p-values across the different driver detection algorithm results were generated for a few representative tumor cohorts (shown here for coding regions). Brown combined p-values (light blue) generally followed the null distribution as expected, while Fisher combined p-values were significantly inflated (dark blue), confirming that dependencies existed between the results reported by the various driver detection algorithms. To simplify the integration procedure, we calculated covariances using p-values from the observed data instead of simulated data and found that the integrated results based on the observed covariances (first column of plots) were essentially the same as the results obtained using the simulated covariances (second, third, and fourth columns of plots). **c**, Triangular heatmaps showing the Spearman correlations of p-values among the various driver detection methods in observed versus simulated data (coding regions, meta-carcinoma cohort) are highly similar. Differences in the observed and simulated correlation values (shown in the far-right heatmaps) were minimal, and thus the final integration of p-values across methods was performed using covariances estimated on observed data. **d**, Integrated p-values based on observed and simulated covariance estimations (shown on the right, top heatmap, for coding regions in glioblastoma) did not differ noticeably. In cases where individual methods reported results that yielded substantially fewer hits than the median across all methods (bottom heatmap, methods in light grey with results in dashed box), removing the methods from the integration did not affect the number of significant genes identified (right column of results in bottom heatmap, shown for coding regions in lung adenocarcinoma).

**Extended Data Fig. 4:**
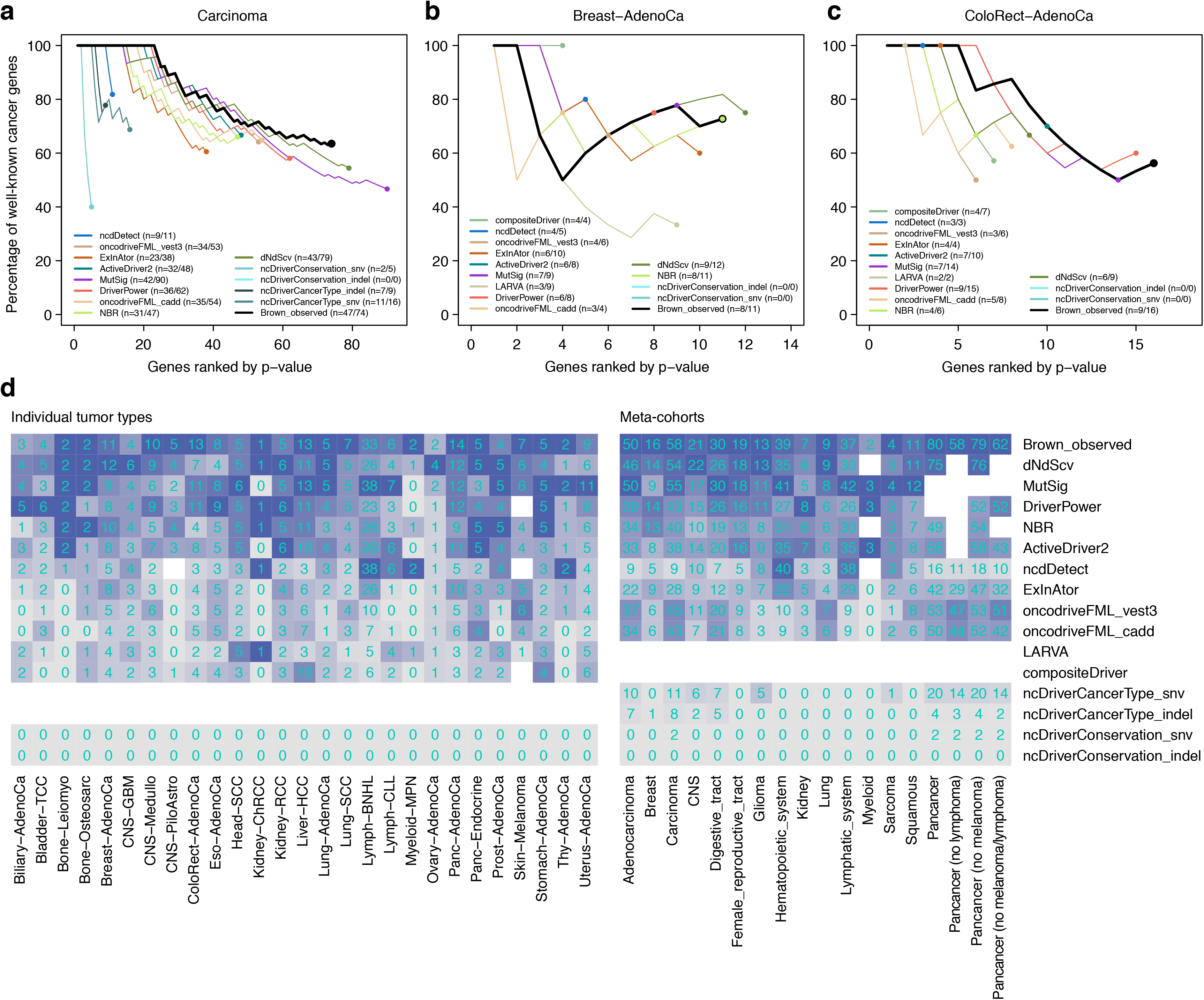
Sensitivity of driver discovery methods. The performance of different driver discovery methods and of the integration method on protein-coding genes was evaluated using known cancer genes. A list of 603 Cancer Gene Census genes (CGC, v80) was used as a reference gold-standard set of known cancer genes. **a-c**, For each method, genes with q-value<0.10 were sorted according to their p-value and the fraction of CGC genes shown in the y-axis. The total number of significant genes identified by each method is shown in the x-axis. The integration approach tends to outperform most methods across cohorts, yielding longer lists of significant genes and more strongly enriched in known cancer genes. **d**, Heatmap depicting the number of known cancer genes identified by each method and by the integration approach in each cohort. The color of each cell reflects the relative sensitivity of each method in each cohort, measured as the number of cancer genes detected by a method (also shown as a number inside each cell) divided by the maximum number detected by any of the methods. Methods are sorted from top to bottom according to their mean relative sensitivity across datasets.

**Extended Data Fig. 5:**
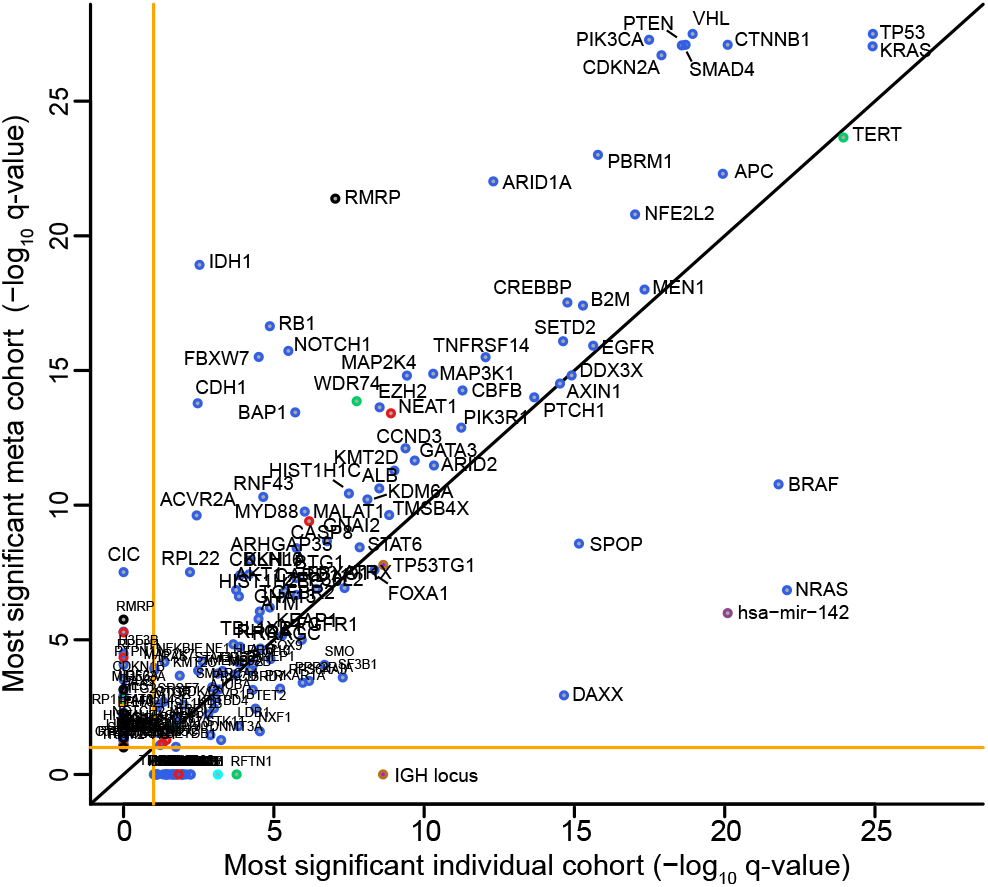
Sensitivity vs. specificity in individual cohorts vs. meta cohorts for candidate drivers. q-values for the most significant individual cohort (x-axis) vs. meta cohort (y-axis) are shown. Driver elements are colored by their element type.

**Extended Data Fig. 6:**
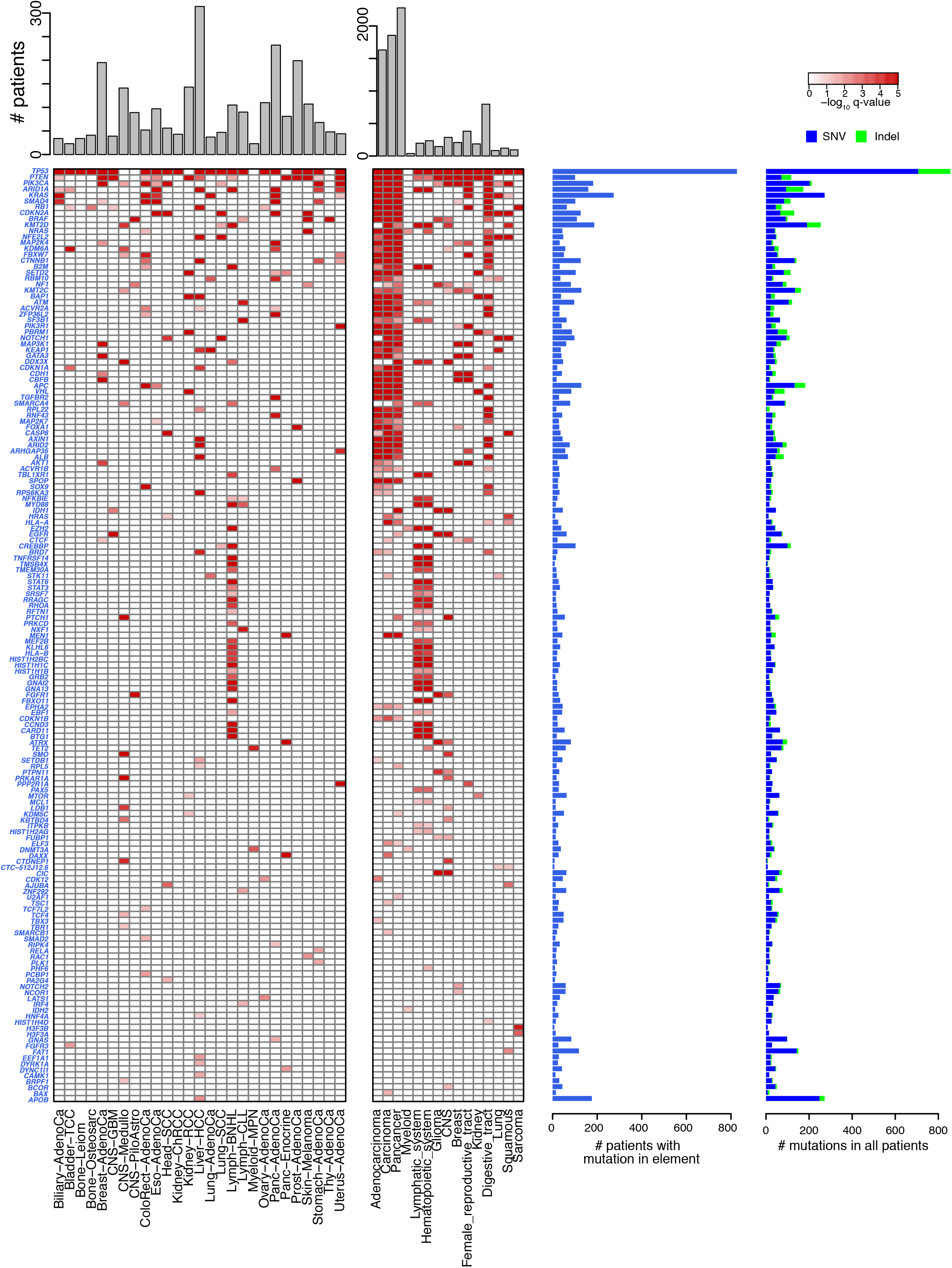
Protein-coding driver elements identified in PCAWG cohorts. Significant protein-coding elements identified in this study. Compare to figure 4.

**Extended Data Fig. 7:**
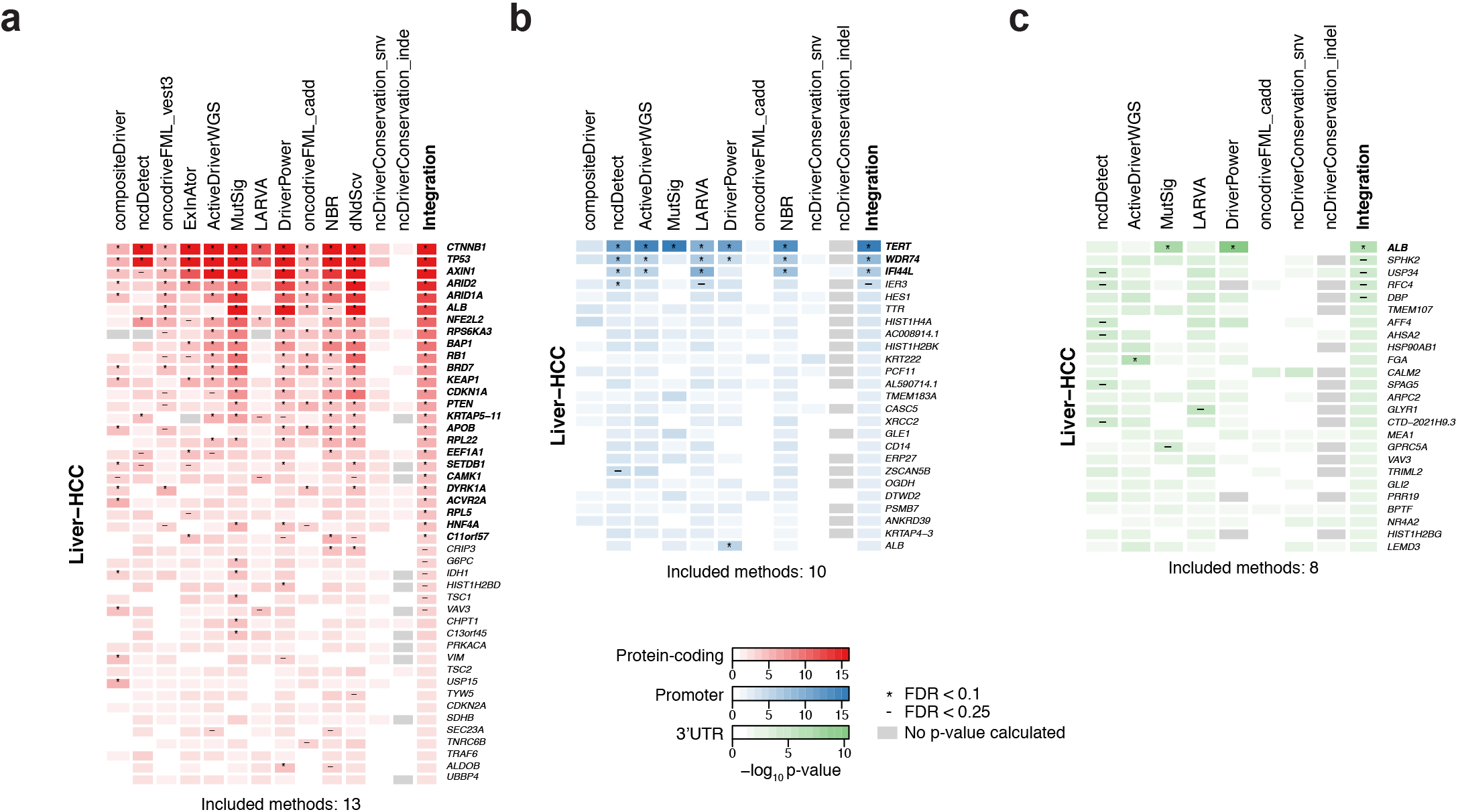
Lack of concordance for non-coding results. Heatmaps depicting p-values for methods included in integration for Liver-HCC cohorts. **a**, Most methods agree on the top protein-coding hits. **b, c**: Agreement between methods on non-coding hits is far lower for Liver-HCC promoters (b) and 3’UTRs (c).

**Extended Data Fig. 8:**
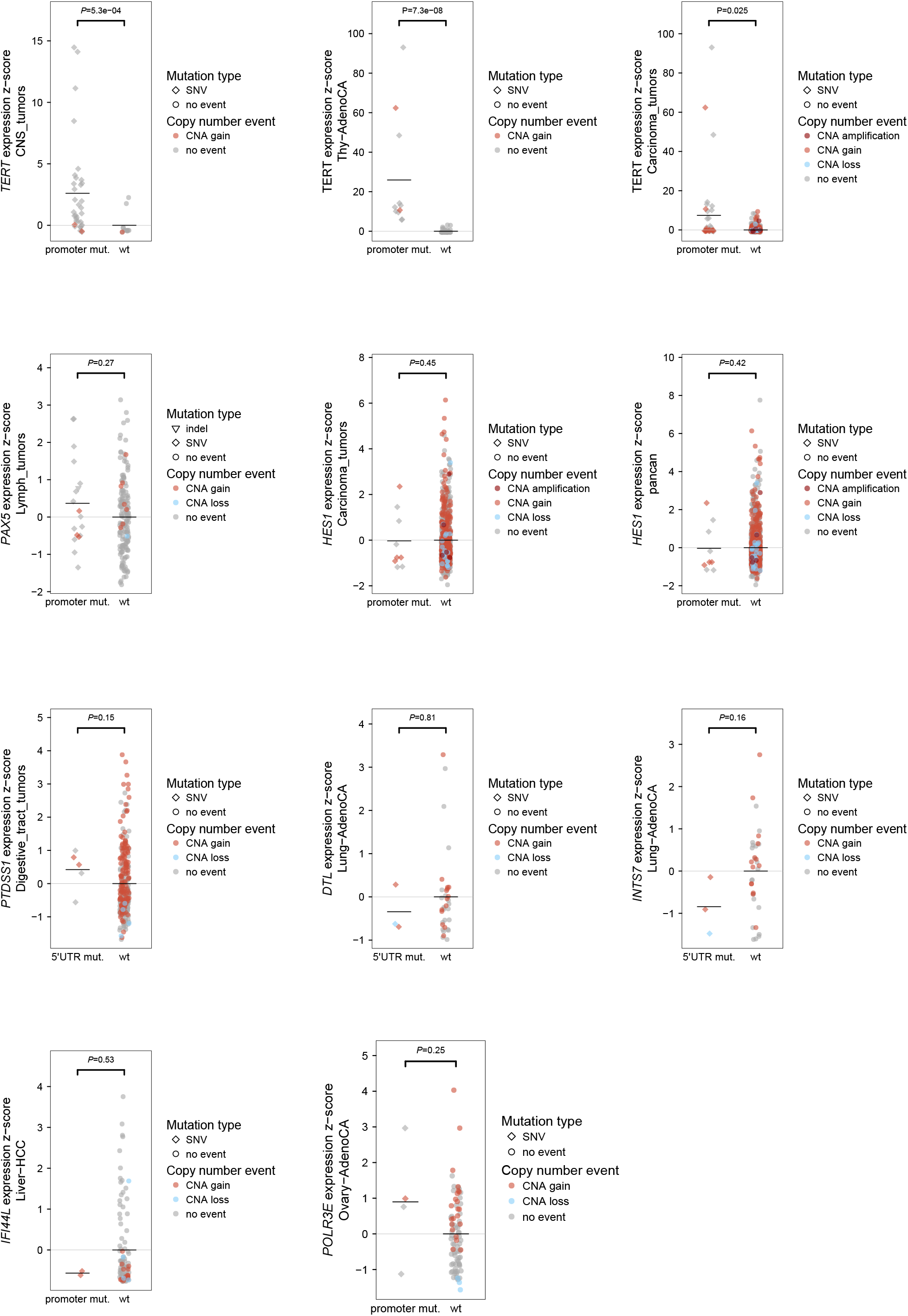
Mutation to expression correlation. Expression is compared between mutated and non-mutated samples. For each element, the z-score of the expression values for mutated and wild type in the significant cohort is plotted. For copy numbers, SCNA amplification indicates SCNA > 10, SCNA gain indicates SCNA ≥ 3, SCNA loss indicates SCNA ≤ 1 and no events indicates SCNA < 3 and SCNA >1. If a patient is mutated with multiple point mutation types, indels are indicated over SNVs.

**Extended Data Fig. 9:**
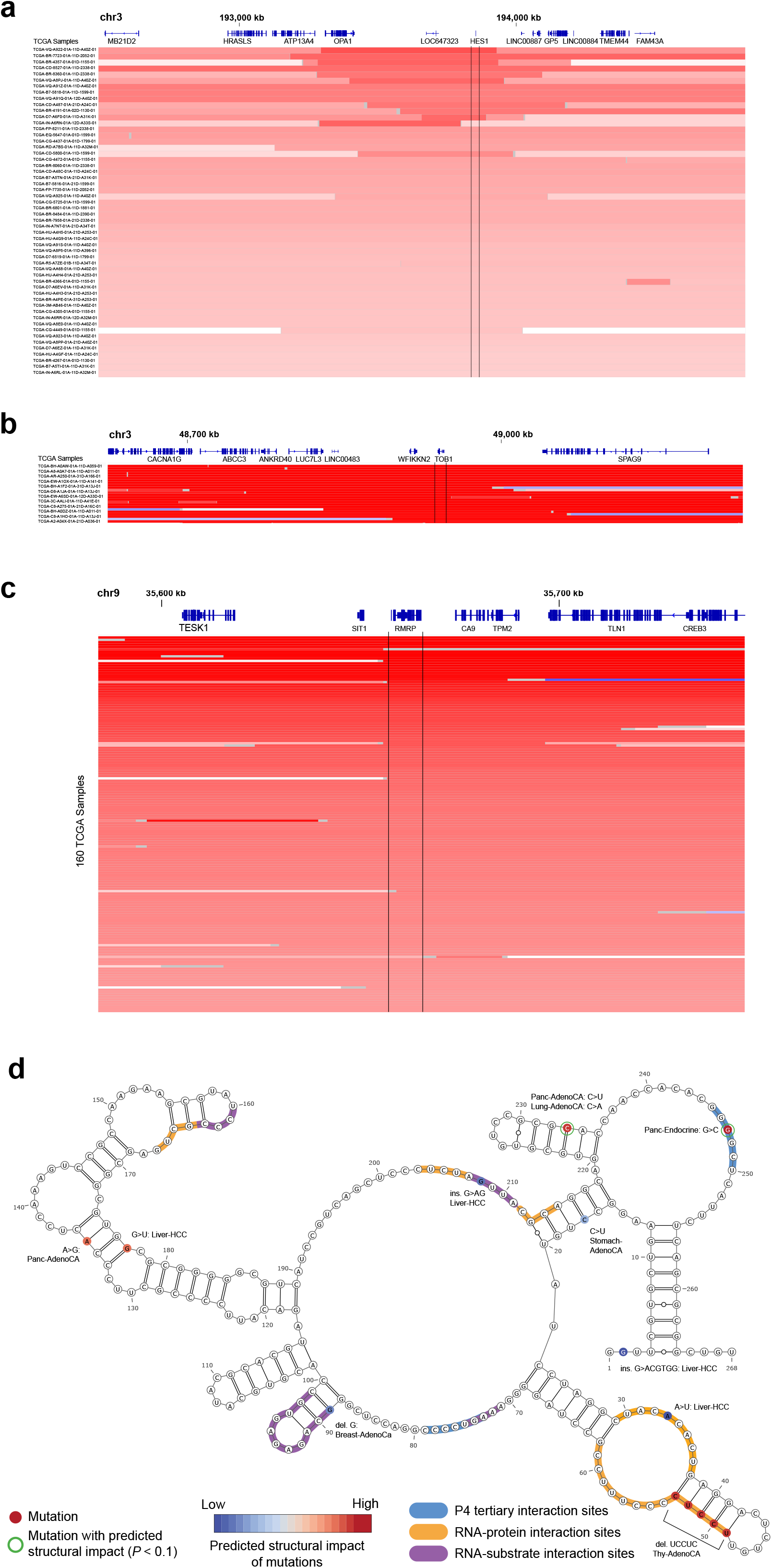
Focal amplifications including *TOB1, HES1* and *RMRP* in TCGA tumors. **a**, Copy number profiles of 55 of 441 stomach adenocarcinomas from TCGA show copy number gains around *HES1.* **b**, *TOB1* and its gene neighbor *WIFKKN2* are focally amplified in cancer. 172 of 10844 total samples from 33 cancer types are shown. **c**, *RMRP* focal amplifications in TCGA cancers (160 of 10844 total tumors shown). **d**, *RMRP* secondary structure (RF00030 in Rfam) labeled with P4 tertiary interactions as well as protein and substrate interactions (reported as ± 3bp windows). Mutations and their predicted structural impact are indicated. Lymphoma and melanoma mutations are excluded.

**Extended Data Fig. 10:**
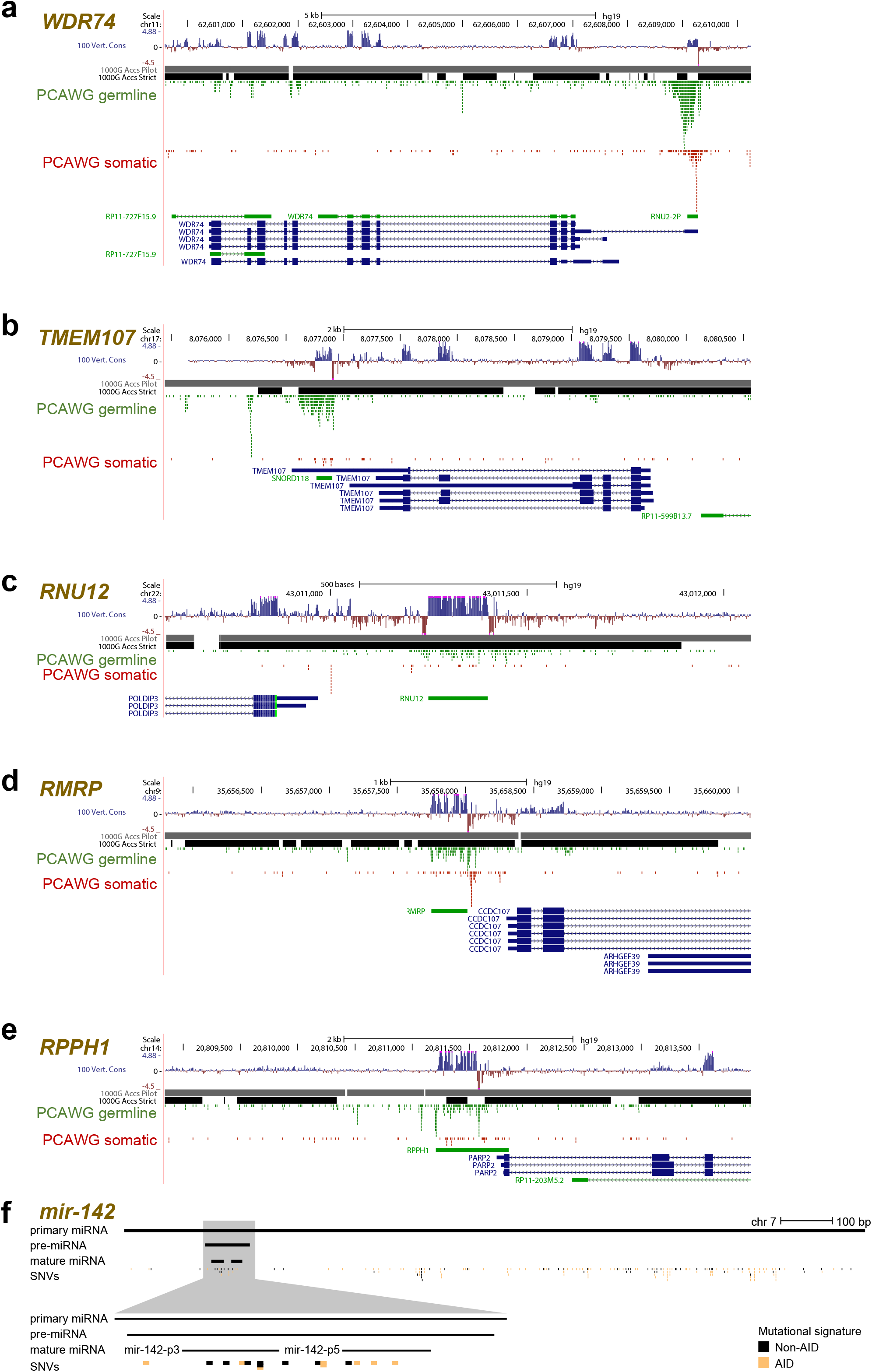
Density of potential germline polymorphisms and somatic mutations in several candidates. Genomic overviews showing mapping quality masks from the 1,000 Genomes Project (pilot/strict), and densities of germline SNP and somatic SNV calls on PCAWG normal and tumor samples, respectively. Excess of germline SNP calls can be indicative of artifacts, thus an increased number of somatic mutations in such a region raises concerns about true mutational recurrence. **a**, *WDR74* promoter, *U2* RNA element. **b**, *TMEM107* 3’UTR. **c**, *RNU12* small RNA. **d**, *RMRP* promoter and ncRNA transcript. **e**, *RPPH1* ncRNA. **f**, Genomic overview showing SNVs in *mir-142.* SNVs attributed to the AID mutational signature are marked in orange.

**Extended Data Fig. 11:**
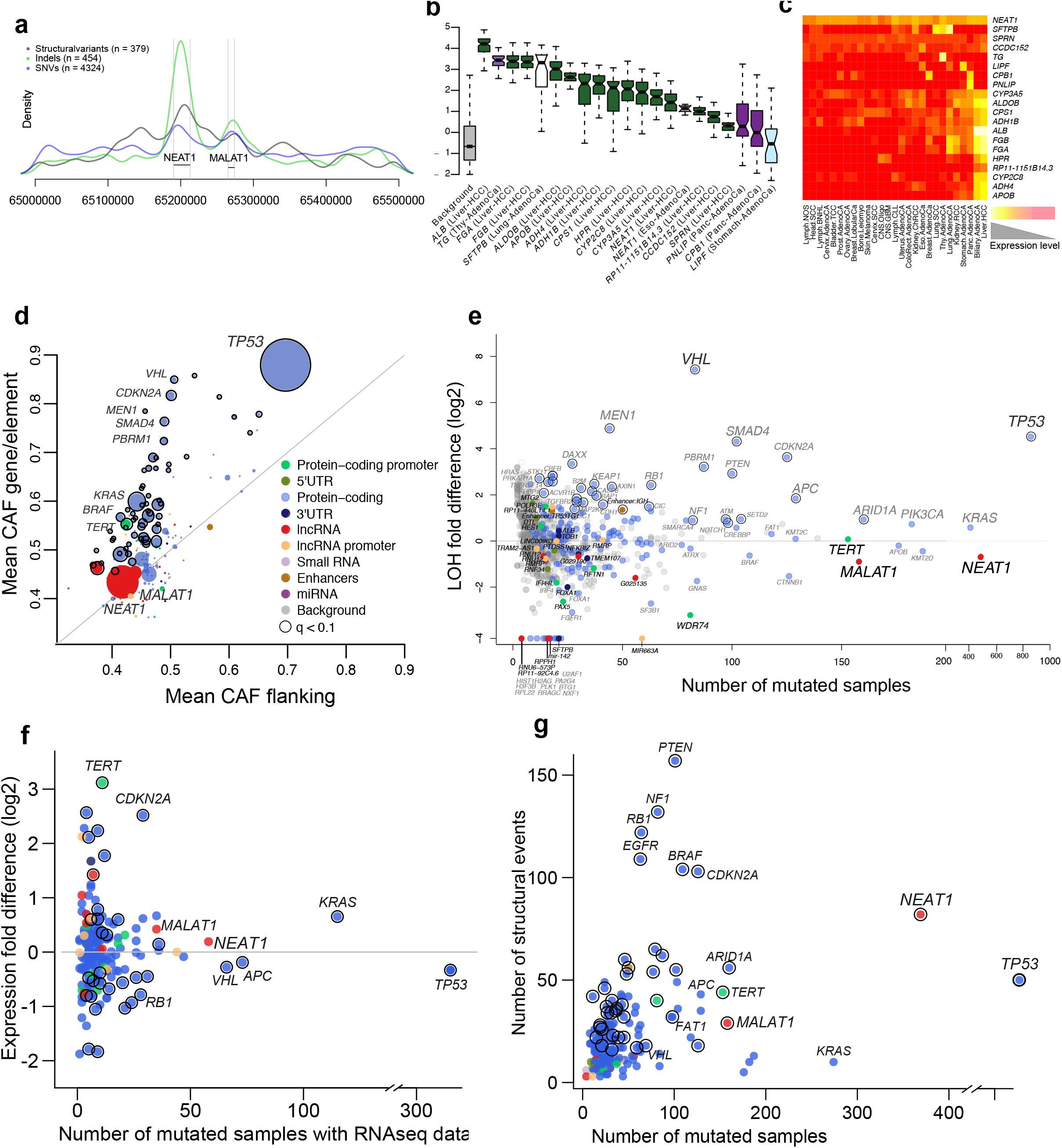
Supporting information for *NEAT1* and *MALAT1.* **a**, Mutation density in the *NEAT1* and *MALAT1* genomic loci. Co-localized peaks of densities in the loci are seen for structural variants, SNVs and indels. **b**, Levels of expression of the genes enriched in 2-5 bp indels in their respective tissues. Background gene expression levels are shown to the left for comparison. **c**, Heatmap showing the levels of expression across tissues for the genes enriched in 2-5 bp indels. **d**, The average cancer allelic fraction (CAF) is compared between each genomic element and the corresponding flanking regions (+/-2Kb and introns; overlapping coding exons were excluded). The size of the points represent the number of mutated samples for each particular element. **e**, The relative rate of loss of heterozygosity (LOH) is compared between mutated and wild-type samples, coloured by element type and highlighting significant LOH enrichments with an outside black circle. **f**, Expression comparison between mutated and wild-type samples. For each element, the median log**2** fold expression difference of cohorts found significant in the driver discovery is plotted. If elements were significant in multiple cohorts, the one with the most significant expression difference is shown. Number of mutated samples refers to tumors for which RNA-seq data could be obtained. Element types are colored as indicated in the legend. Elements with a significant expression difference (q < 0.1 or q < 0.01) are indicated by circles. **g**, Number samples with structural variants (SV) near or in each element is plotted against the number of patients with a point mutation. Number of SVs is the sum of event counts in bins of 50 kb regions that overlap with a given element. Elements with a significantly higher than expected number of samples with SVs are indicated with black circles (q < 0.1).

**Extended Data Fig. 12:**
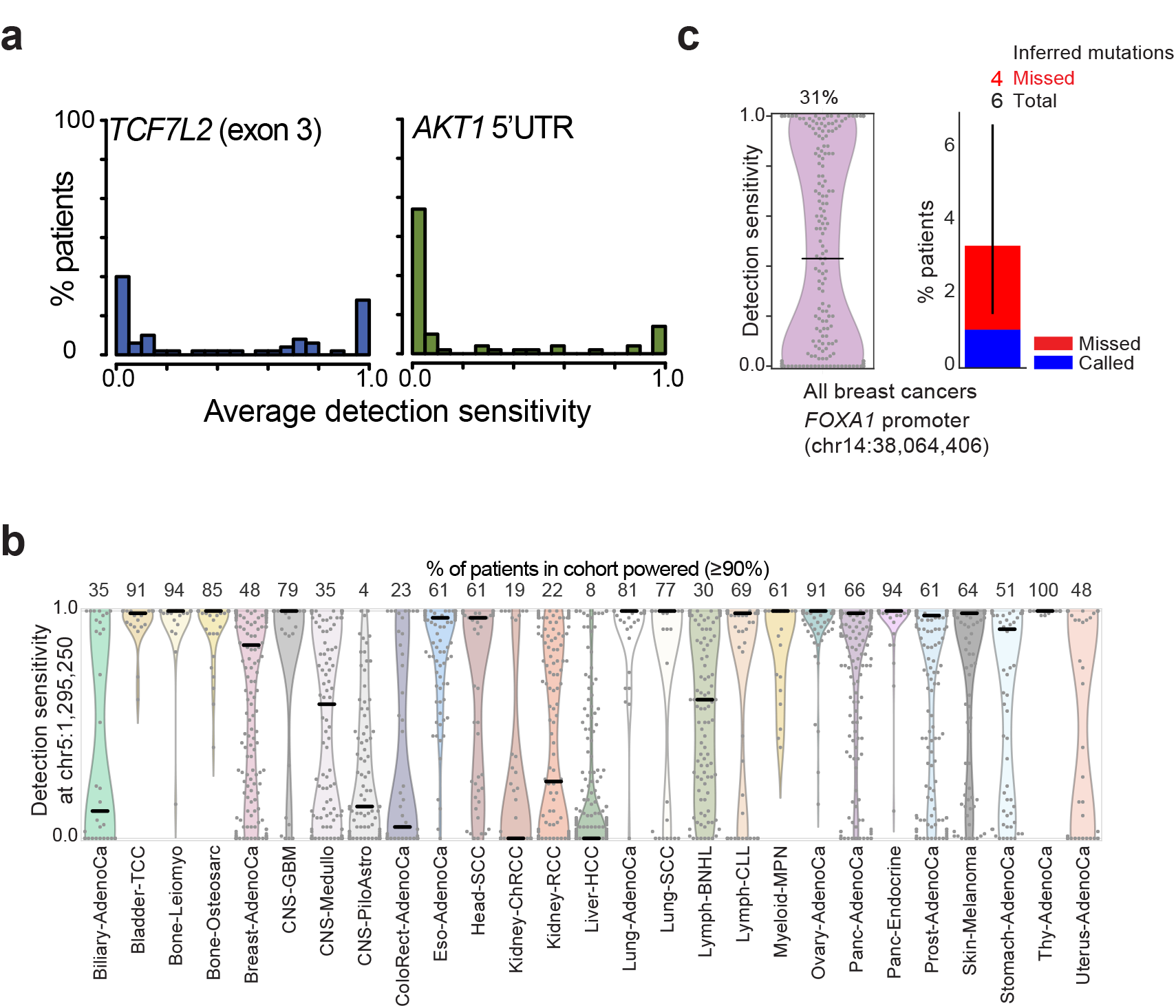
Lack of detection power in specific elements. **a**, Cumulative Average d.s. for selected elements across 60 patients. Exon 3 of the colon cancer gene *TCF7L2* is 90% powered in only 14/60 (23%) of patients. The *TERT* promoter and 5’UTR element of *AKT1* show lack of detection power in the majority of patients. **b**, Distribution of *TERT* promoter hotspot (chr5:1295250; hg19) detection sensitivity for each each patient, by cohort. Grey dots indicate values for individual patients inside estimated distribution (areas colored by cohort). Horizontal black bars mark the medians. Numbers above distributions indicate the percentage of patients powered (d.s. ≥90%) in each cohort. **c**, Left: distribution (pink) of detection sensitivity across all breast cancer patients (Breast-AdenoCa, Breast-LobularCa, Breast-DCIS; grey dots) at the *FOXA1* promoter hotspot site (chr14:38064406; hg19). The number above the distribution plot indicates the percentage of patients powered (d.s. ≥90%). Black horizontal bar indicates the median of the distribution. Right: percentage of patients with observed (blue) and inferred missed (red) mutations. Error bar indicates 95% confidence interval.

## Supplementary Tables

**Supplementary Table 1:** Genomic element type summary statistics

**Supplementary Table 2:** Driver discovery method description

**Supplementary Table 3:** Driver discovery methods included in integration

**Supplementary Table 4:** Summary and annotation of protein-coding driver candidates

**Supplementary Table 5:** Summary and annotation of non-coding driver candidates

**Supplementary Table 6:** Summary of additional evidence for non-coding driver candidates

**Supplementary Table 7:** List of cancer genes used in this study

**Supplementary Table 8:** List of cases used in detection sensitivity analysis

**Supplementary Table 9:** Impact of covariates on obs/exp ratio Supplementary Note

## Methods

**Table of contents:**

1. Patient cohorts
2. Mutational hotspot analysis
3. Mutational signatures
4. Genomic element definition
5. Driver discovery methods
6. Simulated data sets
7. Statistical framework for the integration of results from multiple driver discovery methods
8. Post-filtering of candidates
9. Sensitivity and specificity analysis
10. Gene expression analyses
11. Normalization for copy number variation
12. Mutation to expression association
13. Copy number analyses
14. Power analysis
15. Associations between mutation and signatures of selection: loss of heterozygosity and cancer allelic fractions
16. Signals of selection in aggregates of non-coding regions of known cancer genes
17. Mutational process and indel enrichment
18. Mutation association with splicing
19. Structural variation analysis
20. RNA structural analysis
21. Cancer associated germline variant distance to non-coding driver candidates

### 1. Patient cohorts

**Generation of high-quality tumor set** (Loris Mularoni)

We selected a total of 2583 samples to be included in the driver detection analyses. This list contains all the samples that were not flagged as problematic by the PCAWG-TECH group.

A single aliquot was assigned to each sample; in cases where multiple aliquots were present, we selected a single aliquot based on the following criteria, in order of importance:

- we prioritized primary tumors over metastatic or recurrent tumors
- we selected aliquots with an OxoG score higher than 40
- we prioritized aliquots with the highest quality (as indicated by the Stars values)
- we prioritized aliquots with RNA-seq data availability
- we prioritized aliquots with the lowest contamination (as indicated by the ContEst values)
- if a selection could not be made after applying the above filters we selected an aliquot randomly

**Selection of tumor cohorts for analysis** (Esther Rheinbay)

Individual tumor type cohorts from the high-quality tumor set were selected for analysis if they met a minimum size. This size was determined based on the cumulative number of patients, such that no more than 2.5% of total patients were excluded. This led to a minimum cohort size criterion of 20 patients, and removed the Bone-Cart (9 donors), Bone-Epith (11), Bone-Osteoblast (5), Breast-DCIS (3), Breast-LobularCa (13), Cervix-AdenoCa (2), Cervix-SCC (18), CNS-Oligo (18), Lymph-NOS (2), Myeloid-AML (13) and Myeloid-MDS (2) individual cohorts. Samples from these cohorts were still included in meta-cohort analysis (see below).

**Tumor meta-cohorts** (Esther Rheinbay)

Tumor meta-cohorts were assembled for identification of drivers and increase of discovery power across cell lineages and organ systems. The following meta cohorts were used in driver analyses:

**By cell type of origin:**

**Epithelial: Carcinoma** (comprised of tumor cohorts Bladder-TCC, Biliary-AdenoCa, Breast-AdenoCa, Breast-LobularCa, Cervix-AdenoCa, ColoRect-AdenoCa, Eso-AdenoCa, Kidney-ChRCC, Kidney-RCC, Liver-HCC, Lung-AdenoCa, Ovary-AdenoCa, Panc-AdenoCa, Panc-Endocrine, Prost-AdenoCa, Stomach-AdenoCa, Thy-AdenoCa, Uterus-AdenoCa,Head-SCC, Cervix-SCC, Lung-SCC), **Adenocarcinoma** (Biliary-AdenoCa, Breast-AdenoCa, Breast-LobularCa, Cervix-AdenoCa, ColoRect-AdenoCa, Eso-AdenoCa, Kidney-ChRCC, Kidney-RCC, Liver-HCC, Lung-AdenoCa, Ovary-AdenoCa, Panc-AdenoCa, Prost-AdenoCa, Stomach-AdenoCa, Thy-AdenoCa, Uterus-AdenoCa), **squamous epithelium** (Head-SCC, Cervix-SCC, Lung-SCC)

**Mesenchymal cells/sarcoma** (Bone-Cart, Bone-Epith, Bone-Leiomyo, Bone-Osteosarc)

**Glioma** (CNS-PiloAstro, CNS-Oligo, CNS-GBM)

**Hematopoietic system** (Lymph-BNHL, Lymph-CLL, Lymph-NOS, Myeloid-AML, Myeloid-MDS, Myeloid-MPN)

**By organ system:**

**digestive tract** (Liver-HCC, ColoRect-AdenoCa, Panc-AdenoCa, Eso-AdenoCa, Stomach-AdenoCa, Biliary-AdenoCa), **kidney** (Kidney-RCC, Kidney-ChRCC), **lung** (Lung-AdenoCa, Lung-SCC), **lymphatic system** (Lymph-BNHL, Lymph-CLL, Lymph-NOS), **myeloid** (Myeloid-AML, Myeloid-MDS, Myeloid-MPN), breast (Breast-AdenoCa, Breast-LobularCa), **female_reproductive_system** (Breast-AdenoCa, Breast-LobularCa, Cervix-AdenoCa, Cervix-SCC, Ovary-AdenoCa, Uterus-AdenoCa), **central nervous system** (CNS-PiloAstro, CNS-Oligo, CNS-Medullo, CNS-GBM)

**Pan-cancer:**

Two “Pan-cancer” cohorts were created: “Pancan-no-skin-melanoma” containing all tumor types with the exception of Skin-Melanoma to remove issues caused by very high mutation rate tumors; and “Pancan-no-skin-melanoma-lymph” with the additional removal of lymphoid tumors (Lymph-BNHL, Lymph-CLL, Lymph-NOS) that have local somatic hypermutation caused by AID.

### 2. Mutational hotspot analysis (Randi Istrup Pedersen)

We selected the top 50 single position hotspots based on the number of patients with an SNV mutation. The individual positions marked as problematic by the site-specific noise filter (see below) analysis were excluded.

Each hotspot was defined by its genomic position and annotated by the number of patients with an SNV mutation in the given hotspot. We also annotated each hotspot with whether it falled in one of the genomic element types analyzed in the driver discovery. We further overlapped with loop-regions of palindromes, which are hypothesized to fold into DNA-level hairpins, and with location in immunoglobulin loci. When a hotspot overlapped a protein-coding gene we extracted the corresponding amino acids changes from Oncotator^1^ (<u>http://portals.broadinstitute.org/oncotator/</u>).

We identified known driver hotspots, by overlap with the somatic driver positions compiled in the Cancer Genome Interpreter repository (<u>https://www.cancergenomeinterpreter.org/mutations</u>), which among others include mutations from ClinVar, DoCM and the literature (ref: https://doi.org/10.1101/140475).

For each hotspot we calculated the proportion of mutations in the defined cohorts and metacohorts. Only cohorts with at least 20 patients, and at least 10 patients or 10% of patients with an SNV were included in **Fig. 1a** (for the distribution in all cohorts and meta-cohorts see **Extended Data Fig. 1**). Lymph-BNHL and Lymph-CLL were shown together as Lymphoid malignancies.

Based on mutational signature analysis of all the cancer samples, we extracted the posterior probability that each hotspot mutation from a given patient was generated by one of 37 identified mutational signatures. In lymphoid malignancies somatic hyper-mutations generated by AID come in clusters along the genome. Posterior probabilities for the ten signatures relevant for the lymph cohorts were therefore derived from models that consider the correlation of AID mutations along the genome. For each hotspot the collected posterior probabilities were averaged.

### 3. Mutational Signatures (Jaegil Kim)

We performed a *de novo* global signature discovery to identify mutational signatures operating in PCAWG WGS cohort (*N* = 2,583). All single nucleotide variants (SNVs) were stratified into *M* = 4608 mutation channels according to the combination of six base substitutions at pyrimidine bases (C>A/G/T and T>A/C/G) in penta-nucleotide sequence contexts and a transcription strand direction, transcribed (-), non-transcribed (+), non-coding regions. All insertions and deletions were classified into additional 20 channels depending on the number of inserted or deleted bases and any indels beyond 9 bases were grouped into a single channel. The resulting mutation frequency matrix *X* (4,628 channels across 2,583 samples) were ingested to the Broad Institute’s signature analysis pipeline, *SignatureAnalyzer,* to determine the optimal number of signatures (*K*) and the signature profiles by de-convoluting *X* into a product of two non-negative matrices as *X* ~ *WH, W* (*M* × *K*) and *H* (*K* × *N*) being a signature-loading and an activity-loading matrix, respectively. The most distinguished feature of *SignatureAnalyzer* is that it suggests an optimal number of signatures best explaining *X* at the balance between the error measure and the model complexity exploiting Bayesian non-negative matrix algorithm (NMF)^2,3,4^. Our de-novo signature discovery identified 37 signatures including a split of 4 UV-related signatures, 3 APOBEC-related signatures, 5 POLE-related signatures, 3 MSI-related signatures, 2 COSMIC17 signatures, and other 21 singleton signatures (see PCAWG7 paper for details).

Although the matrix *H* contains the most representative activity across samples it also has spurious activity assignments intrinsic to the global signature discovery (i.e., applying *SignatureAnalyzer* on samples with different histological subtypes and mutational processes). To minimize this contamination and interference we re-assigned activity separately in each histological subtype using a more refined projection approach. We first identified a subset of ultra-mutant samples with a dominant activity of *POLE* (*n* = 8), MSI (*n* = 18), and alkylating signatures (*n* = 1) from the original *H*, which are usually very exclusive to samples with a specific DNA repair or replication defect, or a treatment. To prevent a false-positive assignment of these signatures in remaining samples we introduced a binary matrix *Z* (37 × *N*), where the row elements corresponding to *POLE,* MSI, and alkylating signatures, are set to zero except for samples with a dominant activity, while all other elements are set to one. More specifically, the projection was done by keeping the signature-loading matrix *W*’ (*M* × *37*) frozen (*W*’ represents the normalized signature profiles of 37 global signatures), while allowing *SignatureAnalyzer* to determine a subset of signatures and infer the activity-loading matrix *H*’ (*37 × N^r^*) that best approximates the mutation frequency matrix *X*’ (*M × N*), where *N*’ represents the number of samples in each histological subtype, as *X’ ~ W*’ (*Z¤H*), Here “¤” denotes element-wise multiplications.

### 4. Definition of genomic elements

(Morten Muhlig Nielsen; Nicholas Sinnott-Armstrong) For coding elements and elements pertaining to protein coding genes (3’UTR, 5’UTR and protein coding promoters) regions were defined based on GENCODE annotations (v.19)^5^:

#### Coding elements (CDS)

The set of coding bases collapsed across all coding transcripts with a given GENCODE gene ID.

#### Protein coding splice site elements (pcSS)

Intronic regions extending 6 bases from donor splice sites and 20 bases from acceptor splice sites were collected for all coding transcripts. Bases were collapsed across all coding transcripts with a given GENCODE gene ID. The global set of CDS bases were subtracted.

#### 5’UTR elements (5UTR)

The set of 5’UTR bases collapsed across all coding transcripts with a given GENCODE gene ID. The global set of CDS and pcSS bases were subtracted.

#### 3’UTR elements (3UTR)

The set of 3’UTR bases collapsed across all coding transcripts with a given GENCODE gene ID. The global set of CDS, pcSS and 5UTR bases were subtracted.

#### Protein coding promoter elements (pcPROM)

Regions extending 200 bases in both directions from all protein coding transcripts’ transcription start sites (5’ ends). Bases were collapsed across all coding transcripts with a given GENCODE gene ID. The global set of CDS and pcSS bases were subtracted.

#### IncRNA elements

IncRNA transcripts were defined based on annotations from GENCODE (v.19) and MiTranscriptome (v.2)^6^ not overlapping GENCODE. Transcripts were included if fulfilling criteria 1-5 and 6 or 7 below:

1. No sense overlap to protein coding gene regions
2. More than 5kb away from protein coding genes on sense strand.
3. Longer than 200 bases.
4. Not annotated as the following biotypes: immunoglobulin, T-cell receptor, Mt_rRNA, Mt_tRNA, miRNA, misc_RNA, rRNA, scRNA, snRNA, snoRNA, ribozyme, sRNA or scaRNA.
5. Not overlapping genomic regions aligning back to the human genome (self chained regions).
6. More than 20% of bases overlap conserved elements (except if annotated as pseudogene)
7. Expressed in more than 10% of PCAWG samples with RNAseq data.

Genes corresponding to the selected transcripts were supplemented with a set of known functional lncRNA genes from the literature in addition to GENCODE annotated non-coding snoRNA and miRNA host genes. The elements were made by collapsing bases across transcripts with given gene ID. The global set of CDS, pc_SS, 5UTR, 3UTR, pcPROM and lncRNA_SS bases were subtracted.

#### lncRNA splice site elements (lncRNASS)

Intronic regions extending 6 bases from donor splice sites and 20 bases from acceptor splice sites were collected for all lncRNA transcripts. Bases were collapsed across all lncRNA transcripts with a given gene ID. The global set of CDS, pc_SS, 5UTR, 3UTR and pcPROM bases were subtracted.

#### lncRNA promoter elements

Regions extending 200 bases in both directions from all lncRNA transcripts’ transcription start sites (5’ends). Bases were collapsed across all lncRNA transcripts with a given gene ID. The global set of CDS, pc_SS, 5UTR, 3UTR, pcPROM and lncRNA_SS bases were subtracted.

#### Short RNA elements

Short RNA transcripts were defined based on annotations from databases Rfam (v.11)^7^, tRNAscanSE (v.2.0)^8^ and snoRNAdb (v.3)^9^ in addition to GENCODE transcripts with biotype annotations mt_rRNA, mt_tRNA, misc_RNA, rRNA and snoRNA.

Bases were collapsed across all smallRNA transcripts with a given gene ID. The global set of CDS, pc_SS, 5UTR, 3UTR and pcPROM bases were subtracted.

#### microRNA elements

Mature miRNAs were defined based on mirBase (v.20)^10^ and a set of potential novel miRNAs^11^

#### Enhancers

Contiguous 15-state ChromHMM called enhancers correlated between H3K4me1 and RNA-seq across 57 human tissues were downloaded from Roadmap Epigenomics Consortium extended data^12^. Associated links, defined by co-occurring activity in a given cell type, were merged across cell types at FDR = 0.1. HoneyBadger2^13^ p10 calls for all DNase I sites were filtered to peaks with signal strength 0.8 or greater and intersected with enhancer elements. The union of all DNase I peaks which overlapped with a given element, with all CDS regions filtered out, were used as the input to driver detection.

### 5. Candidate driver identification methods

A summary of approaches used by each method is listed in **Supplementary Table 2.**

#### ActiveDriverWGS (Juri Reimand)

Driver analysis with ActiveDriverWGS (ref) was performed after discarding hypermutated samples (>90,000 mutations) from the PCAWG cancer cohort. To avoid leakage of signals from known cancer drivers, we removed missense mutations in analyses of non-coding regions. ActiveDriverWGS is a local mutation enrichment method for genome-wide discovery of cancer driver mutations with increased mutation burden of single nucleotide variants (SNVs) and indels. ActiveDriverWGS performs a model-based test whether a given genomic element is significantly more mutated than adjacent background genomic sequence (+/-10kb and introns). Statistical significance of mutations is computed with a Poisson-linked generalised linear regression model. The null model treats all SNVs with trinucleotide context as cofactor, while indels are modelled with a separate cofactor for all nucleotides. Mutation counts per nucleotide are presented as the response variable The alternative model tests whether the element has different mutation burden than the background sequence. The null and alternative models are compared with chi-square tests and confidence intervals of expected mutations were derived from the null model using resampling. If the confidence intervals indicated significant excess of mutation in the background and depletion in the element of interest, we inverted corresponding small p-values (p= 1-p if p<0.5). Elements with no mutations were automatically assigned p=1.

#### CompositeDriver0.2 (Eric Minwei Liu, Ekta Khurana)

We have developed CompositeDriver (ref) – a computational method that combines signals of mutation recurrence and the functional impact score derived from FunSeq2 scheme^14^ to identify coding and non-coding elements under positive selection. CompositeDriver assigns a score to each region of interest (i.e., CDS, promoter, UTR, enhancer or ncRNA) through summation of positional mutation recurrence multiplied by the functional impact score for all mutations within the region. A null CompositeDriver score distribution is built to calculate the p-value for a region of interest. Mutations in the same element type but outside the region of interest are defined as background mutations. To build the null distribution, the same numbers of mutated positions are repeatedly drawn (default is 10^5^ times) from background mutations with similar replication timing and similar mutation context^15^. By drawing random mutations from the same element type, CompositeDriver incorporates DNase I hypersensitive sites and histone modification marks as covariates into the null model^16^. Finally, the Benjamini-Hochberg method is used for multiple hypothesis correction^17^.

#### dNdScv (Inigo Martincorena)

dNdScv is a maximum-likelihood algorithm designed to test for positive or negative selection in cancer genomes or other sparse resequencing studies. dNdScv models somatic mutations in a given gene as a Poisson process, accounting for sequence composition and mutational signatures using 192 trinucleotide substitution rates. Mutation rates are also known to vary across genes, often co-varying with functional features of the human genome, such as replication time and chromatin state. This information is exploited by dNdScv to refine the estimates of the background mutation rate of each gene, using a negative binomial regression. This regression removes known sources of variation of the mutation rates and models the remaining unexplained variation of the mutation rate across genes as being Gamma distributed, which protects the method against overconfidence in the estimated background mutation rate for a gene. Overall, the local mutation rate for a gene is estimated accounting for mutational signatures in the samples analysed, the sequence composition of a gene in a trinucleotide context, 20 epigenomic covariates and the local number of synonymous mutations in the gene. Inferences on selection are carried out separately for missense substitutions, truncating substitutions (nonsense and essential splice site mutations) and indels, and then combined into a global P-value per gene. dNdScv has been described in much greater detail elsewhere^18^.

#### DriverPower (Shimin Shuai)

DriverPower is a combined burden and functional impact test for coding and non-coding cancer driver elements. In the DriverPower framework, randomized non-coding genome elements are used as training set. In total 1373 reference features covering nucleotide compositions, conservation, replication timing, expression levels, epigenomic marks and compartments are collected for downstream modelling. For the modelling, a feature selection step by randomized Lasso is performed at first. Then the expected background mutation rate is estimated with selected highly important features by binomial generalized linear model. The predicted mutation rate is further calibrated with functional impact scores measured by CADD and Eigen scores. Finally, a p-value is generated for each test element by binomial test with the alternative hypothesis that the observed mutation rate is higher than the adjusted mutation rate.

#### ExInAtor (Andres Lanzos; Rory Johnson)

ExInAtor was specifically created for prediction of cancer driver lncRNAs, but is agnostic to gene type and can also be used for protein-coding genes. The exons of each gene are identified and collapsed across transcript isoforms. For each gene, the trinucleotide content of the exonic region is calculated. The remaining intronic regions, along with 10 kb of sequence upstream and downstream, are defined as the background region. From this background, a new background region is created by randomly sampling the maximum number of nucleotides, such that the trinucleotide content exactly matches that of the exonic region. Next, the number of mutations in the exonic and sampled background regions are compared by hypergeometric test. Genes with elevated exonic mutational density are considered candidate driver genes. ExInAtor was used with randomisation seed of 256. Otherwise ExInAtor was run exactly as described in Lanzós et al, 2017^19^.

#### LARVA (Jing Zhang; Lucas Lochovsky)

LARVA^20^, or Large-scale Analysis of Recurrent Variants in noncoding Annotations, is a computational method that detects significantly elevated somatic mutation burdens in genomic elements—both coding and non-coding—to identify putative cancer-driving elements. Given a cancer cohort variant call set, and a list of genomic elements, LARVA models the expected background somatic mutation rate by fitting a beta-binomial distribution to the elements’ variant counts. This model properly accounts for the high mutation rate variability seen throughout the genome, which improves over some previous models’ assumption of a constant mutation rate. LARVA’s model also incorporates the influence of mutation rate covariates, such as DNA replication timing. LARVA’s output lists each genomic element from the input, along with a p-value based on the deviation of the element’s observed variant count from the expected variant count under LARVA’s model.

#### MutSig (Julian Hess, Esther Rheinbay)

The MutSig suite^21^ classifies whether genomics features, both coding and non-coding, are highly mutated relative to a predicted background mutation rate (BMR), which varies on a macroscopic-level across patients (patient-specific mutation rates can span orders of magnitude across pan-cancer cohorts) and genes (known covariates such as replication timing are strongly correlated with mutation rate) and on a microscopic level across sequence contexts (since mutational signatures are heterogeneous across a cohort and highly context-dependent). MutSig accounts for all three of these to compute the joint BMR distribution across genes/patients/contexts, and then convolves across the latter two dimensions to estimate the expected distribution of total background burden for a given gene across a whole cohort. Genes are then scored by how their total non-background burden exceeds this null distribution.

MutSig estimates a gene’s BMR by its synonymous mutation rate for coding genes, and by its mutation rate at nonconserved positions for non-coding genes. If the number of background mutations in a given gene is insufficient to provide a confident estimate of its BMR, MutSig will incorporate the background counts from other genes with similar covariate profiles into its estimator.

MutSig (MutSig2CV) was originally designed for coding regions only^21^. Modifications to this version of the algorithm to run on non-coding regions are novel to PCAWG. Coding MutSig also incorporates per-gene functional impact and clustering tests, which were not run on non-coding regions.

#### NBR (Inigo Martincorena)

NBR is a method that test for evidence of higher mutation density than expected by chance in a given region of the genome, while accounting for trinucleotide mutational signatures, sequence composition and the local density of mutations around each element. This method has been described in detail in a previous publication^22^, where it was used to identify candidate driver noncoding elements across 560 breast cancer whole-genomes.

Based on some of the features of dNdScv, NBR involves two main steps. First, all mutations across all elements tested are used to obtain maximum-likelihood estimates for the 192 rate parameters (r_j_) describing each of the possible trinucleotide substitutions in a strand-specific manner. r_j_ = n_j_/L_j_, where n_j_ is the total number of mutations observed across samples of a given trinucleotide class (j) and L_j_ is the number of available sites for each trinucleotide. These rates are used to estimate the total number of mutations across samples expected under neutrality in each element considering the mutational signatures active in the cohort and the sequence of the elements (E_h_ = S_j_ rL_j,h_). This estimate assumes no variation of the mutation rate across elements in the genome. Second, a negative binomial regression is used to refine this estimate of the background mutation rate of an element, using covariates and E_h_ as an offset. In this study, the local density of somatic mutations (normalized by sequence composition) was used as a covariate, using a window around the element of a variable size across cohorts to ensure sufficient numbers of mutations in each window around each element and excluding coding sequences and previously identified candidate noncoding driver regions. Replication time and average gene expression level for 100 kb genomic bins were also used as covariates. The negative binomial regression models mutation counts as Poisson-distributed within an element with mutation rates varying across elements according to a Gamma distribution. As in dNdScv, this provides a refined estimate of the background mutation rate for each element (E_h_^*^) as well as a data-driven measure of uncertainty around this estimate (-q-the overdispersion parameter of the negative binomial regression). P-values for each element are calculated using a cumulative negative binomial distribution with the mean (E_h_^*^) and dispersion (q) parameters estimated by the negative binomial regression.

To protect against neutral indel hotspots or indel artefacts, unique indel sites rather than total indels per element were used. To protect against misannotation of a mutation clusters as sets of independent events, a maximum of two mutations per region and per sample were considered in the analysis.

#### ncdDetect (Malene Juul)

ncdDetect^23^ is a driver detection method tailored for the non-coding part of the genome. It uses a burden based approach, in which the frequency of mutations is considered, to reveal signs of recurrent positive selection across cancer genomes. For each candidate region, the observed mutation frequency is compared to a sample- and position-specific background mutation rate. A scoring scheme is applied to further account for functional impact in the significance evaluation of a candidate cancer driver element. In the present application, the scoring scheme is defined as log-likelihoods, i.e. minus the natural logarithm of the sample- and position-specific probabilities of mutation.

The position- and sample-specific probabilities of mutation used by ncdDetect are obtained by a statistical null model, inferred from somatic mutation calls of a collection of cancer samples (https://doi.org/10.1101/122879). The model includes a set of genomic annotations, known to correlate with the mutation rate in cancer. These are replication timing, trinucleotides (the nucleotide under consideration and its left and right flanking bases), genomic segment (a variable segmenting the genome into regulatory element types) and a position-specific measure of the local mutation rate (a weighted average of the mutation rate, calculated across samples in a 40 kb window flanking each specific position plus/minus 10 kb).

#### ncDriver (Henrik Hornshøj)

The *ncDriver* method (doi: <u>https://doi.org/10.1101/182642</u>) provides separate evaluations of the significance for two functional mutation properties, the level of conservation and the level of cancer type specificity. In the *ncDriverConservation* test, the conservation level of mutated positions were evaluated *locally* for being surprisingly high, given the distribution of conservation within the element. The p-value of the mean mutation phyloP conservation score for an element was obtained by Monte Carlo simulation of 100,000 mean phyloP scores based on the same number of mutations. Each mutated element was also evaluated *globally* by looking up the rank of the element mean phyloP conservation score among all elements annotated as the same type. This provided p-values for both *local* and *global* mutation conservation level, which were combined into a single conservation p-value using Fisher’s method. In the *ncDriverCancerType* test, the distribution of observed mutation counts of an element across the cancer types were evaluated for being surprising compared to expected counts estimated from a background null model (as described for the *ncdDetect* method) that accounts for cancer type specific mutation signatures and other co-variates. A Goodness-of-fit test with Monte Carlo simulation was used to determine whether the distribution of observed mutation counts across cancer types within the element is surprising given the expected mutation counts based on cancer types, mutation contexts and element type. For indels, the expected mutation counts were estimated solely from the mutation rates calculated from the mutation context, cancer type and element type.

#### OncodriveFML (Loris Mularoni)

OncodriveFML^24^ is a method designed to estimate the accumulated functional impact bias of tumor somatic mutations in genomic regions of interest, both coding and non-coding, based on a local simulation of the mutational process affecting it. The rationale behind OncodriveFML is that the observation of somatic mutations on a genomic element across tumors, whose average impact score is significantly greater than expected for said element constitutes a signal that these mutations have undergone positive selection during tumorigenesis. This, in turn is considered as a direct indication that this element drives tumorigenesis.

OncodriveFML first computes the average functional impact score of the observed mutations in the element of interest. The functional impact scores of mutations have been calculated using both CADD^25^ (coding and non-coding regions) and VEST3^26^ (only coding regions). Then the method randomly samples the same number of observed mutations following the probability of mutation of different tri-nucleotides, computed from the mutations observed in each cohort. The randomization step is repeated many times (1,000,000 in these analyses) and each time an average functional impact score is calculated. Finally, OncodriveFML derives an empirical p-value for each element by comparing the average functional impact score observed in the element to its local expected average functional impact score resulting from the random sampling. The empirical p-values are then corrected for false discovery rate and genomic elements that after the correction are still significant are considered candidate drivers.

#### regDriver (Husen M. Umer)

regDriver assesses the significance of mutations affecting transcription factor motifs using tissue-specific functional annotations (https://doi.org/10.1002/humu.23014). For each tumor cohort, functional annotations from the cell lines most similar to the respective tumor type are gathered. A functionality score is computed for each mutation based on its overlapping functional annotations. regDriver, collects highly-scored mutations in each of the defined elements and assesses the elements’ significance by comparing its accumulative score to a background score distribution obtained from the simulated sets. Therefore, only candidate regulatory mutations are considered in evaluating mutation enrichment per element.

### 6. Simulated datasets

#### Broad simulations

(Yosef Maruvka, Gad Getz)

Due to their differing context characteristics, we simulated SNVs and indels with different approaches. For SNVs, we divided the genome into 50kb regions. For each region, we counted the number of mutations in it across all the PCAWG patients and divided it by the total number of mutations. Every mutation was randomly assigned into a new region based on the region’s rate. The position inside the region was chosen to maintain the trinucleotide context of each mutation (the 5’ and 3’ nearest neighbors and the mutated position itself) and the alternate allele. In addition, for every base we counted how many times it was covered sufficiently, in 401 tumor and normal WGS pairs, in order to enable calling of a mutation^27^. The fraction of patients with enough coverage at a given site was used as the position’s probability for being mutated inside the new current region.

For indels, a new, randomized position was chosen in a region of 50,000 bases around the indel. The position of the new indel was chosen to match the indel 5’ and 3’ neighboring reference bases. For insertions, the inserted motif was the same as the original insertion, but for deletions only the length of the indels was kept but not the exact sequence.

#### DKFZ simulations (Carl Hermann, Calvin Chan)

This simulation utilizes the SNV calls to perform a localised randomisation. The original SNV entries which do not map to chromosome 1-22, X, Y are first filtered and excluded from randomization. All SNVs located in the protein coding regions (CDS) corresponding to GENCODE19 definition are erased before performing randomisation. The trinucleotide centered at each SNV position is determined and an identical trinucleotide is randomly sampled within the 50kb window. In case of insertion, instead of the mutated trinucleotide, the neighboring nucleotide of the insertion site is scanned within the randomisation window. For deletion and multi-nucleotides variants, the altered sequence is scanned within the randomization window with a ranked probability assigned for each position. The randomised sample is then selected from the top 100 matched positions with scaled probability.

#### Sanger simulations (Inigo Martincorena)

This simulation aimed to generate datasets of neutral somatic mutations that retain key sources of variation in mutation rates known to exist in cancer genomes, including mutational signatures, and variable mutation rates across the genome and among individuals and cancer types. To do so while minimizing the number of assumptions in the simulation, we used a simple local randomization approach. First, all coding mutations as well as mutations in the *TERT* promoter, *MALAT1* or *NEAT1* were excluded. Second, each mutation in each patient was randomly moved to an identical trinucleotide within a 50 kb window, while retaining the patient ID. Third, mutations falling within 50 bp of their original position were filtered out. This simple randomization retains the variation of the mutation rate and mutational signatures across large regions of the genome, across individuals and across cancer types.

### 7. Statistical framework for the integration of results from multiple driver discovery methods

(Grace Tiao, Gad Getz)

The classical approach for combining p-values obtained from independent tests of a given null hypothesis was described by R. A. Fisher in 1948. He noted that for a set of *k* p-values, the sum *X* of the transformed p-values, where

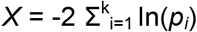

and *p_i_* is the p-value for the ith test, follows a chi-square distribution with 2*k* degrees of freedom^28^. Thus, to obtain a single combined p-value for a set of independent tests, the new test statistic *X* is computed from the p-values obtained from the tests and scored against a chi-square distribution with 2*k* degrees of freedom. Fisher’s test is asymptotically optimal among all methods of combining independent tests^29^; however, in cases where tests exhibit dependence, the Fisher combined p-value is generally too small (anti-conservative).

In this study, we combine p-values from several driver detection methods, many of which share similar approaches and whose results are therefore not independent. To address this issue, we used an extension of the Fisher method developed by Morten Brown for cases in which there is dependence among a set of tests^29^. Using the same test statistic, renamed *Ψ* to indicate the difference in the independence assumption, Brown observed that if *Ψ* were assumed to have a scaled chi-square distribution - i.e.,

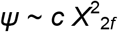

then

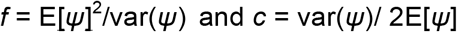

Note that E[*Ψ*] = 2*k* irrespective of the independence requirement, and that

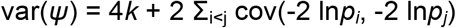

Thus when the *p_i_* are independent, var(*Ψ*) = 4*k*, which gives *f* = *k* and *c* ≠ 1, and the test statistic follows the chi-square distribution with 2k degrees of freedom described by Fisher. However, when the independence condition is relaxed, var(*Ψ*) Φ 4*k*, and the test statistic generally follows a different, scaled, chi-square distribution whose scaling parameter *c* and degrees of freedom 2*f* are determined by the covariances of the *p_i_*’s. The covariances can be computed via numerical integration over the joint distributions of all *p_i_* and *p_j_* pairs, but this requires knowledge of the joint distribution; and even in cases where the joint distribution is known, the integration may not be computationally feasible for large and complex datasets^30^.

In this study, following the example of Poole et. al^31^, we computed the empirical covariance of *p_i_* and *p_j_*, using the samples *w_i_* and *w_j_*, where *w_i_* is the set of all reported p-values for method *i*, and used the empirical covariance to approximate the Brown scaled chi-square distribution. The advantage to this approach is that the empirical covariance estimation is non-parametric – it does not assume an underlying joint distribution of *p_i_* and *p_j_* – and is thus applicable to complex and interrelated biological datasets where data is noisy and not regularly Gaussian. Poole et. al showed that the empirical covariance estimation approach is accurate, robust, and efficient for such datasets.

#### Implementing and evaluating the integration method on simulated and observed data

To evaluate the efficacy of the empirical Brown’s method of dependent p-value integration, we generated three sets of simulated mutation data (see above) and ran the driver detection algorithms on each of the simulated datasets. We checked that the p-value results from the various driver detection algorithms followed the expected null (uniform) distribution (**Extended Data Fig. 3a**). Then, for each simulated data set, we calculated the empirical covariance for each pair of driver algorithm results. We then used these covariance values over simulated datasets to compute the combined Brown p-values on observed data: for each gene in the observed PCAWG somatic mutation dataset, we computed the Brown test statistic from the set of p-values reported by the various driver detection algorithms. The Brown test statistic was then evaluated against the appropriate chi-square distribution, whose scale and degree parameters were approximated by the covariance values calculated on the simulated data (see above).

We ran this procedure, as well as the Fisher method, for six representative tumortype cohorts (Breast-AdenoCa, CNS-GBM, ColoRect-AdenoCa, Lung-AdenoCa, Uterus-AdenoCa, and meta-Carcinoma) and found that the Brown combined p-values generally followed the null distribution as expected (**Extended Data Fig. 3b**). The Fisher combined p-values were significantly inflated (**Extended Data Fig. 3b**), confirming that dependencies existed between the results reported by the various driver detection algorithms.

We next explored whether it was possible to reduce the number of algorithm runs required to complete these calculations for all tumor-type cohorts by computing the covariance values on observed data instead of simulated data. In each of the six representative tumor-type cohorts, we calculated the empirical covariances on the observed data only and then computed the integrated Brown p-values on the observed data using the observed covariances. Significant genes identified using only observed covariances remained mostly unchanged from the significant genes identified using the simulated covariances (**Extended Data Fig. 3d**), and examination of the differences in the covariance values between the simulated estimations and the observed estimations revealed only minor differences in values (**Extended Data Fig. 3c**). The significant drivers presented in this study were identified using this final approach – e.g., by computing integrated Brown p-values using estimations of covariance on observed data only.

Integration of p-values from observed data was performed for 42 tumor-type cohorts and 13 target element types. Methods were selected for each given data set (see “Selecting methods to include in the integration of observed p-values”, below) and raw p-values smaller than 10^−16^ were trimmed to that value before proceeding with the integration. Methods with missing data for a given element (i.e., ones that failed to report a p-value for a given element) were excluded from the calculation for that element, and therefore in some cases the integrated Brown p-value was computed from p-values reported by only a subset of all the driver detection algorithms contributing results for that data set.

#### Selecting methods to include in the integration of observed p-values

In some cases, individual driver detection algorithms reported p-values for a given data set that deviated strongly from the expected uniform null distribution. These were methods for which the quantile-quantile (QQ) plots demonstrated considerable inflation. We removed results that reported an unusual number of significant hits by calculating, for each set of results, the number of significant elements found by each individual method using the Benjamini-Hochberg FDR with q<0.1 as the significance threshold. Any single method that reported four times the median number of significant elements identified by individual methods was discarded from the integration. In a separate analysis, we found that removing methods that yielded fewer hits than the median (i.e., methods with deflated QQ-plots) did not affect the number of significant genes identified through the integration of the reported p-values (**Extended Data Fig. 3d**); hence we did not remove such methods.

### 8. Post-filtering of candidates

(Esther Rheinbay, Morten Muhlig Nielsen, Lars Feuerbach, Henrik Tobias Madsen)

Post-filtering of significant hits was performed to remove those with accumulation of mutations caused by sequencing problems or mutational processes. In particular, we applied the following: (i) at least three mutations are present in the element, (ii) mutations are present in at least three patients of the tested cohort, (iii) less than 50% of mutations are located in palindromic DNA sequence^22^, (iv) more than 50% of mutations are located in mappable genomic regions (CRG alignability, DAC blacklisted regions and DUKE uniqueness^32^; (v), less than 50% of mutations occur near indels, (vi) a site-specific noise filter (see below); and (vii) manual review of sequence evidence for novel drivers. For lymphomas, which contain regions of somatic hypermutation caused by AID enzyme activity, we (viii) further required less than 50% of mutations contributed by this process; and for Skin-melanoma, we (ix) excluded mutations occurring in the extended (CTTCCG) context that contributes to promoter hotspot mutations in this tumor type^33,34,35^. Even after this motif-based filter, a large number of promoter and 5’UTR regions remained in the list of candidates. We, therefore, marked these as likely due to failed repair in TF occupied sites since it is unclear how many of them are true or false drivers. Additionally, elements that failed manual mutation review were filtered out at this stage.

#### Site-specific noise filter

Genomic positions of mutations in each significant hit were analyzed in all normal control samples to assess the position-specific noise-level. Therefore, for each of the three nonreference nucleotides *a* and cohort *c ϵ C* the relative frequency of normal samples which had at least two reads supporting the alternative allele *a* were calculated as *p*(*a,c*) ϵ [0;100]. The noise score was then computed as *Σ*_*cϵC*_ *log*_10_(*p*(*a,c*) + 1). Mutations at positions with a score > 20 for at least one of the non-reference nucleotides were flagged. Elements for which the number of mutations at flagged positions exceeded 20% were removed.

#### DNA palindromes

We define a palindrome as a sequence of DNA followed by its complementary reverse with a sequence of variable length in between (**Fig. 3d**). It is hypothesized that these palindromes can temporarily form DNA hairpins^36^. While in the hairpin state the loop region is singlestranded and open to attack by APOBEC enzymes. Based on observations in breast cancer whole genome sequences^37^, we decided to consider palindromes with a minimal repeat length of 6 bp and an intervening sequence (loop) length of 4-8 bp. We call these regions genome-wide using the algorithm described in Ye et al.^38^ however using our own implementation (<u>https://github.com/TobiasMadsen/detectIR</u>). In total, we find 7.3 M palindrome regions covering a total of 135.2 MB of which 33.6 MB are loop sequence.

#### Computing the false discovery rate

We controlled the false discovery rate (FDR) within each of the sets of tested genomic elements by concatenating all integrated Brown p-values from across all tumor-type cohorts and applying the Benjamini-Hochberg procedure^17^ to the integrated Brown p-values. A q-value threshold of 0.1 was chosen to designate cohort-element combinations as significant hits. In addition, we defined cohort-element combinations in the range 0.1 ≤ q<0.25 as “near signif cance”. We next applied several additional, mutation-based filtering criteria to each significant or near-significant candidate and assigned p-values of 1 to candidates that failed these filtering criteria. Final Benjamini-Hochberg FDR values were then re-calculated on the adjusted sets of integrated Brown p-values to arrive at a list of candidate driver cohort-element combinations.

### 9. Sensitivity and precision analysis of driver predictions (Iñigo Martincorena)

All methods employed in this study were shown to have a low rate of false positives when run on a series of neutral simulated datasets without driver mutations. To evaluate the sensitivity and precision of different methods and particularly of our approach for p-value integration, we compared their relative performance in detecting known cancer genes when applied to protein-coding genes. As a reference gold-standard set of known cancer genes we used a list of 603 genes from the manually-curated Cancer Gene Census v80 database. Results are shown in **Extended Data Fig. 4**.

### 10. Gene expression analyses

(Samir B. Amin, Morten M. Nielsen, Andre Kahles, Nuno Fonseca, Lehmann Kjong, members of the PCAWG Transcriptome Working Group and Jakob Skou Pedersen)

To extend the RNAseq-based expression profiling of GENCODE annotations provided by the PCAWG Transcriptome Working Group^39^, we profiled an extended set of gene annotations, including a comprehensive set of non-coding RNAs (described above and at Synapse:syn5325435).

The profiling used the docker-based workflow described in ^39^ for 1,180 RNA-seq donor libraries, matched to WGS data across 27 different cancer types^39^. In brief, raw sequence reads from donor libraries were uniformly evaluated for QC using FastQC tool, and subsequent alignment was performed on QC-passed libraries using two methods: STAR (v2.4.0i)^40^ and TopHat2 (v2.0.12)^41^. Resulting QC-passed bam files were independently used to quantify extended RNA-seq annotations at the gene-level counts using htseq-count method with following parameters: -m intersection-nonempty --stranded=no —idattr gene_id. This step resulted in two sets of gene-level counts files per donor library which were independently normalized using FPKM normalization and upper quartile normalization (FPKM-UQ). The final expression values were provided as a gene-centric table (rows as genes, columns as samples) with each value representing an average of the TopHat2 and STAR-based alignments FPKM values. Gene-centric tables based on both, GENCODE and extended RNA-seq annotations are available at Synapse data portal: syn2364711. Docker-based workflow for quantifying extended RNA-seq annotations is at <u>https://github.com/dyndna/pcawg14_htseq</u>

### 11. Normalization for copy number variation

(Henrik Tobias Madsen, Morten Muhlig Nielsen, and Jakob Skou Pedersen)

To account for the effects of somatic copy number alterations (SCNAs) on expression, we used two different approaches to create two additional versions of the expression profiles: First a conservative approach where we remove all samples not having the regular bi-allelic copy number for the gene in question. Second a less conservative approach where we build a regression model of expression data based oncopy number (CN) data, and then tested for an effect of somatic mutations on the residual (i.e. the expression that is not explained by copy number).

Generally the higher the copy number of a particular gene the higher expression. The relationship between copy number and gene expression is not strictly linear, as various feedback mechanisms in the cell try to compensate for the mostly deleterious effects of SCNAs. This is known as dosage compensation and has been studied extensively in the context of mammalian sex chromosomes, but also in evolution of yeast and in diseases caused by aneuploidy^42–45^. We therefore fit a linear regression model between the logarithm of expression and the logarithm of CN. This effectively amounts to a power-regression model.

A number of factors makes it hard to learn the regression parameters for each gene and cancer type in isolation. (i) for some cancer types we have only a limited number of samples; (ii) for some genes there is not much variation in CN; and (iii) the variation in expression between samples is generally high. We overcome these problems by employing a mixed model strategy, that allows sharing of information between genes, effectively regularizing the parameter estimates for gene cancer-type combinations that carry little information on their own.

Let^FPKM^_g,c,i_ and^SCNA^_g,c,i_ denote the expression and SCNA measurement respectively for gene, *g*, in cancer-type, *c*, and sample *i* respectively. We then define:

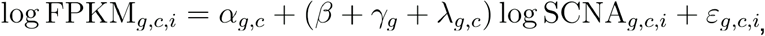

where ^α^_g,c_ and *β* are fixed effects, whereas *γ_g_* and *^λ^_g,c_* are random effects, with 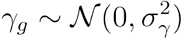 and 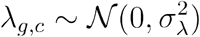 Finally the residual is 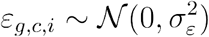.

Using this model we infer a global SCNA to expression regression, *β*, but allow some regularized gene-specific and gene/cancer type-specific variation: *γ_g_* and *λ_g,c_*

Thus we exploit the similarity across genes and similarity within genes across cancer types.

Since the variance increases with the absolute value of the explanatory variable associated with a random slope, this kind of mixed model display heteroskedasticity. Furthermore the model is not invariant under scaling of the explanatory variable, in this case SCNA. We centralise l°g SCNA, such that normal diploid regions have the least variance.

### 12. Mutation to expression association

(Morten Muhlig Nielsen, Henrik Tobias Madsen, and Jakob Skou Pedersen)

Mutation to expression association was calculated using non-parametric rank sum based statistics on z-score normalized expression values. This equalizes expression mean and variance for each cancer type and is a way to allow for comparison across cancer types. For comparisons of expression in mutated vs non-mutated (wild-type) cases within the same cancer type, the test reduces to the Wilcoxon rank sum test. For comparisons involving samples in multiple cancer types, such as meta-cohorts and pan-cancer cohorts, the statistic and associated p-value is still the non-parametric Wilcoxon rank sum test, and thus no assumptions regarding distribution of z-score expressions are made. Tied expression values were broken by adding a small random rank robust value. Association estimates were performed based on original expression values as well as the two copy-number normalized expression sets mentioned above. Fold difference values were calculated per mutation as the log_2_ ratio of the expression of the mutated tumor to the median of all wild-type tumors of the same cancer type. Reported Fold Difference values for an element with multiple mutations represent the median fold difference of all mutations in that element.

### 13. Copy number analyses

(Esther Rheinbay)

We surveyed significant focal copy number alterations for candidate driver genes as orthogonal evidence for their “driverness”. Significant copy number alterations were obtained from the TCGA Copy Number Portal (<u>http://portals.broadinstitute.org/tcga/home</u>), analysis 2015-06-01-stddata-2015_04_02 regular peel-off, a database of recurrent copy number alterations calculated by the GISTIC2 algorithm^46^ across >10,000 samples and 33 tumor types from TCGA. GISTIC2 results were included for candidate drivers if a gene was significant (residual q<0.1) and was located within a peak with ≤ 10 genes. Visualization was performed with the Integrative Genomics Viewer (IGV)^47^.

### 14. Power Calculations

(Esther Rheinbay)

#### Calculation of mutation detection sensitivity

Average detection sensitivity, the power to detect a true somatic variant, was calculated using a binomial model across all exon-like regions for different genomic element types as previously published^27, 48^. Detection sensitivity was based on sequencing coverage and estimated clonal variant allele fraction from 60 representative PCAWG alignments from the pilot cases (<u>https://doi.org/10.1101/161562</u>; **Supplementary Table 7**). Clonal allele fraction was estimated based on PCAWG consensus purity and ploidy estimates (doi: https://doi.org/10.1101/161562) as purity/(average ploidy)^48^. Element-wise averages were calculated as average across all exons for a given element.

#### Estimation of total number of promoter hotspot mutations

Detection sensitivity for all patients was calculated for the two most recurrent *TERT* promoter hotspot sites (chr5:1295228, chr5:1295250; hg19) using total read depth at these positions, sample purity and average ploidy. For each cohort, the number and percentage of powered (≥90%) patients was obtained. The number of total expected mutations was then inferred as number of observed (called) mutations divided by the fraction of patients powered. The number of “missed” mutations is the difference between the total expected and observed mutations. Percentages of these numbers were calculated relative to the size of individual patient cohorts. Confidence intervals (95%) on the total percentage of patients with a *TERT* hotspot mutation were calculated using the beta distribution. Poisson confidence intervals were calculated for the number of missed mutations in the PCAWG cohort. Note that the inference of *TERT* mutations assumes exactly one mutation per patient. Estimates for the *FOXA1* promoter hotspot mutation (chr14:38064406; hg19) were conducted using the same procedure.

#### Calculation of the minimum powered mutation frequency in a population

Power to discover driver elements mutated at a certain frequency in the population were conducted as described before^21, 49^, but solving for the lowest frequency for a driver element in the patient population that is powered (≥90%) for discovery. The calculation of this lowest frequency takes into account the average background mutation frequencies for each cohort/element combination, the median length and average detection sensitivity for each element type, patient cohort size, and a global desired false positive rate of 10%.

### 15. Associations between mutation and signatures of selection: loss of heterozygosity and cancer allelic fractions

(Federico Abascal, Iñigo Martincorena)

For protein-coding sequences mutation recurrence can be analysed in the context of the functional impact of mutations (e.g. missense, truncating) to better distinguish the signal of selection. In contrast, estimating the functional impact of mutations in non-coding elements of the genome is a difficult, yet unsolved problem. To overcome this limitation and be able to compare selection signatures for both coding and non-coding elements under a similar framework, we developed two measures of selection which are agnostic to the functional impact of mutations.

#### Association between mutation and loss of heterozygosity

When a tumour carries a driver mutation in one allele of a given gene, it may be the case that a second hit on the other allele confers a growth advantage and is positively selected. When one of the events involves the loss of one of the alleles the process is referred to as loss of heterozygosity (LOH). This kind of biallelic losses are typical of, but not exclusive to, tumour suppressor genes (TSGs).

For each gene, we build a 2x2 contingency table indicating the number of cases in which the gene was mutated or not and the number of cases in which the gene was subject to LOH or not. We applied a Fisher’s exact test of proportions to identify which genes showed an excess of LOH associated to mutation. P-values were corrected with the FDR method to account for multiple hypotheses testing. This analysis was applied to each tumour type and cohort separately and proved very successful in identifying TSG as well as some oncogenes (OGs) as well.

#### Association between mutation and cancer allelic fractions

Driver mutations that provide an advantage for tumour cells are expected to show higher allelic fractions based on different interacting processes, including: early selection; amplification of the locus carrying the driver mutation; loss of the non-mutated locus (LOH). Comparing cancer allelic fractions (CAF) can be informative to detect signatures of selection, both for TSGs and OGs.

CAFs are defined here as the proportion of reads coming from the tumour and carrying the mutation. To transform observed fractions (VAFs) into CAFs, tumour purity and local ploidy needs to be taken into account according to the following formula:

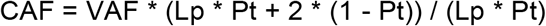

Where *Lp* corresponds to the local ploidy for the mutated locus, and *Pt* denotes the tumour purity. Ploidy and tumour purity predictions were obtained from Dentro et al (in preparation).

To determine whether CAFs for a given gene or element were higher than expected we compared them to the CAFs observed in flanking regions. To define flanking regions we took 2 kb at each side of the gene/element, excluding any eventually overlapping coding exon, and also included introns (if present). The two sets of CAFs associated to each gene/element, i.e. those CAFs lying within the gene and those flanking it, were compared with a t-test to detect significant deviations. P-values were corrected with the FDR method. This approach was able to identify most known TSGs and OGs.

### 16. Signals of selection in aggregates of non-coding regions of known cancer genes

(Federico Abascal, Iñigo Martincorena)

We conducted a series of analyses on regions combined across genes to determine whether the paucity of driver mutations found in non-coding regions was related to lack of statistical power in single-gene analyses. This analysis also aimed to estimate how many driver mutations present in cancer genes were missed in this study. For protein-coding sequences, the number of driver mutations was estimated using dN/dS ratios as described in (^50^). For non-coding, regulatory regions of protein-coding genes (Promoter and UTRs), we relied on a modified version of the NBR negative binomial regression model described above (Section 5) to quantify the overall excess of driver mutations. We applied a second approach to determine whether there was an enrichment of LOH associated to mutations in the different types of non-coding regions associated to protein-coding genes.

#### Observed vs. expected numbers of mutations based on the NBR mutation model

NBR was used to estimate the background mutation rate expected across cancer genes, using a conservative list of 19,082 putative passenger genes as background. The resulting model is used to predict the numbers of expected SNVs and indels per element type per gene, and aggregate sums across genes. For this analysis we selected a reduced but diverse set of 142 cancer genes, encompassing 27 focally amplified and 22 deleted genes found by TCGA and present in the Cancer Gene Census^51^, and 112 genes with a significant signal of selection in this study (q<0.1) and present in the Cancer Gene Census. The list of 142 genes can be found in **Supplementary. Table 7**. To be as accurate as possible, we used a diverse set of covariates in the NBR model, including: local mutation rate (estimated on neutral regions +/-100 kb around each gene), detection sensitivity (defined as the element-average proportion of callable samples per site according to MuTect), gene expression covariates (first 8 principal components of the matrix of average gene expression values in each tumour type, as well as two binary variables marking the 500 genes with highest expression values in any tumour and 1,229 genes with a maximum FPKM lower than 0.1 across tumour types), and averaged copy-number calls for each gene across all samples (see **Supplementary Note** for more details). For each element type, the sum of observed mutations across the 142 cancer genes were compared to the sum of the expected rates to estimate the excess of mutations in regulatory and coding regions of cancer genes. An excess of observed mutations provides an estimate of the number of driver events^18^. Confidence intervals were calculated using the equation for the ratio of two Poisson observations, which are the number of mutations in the list of known cancer genes and in the list of passenger genes. It is important to know that these confidence intervals do not capture uncertainty in the background model and should be interpreted with caution. For this reason, we systematically evaluated the impact of a diverse array of covariates on our estimates (see Supplementary Note). We also note that this test can underestimate the number of non-coding drivers since some driver mutations can be present in the list of putative passenger genes, although this effect is expected to be quantitatively small if the density of driver mutations in regulatory regions of known cancer genes is higher than in those of putative passenger genes.

#### Mutation-LOH association for aggregates of genes

For this analysis we combined data across known cancer genes, including 603 genes in the Cancer Gene Census v80 and 154 additional significantly mutated genes found by exome studies^18, 21^. To estimate whether there was an excess of LOH associated to mutation in regulatory and coding regions of cancer genes, we calculated the fold change in LOH for the aggregate of cancer genes and normalized it dividing by the fold change observed in passenger genes. Confidence intervals were estimated using parametric bootstrapping (100,000 pseudoreplicates) for both cancer and passenger genes.

### 17. Mutational process and indel enrichment

(Federico Abascal, Iñigo Martincorena)

After noticing the skewed distribution of indel lengths in genes like *ALB, SFTBP, NEAT1* and *MALAT1*, we carried a search for other genes showing the same pattern, which may be subject to the same mutational process. For every gene we record the proportion of indels of length 2-5 bp out of the total number of indels and compared this proportion with the background proportion using a binomial test. The background proportion was calculated using all protein-coding and lncRNAs genes. For every gene we also calculated the indel rate and compared it to the background indel rate using a binomial test. Both sets of p-values were independently corrected with the FDR method. The analysis was done for each tumour type separately. Genes with a q-value < 0.1 both for enrichment in 2-5 bp indels and for higher indel rates were further analyzed as candidates to be under the process of localized indel hypermutation described in this study. The levels of expression of these genes were analysed across all tumour types.

### 18. Mutations association to splicing (Andre Kahles)

For the assessment of the relationship between mutations in the U2 locus upstream of *WDR74* and local changes in alternative splicing, we analysed changes in the percent-spliced-in (PSI) value^52^ of alternative splicing events located in *WDR74* relative to the given genotype. The analysis was based on four different alternative splicing event types extracted and quantified by the PCAWG Transcriptome Working Group (exon skip, intron retention, alternative 3’ splice site, alternative 5’ splice site)^39^. In total we analysed 116, 103, 27 and 98 events, respectively, for the above event types. We then filtered the events for a minimum of expression evidence (non-NaN PSI) in at least 10 samples, presence of the alternate allele together with expression evidence in at least 5 samples, and a minimum absolute distance of mean PSI values of alternate and reference group of 0.05. Based on the selected events, we retained 3, 3, 1 and 0 events for analysis. Using an ordinary least squares model with the genotypes as factors and the the PSI values as responses, we computed p-values and the variance explained by presence of mutations. Multiple testing correction followed the Bonferroni method.

To assess the relationship between mutations in the U2 locus upstream of *WDR74* and global changes in splicing, we used two global splicing statistics. We measured the amount of splicing as the number of edges in the splicing graph of a gene in a given samples. All splicing graphs were taken from the analyses of the PCAWG Transcriptome Working Group^39^. For each sample, we computed the mean number of splicing edges over all genes, resulting in the extent of splicing per sample. We measured the extent of splicing as how far each event is outlying from the mean over all event in the same cancer type (encoded by project code x histotype). The splicing outlier values per sample and gene were taken from the PCAWG Transcriptome Working Group analyses^39^. The mean over all genes of a sample was taken as a statistic for the extent of splicing.

For both statistics, we again used an ordinary least squares model as described above to model the relationship between genotype and splicing. As splicing is highly tissue dependent, we included the project codes as additional factors, accounting for both tissue specificity as well as possible underlying batch effects. To extract the amount of additional variance explained through the presence of mutations, we computed one model on project codes and presence of mutations and one model on project codes alone. For the model, we only included samples of project codes, where we had at least one sample with a mutation present, resulting in a total of 618 samples over 14 project codes.

### 19. Structural variation analysis (Morten Muhlig Nielsen, Lars Feuerbach)

Structural variant data was provided by the PCAWG Structural Variation Working Group. The data provide p-values for the observed breakpoint counts in 50kb bins along the genome. Candidate elements were overlapped with the bins, and fisher’s method was used to calculate a single p-value for each element. The set of element p-values were corrected with the FDR method.

### 20. RNA structural analysis

(Radhakrishnan Sabarinathan, Ciyue Shen, Chris Sander, Jakob Skou Pedersen)

In order to test if the observed mutations (SNVs) in the RMRP gene are biased towards high RNA secondary structure impact, we performed a permutation test by following the steps used in oncodriveFML^24^ together with the predicted structural impact scores from RNAsnp^53^. At first, the RNAsnp was run with the options -m 1 -w 300 and other default parameters to obtain the minimum correlation coefficient (r_min) score for each possible mutations in the RMRP gene. The r_min scores were then transformed, 1-((r_min+1)/2), to range between 0 and 1, where 1 indicates high structural impact score. Further, we followed the steps of oncodriveFML (see section ‘oncodriveFML’ for more details) with 1,000,000 randomizations and by using per sample mutational signatures (i.e., the probability of observing a mutation in a particular tri-nucleotide context in a given sample) to compute the p-value at the cohort and sample level.

Furthermore, the RNA secondary structure impact scores (r_min) of indels (insertions/deletion) were computed by using a modified version of RNAsnp (since the current version of RNAsnp is limited to substitutions only). Briefly, we first computed the base pair probability matrices of wild-type and mutant sequences (by taking into account the insertion or deletion) and then adjusted the size of matrices to be equal (by introducing additional rows and columns with zeros in one of the matrices with respect to insertion or deletion). Further, by following the steps of RNAsnp, we computed the r_min score. The structure shown in **Fig. 6d** is based on the conserved secondary structure annotation obtained from Rfam (RF00030)^54^.

Tertiary structure contacts in RMRP were predicted using evolutionary couplings co-variation analysis (EC analysis ^55^) of the multiple sequence alignment of 933 eukaryotic *RMRP* sequences from Rfam (RF00030). The EC analysis (software available at https://github.com/debbiemarkslab/plmc) was run with the options -le 20.0, -lh 0.01, -t 0.2, -m 100 and the top 100 interactions were chosen as predicted contacts, either in secondary or tertiary structure, depending on local context. As no experimental 3D structure or crosslinking experiments of the mammalian RMRP are available, interaction sites were inferred by homology to the partially known yeast RMRP crystal structure. We (1) aligned the human RMRP sequence with the Saccharomyces cerevisiae RMRP sequence using the sequence family covariance model from Rfam and (2) mapped the locations of RNA-protein interactions within 4Å^56^ from the crystal structure and the experimentally determined RNA-protein crosslinking sites^57^, and RNA substrate crosslinking sites^58^ from the yeast sequence to the human RMRP sequence. For the crosslinking sites, a ±3 nucleotide window is reported as the interaction site. In order to test if the locations of the observed indels are biased towards tertiary structure, protein- or substrate-interaction sites, 1,000,000 randomizations of five indels were performed assuming uniform distribution of indels across the RMRP gene body, and an empirical p-value was calculated (P = .08), showing a potential for functional impact.

Two different overlapping deletion calls in the *RMRP* gene body were observed in the same thyroid cancer patient. After manual inspection of the tumor and normal bam files, it was found that these calls were based on the same mutational event.

### 21. Cancer associated germline variant distance to non-coding driver candidates

(Morten Muhlig Nielsen)

We used a set of genome-wide significant cancer associated germline SNPs (n=650) from the NHGRI-EBI GWAS catalog^59^ as collected by Sud et al^60^. We evaluated the genomic distance from candidate non-coding drivers to the closest germline variant. All distances were above 50 kb with the exception of the *TERT* promoter which was 1 kb away from a coding variant (rs2736098) in the *TERT* gene.

## Supplementary Note

1. Overview of non-coding hotspots in top 50 not categorized in main text
2. Discussion of additional significant non-coding elements
3. Evaluation of splicing association of *U2* mutations upstream of *WDR74*
4. Enrichment of protein-coding drivers
5. Impact of covariates on the estimation of driver mutations in functional regions of cancer genes
6. Author contributions by category
7. References

### 1. Overview of non-coding hotspots in top 50 not categorized in main text

Six of the 35 non-coding hotspots in top 50 are not assigned to the mutational processes described in the main text and highlighted in **Fig. 1b**. Below we describe these in detail.

Four of the six remaining non-coding hotspots were found on the X chromosome (X: 116579329, X:7791111, X:83966025, and X:83967552). The mutations in these hotspots were mainly contributed by males (61-94% of mutations per hotspot), suggesting the possibility of uncaught noise. One of these hotspots is located in palindromic DNA (X:7791111). The DNA sequence around this position is composed of two mononucleotide repeats of pairing bases (ATTTTTAAAAAAAAAT), which fits our definition of palindromic structures (**Methods**). The sequence context around this hotspot do not match the sequence recognized by APOBEC enzymes^1^ and the hotspot had a low APOBEC signature probability (**Fig. 1b**). It is therefore unlikely to be caused by the same mutational processes as the other palindromic hotspots in the top 50 hotspots, but rather likely represents noise associated with homopolymer runs (PCAWG variants paper).

A hotspot (3:164903710) contained mutations in multiple cancer types with most mutations contributed by Liver-HCC (6/17). It is located about 1 kb downstream of ***SLITRK3,*** two base pairs from an annotated CTCF transcription factor binding site. CTCF sites and the base pairs immediately downstream are known to be highly mutated in several cancer types including Liver-HCC. This position is lowly conserved, suggesting that it would not have a functional impact^2^.

The last non-coding hotspot (1:103599442) had a high proportion of mutations attributed to the COSMIC5 signature. It overlaps a repetitive LINE region, suggesting potential mapping issues. Moreover, the position is flanked by two other positions, two base pairs away in both directions, which also have mutations in the same samples and reads, however these positions were called problematic in the Panel-of-Normals (PoN) filter, further suggesting mapping issues.

### 2. Discussion of additional significant non-coding elements

#### *WDR74* promoter

The *WDR74* promoter has already been suggested as a potential driver in several studies^3–6^. However, we found that mutations are concentrated inside a *U2* RNA where the density of putative polymorphisms is abnormally high. This outstanding level of diversity could in principle be due to higher mutability in the germline. However, given the repetitive nature of the *U2* element (there are hundreds of copies in the genome) and the extreme levels of putative diversity, it is more likely that this region is a source of mapping artefacts. This observation raise concerns about the validity of *WDR74* promoter and, hence, on its potential as a cancer driver (**Extended Data Fig. 10**).

#### *TMEM107* 3’UTR

The case of the *TMEM107* 3’UTR is very similar to that of the *WDR74* promoter. There is a small RNA *(SNORD118)* embedded in the 3’UTR, and is there where most mutations concentrate. In PCAWG normals data there is a remarkable excess of putative SNP diversity in this element. (**Extended Data Fig. 10**).

#### Additional non-coding RNAs hits

Eleven additional ncRNAs or ncRNA promoter hits, not described in detail in the main text, passed the flagging of potential false positive hits *(RNU6-573P, RPPH1, RNU12, TRAM2-AS1, G025135, G029190,RP11-92C4.6* and promoters of *RNU12, MIR663A, RP11-440L14.1,* and *LINC00963).* The driver role of these were generally not supported by additional lines of evidence and lacked functional evidence or appeared to be affected by technical artefacts. They are described individually below.

#### RNU6-573P

The small RNA *RNU6-573P* is detected in Endocrine pancreas (q = 7.3x10^−4^) with three mutations. However, it appears to be a nonfunctional pseudogene recently inserted in the human lineage. Not only the locus, but the wider region is subject to increased mutational burden, further supporting that mutational mechanisms or technical issues rather than selection underlies the mutational recurrence.

#### RNU12

*RNU12* is a spliceosomal RNA, which was found significant in Lymph-BNHL (q = 5.0x10^−2^) and in the Hematopoietic system meta cohort (q = 7.0x10^−2^). The mutation rate is 1.6 times higher inside the gene in pan-cancer compared to the flanking regions. However, the mutation rate is 4 times higher in gnomAD, which renders it as a likely problematic region. Significance in the promoter of *RNU12* was also identified in Lymphomas (q = 3.5x10^−2^) and the Hematopoietic system meta cohort (q = 1.8x10^−6^). But this region is overlapping the promoter of *POLDIP3* (Polymerase delta-interacting protein 3), which makes interpretation difficult.

#### MIR663A

The promoter of *MIR663A* was recurrently mutated in the carcinoma meta cohort (q = 7.2x10^−4^). It is the primary transcript of *MiR-663,* which has tumorigenic functions in gastric cancer and nasopharyngeal carcinoma^7^. The mutation rate is 1.2 times higher inside the element compared to the flanking regions, but the mutation rate is 2.4 times higher in gnomAD. The mutations were also not correlated with expression.

#### RPPH1

*RPPH1* forms the RNA component of the RNase P ribonucleoprotein, which matures precursor-tRNAs by cleaving their 5’ end^8^. It is transcribed from a divergent promoter together with the protein-coding gene *PARP2,* however, no association with expression was observed for either gene. The region has a high level of germline polymorphisms in normal samples indicative of a problematic region to map.

#### TRAM2-AS1

The promoter element of the antisense non-coding RNA *TRAM2-AS1* is recurrently mutated in the Female reproductive system meta cohort (q = 0.10). The promoter is shared with the divergently transcribed protein-coding gene *TRAM2.* Interestingly, the promoter has 8 mutations in other cohorts, which collectively associate with the expression of *TRAM2* (*P* = 0. 03; carcinoma) but not *TRAM2-AS1* (*P* = 0.9). It is uncertain which gene is controlled by this significant promoter element.

#### G025135

*G025135* is a ncRNA element from MiTranscriptome^9^ identified here as a significant driver candidate in Lymph-CLL (q = 0.01). There are patients with many mutations, indicating that an unknown mutation process is at play leading to false driver prediction.

#### G029190

The *G029190* ncRNA is also from MiTranscriptome with significance in Kidney-RCC (q = 0. 04) and in the Kidney meta cohort (q = 0.05). It is located downstream of *RAB11FIP3* and upstream of *CAPN15.* A possible function of this gene has not been established.

#### LINC00963

*LINC00963* was found significant in the Kidney meta cohort (q = 0.04). It has many alternative transcripts with different TSSs and thus a large promoter. *LINC00963* is known to be involved in the prostate cancer transition from androgen-dependent to androgen-independent and metastasis via the EGFR signaling pathway^10^. The SNV mutation rate is generally high (0.05 mutations per position), although lower than the flanking regions (0.08).

#### RP11-92C4.6

*RP11-92C4.6* is a predicted antisense ncRNA, which was identified as a significant driver candidate in the Breast meta cohort (q = 0.08). It is a short region with low conservation containing four SNVs. It is overlapping the *COL15A1* protein-coding gene with the upstream promoter region. *COL15A1* has previously been linked to ovarian cancer. It is unclear whether this ncRNA annotation is valid and whether these reflect true functional mutations.

#### RP11-440L14.1

The promoter of *RP11-440L14.1* was identified as a candidate driver in the Carcinoma meta cohort with 14 mutations (q = 5.4x10^−3^). It has a hotspot position with four mutations overlapping two different deletions. The lncRNA is located between *CPLX1* and *PCGF3.* The validity of the annotation and possible function for this lncRNA has not been established.

### 3. Evaluation of splicing association of *U2* mutations upstream of *WDR74*

We hypothesized that the mutations observed in the evolutionarily conserved spliceosomal *U2* RNA upstream of *WDR74* may affect splicing. As prior sequencing evidence, summarised in the GENCODE version 19 gene annotations^11^, shows that the *U2* is cotranscribed as an alternative 5’ exon in some cases, it might play a role in splicing and regulation of the *WDR74* transcript. We therefore both evaluated (A) whether the mutations associate with alternative splicing of the *WDR74* gene; and (B) whether they associate with transcriptome-wide changes in splicing.

A) We modelled the association of the the somatic mutations with the amount of alternative splicing, as measured by percent-spliced-in values^12^, using ordinary least squares regression, which did not reveal any significant associations. For none of the seven tested alternative splicing events (three exon skips, three intron retentions, one alternative 3-prime splice site) the genotype was able to explain a substantial fraction of the observed variation. The largest R-squared value was 0.01 and no p-value reached nominal significance.

B) A similar result was obtained for the relationship between the presence of *U2* mutations and global changes in alternative splicing. Modeling the amount of alternative splicing on the ICGC project codes alone, which reflect cancer types and contributing institutions, we reached an R-squared value of 0.612, reflecting the strong relationship between splicing and tissue identity. Including the presence of somatic mutations in the model did not change this value. We obtained an analogous result for the extent of alternative splicing, reaching an R-squared of 0.414 on the project codes alone and the same value when including the genotype into the model.

In conclusion, we found no evidence for an association between mutations in the *U2* upstream of *WDR74* and alternative splicing of *WDR74* transcripts or global changes in splicing.

### 4. Enrichment of protein-coding drivers

Collectively, mutations occurring in the promoter region of the 757 cancer genes did not have a significantly different association with expression than synonymous mutations (**Supplementary fig. 1a**). Similarly, promoter and UTR mutations in cancer genes are not significantly enriched in LOH with respect to mutations in putative passenger genes (**Supplementary fig. 1b**). This is consistent with the prediction above that only a very small fraction of the mutations observed in the promoters and UTRs of known cancer genes are genuine driver events.

**Supplementary figure 1:**
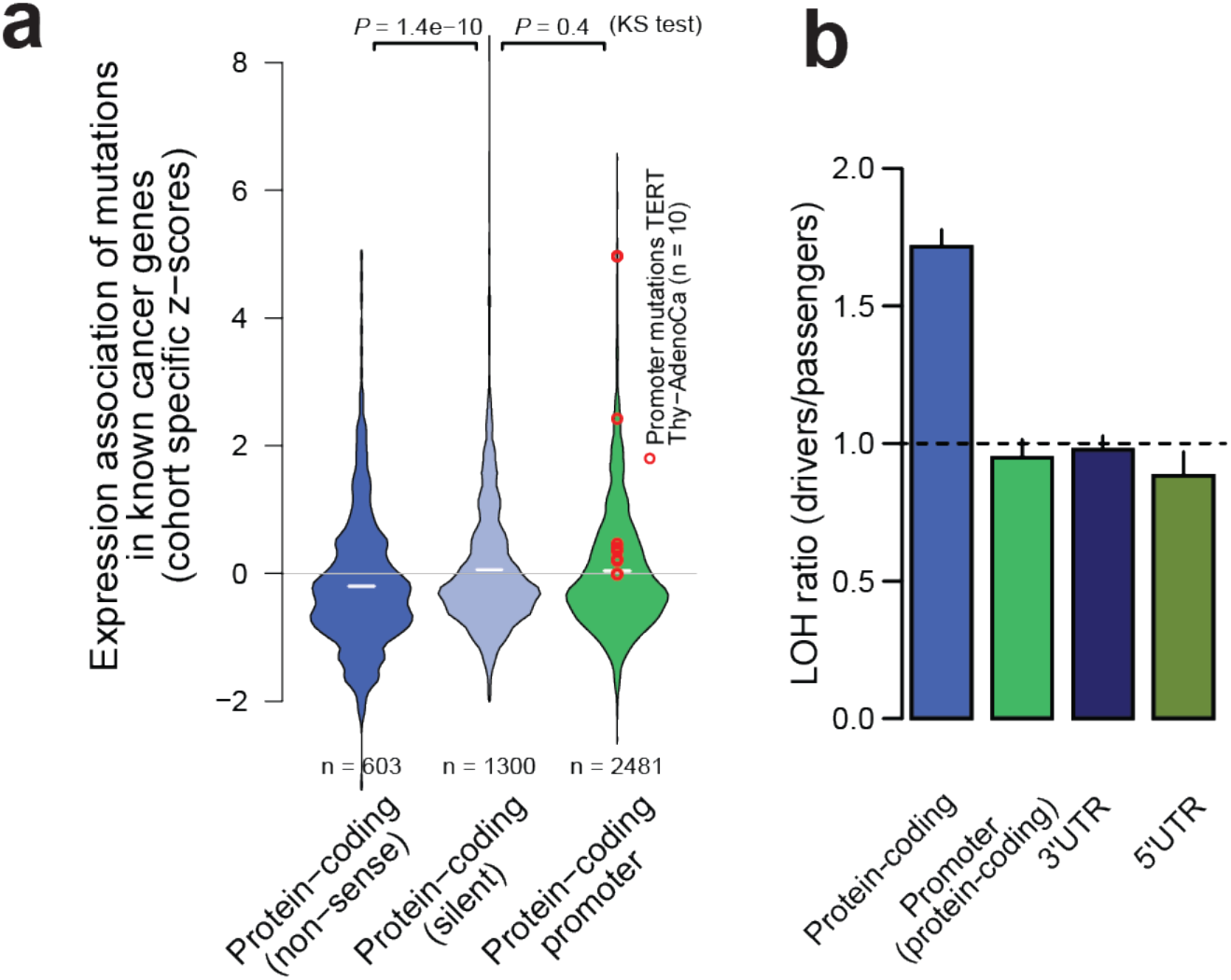
**a**, Expression associated with mutations in coding and promoter regions of cancer genes. Z-score expressions associated with non-sense mutations deviate significantly from silent mutations, likely through nonsense mediated decay, whereas expressions associated with promoter mutations do not differ from that of silent mutations. Only mutations in diploid positions were used. **b**, Excess of LOH associated to mutation in regulatory and coding regions of cancer genes. The y-axes shows the ratio of fold changes in cancer vs. passenger genes, with fold changes representing the excess or depletion of LOH associated with mutation.

### 5. Impact of covariates on the estimation of driver mutations in functional regions of cancer genes

As described in **Methods**, we estimated the abundance of driver mutations in coding and regulatory regions (promoter and UTRs) of 142 known cancer genes using the NBR background model fitted on putative passenger genes.

Different covariates were included to improve the fit of the model. The local mutation rate, calculated on neutral regions within +/-100 kb around each element, was included to account for regional variation of mutation rates. Detection sensitivity (d.s.) was averaged for each element using the MuTect estimates of callable sites from each sample. For genes in chromosomes X and Y, we did not have MuTect estimates and d.s. was imputed using a linear regression model of d.s. as a function of GC content. Detection sensitivity accounts for elements where the mutation rate is lower than expected just because of poor sequencing coverage. A third type of covariate was included in the model to account for associations between gene expression levels and mutation rates. Starting with a matrix of mean FPKM expression values for each gene and tumour type, we log-transformed and scaled the expression matrix using pseudocounts and applied Principal Component Analysis to reduce the dimensionality. We selected the first 8 components as covariates, which together explained 95.5% of the variance. In addition, we added two additional covariates to account for non-linearity between expression and mutation rate in the tails of the expression spectrum. To accomplish this, we created two binary variables, one marking the 500 genes with highest maximum expression values across tumour types, the other marking 1,229 genes whose expression did not exceed FPKM values of 0.1 in any tumour type. Finally, since tumours are rich in amplifications and deletions and these events may result in seemingly increased or decreased mutation rates, we included a copy-number covariate, calculated as the average copy number of each gene across all PCAWG samples.

We intentionally selected a small set of cancer genes to estimate the abundance of driver mutations because with larger numbers of genes the signal of selection becomes weaker while any systematic bias may become more prominent. We noticed these biases becoming increasingly apparent when analyzing larger sets of genes. In an attempt to capture a diverse but relatively small group of cancer genes, we included Cancer Gene Census genes recurrently mutated by coding mutations in PCAWG and genes frequently altered by copy number gains or losses, as described in Methods.

**Supplementary Table 9** shows the impact of using different covariates on the 142 genes selected for this analysis. Reassuringly, this shows that the estimates are broadly consistent across models with different covariates, with variations typically within the confidence intervals of alternative models. This confirms that the overall conclusions are largely unaffected by the use of different models. **Supplementary Figure 2** shows the results for the 142 genes and for a larger set of 603 genes from the Cancer Gene Census.

To evaluate the performance of the NBR model, we compared the number of driver substitutions predicted by NBR in the CDS regions of the 142 cancer genes to the number predicted by dN/dS (calculated by dNdScv). dN/dS offers an independent estimate of the number of driver substitutions in a group of genes using the local density of synonymous mutations to estimate the neutral expectation^13^, instead of predicting the background mutation rate by extrapolation from putative passenger genes using a regression model. Reassuringly, in these 142 genes, NBR predicts 2,258 (CI_95_%: 2,127-2,382) driver substitutions and dN/dS predicts 2,384 (CI_95_%: 2,043-2,759).

**Supplementary figure 2:**
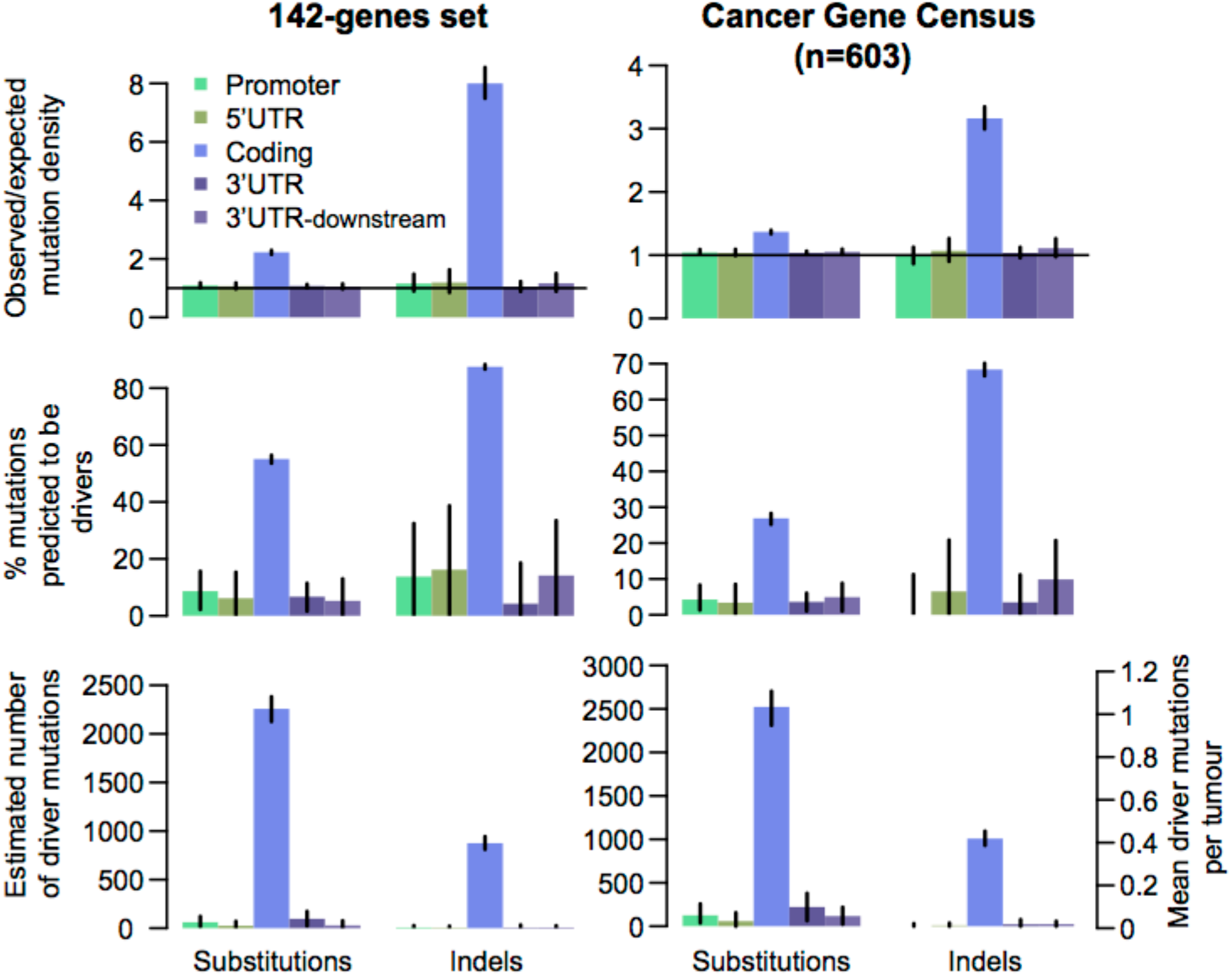
Estimation of the excess substitutions (left) and indels (right) in regulatory and protein-coding regions of cancer genes and for the 142 (left) genes and a larger set of genes from the Cancer Gene Census (right). Ratios of observed vs. expected number of mutations (top), the percentage of mutations predicted to be drivers (middle), and the total number of predicted drivers in all cancers and in each patient (bottom) are shown.

### 6. Author contributions by category

#### Driver discovery

A.L., C.H., C.W., D.A.W., E.K., E.M.L., E.R., G.G., G.T., H.M.U., I.M., J.K., J.R., J.S.P., K.A.B., K.D., K.I., L.M., L.U.-R., M.M.N., M.P.H., N.A.S., P.D., P.J.C., R.J., S.B.A., T.A.J., T.T. and, Y.F. contributed and curated genomic annotations. C.H., C.W.Y.C., I.M., S.B.A. and Y.E.M. contributed randomized mutational data sets for driver discovery. A.L., A.G.-P., A.H., D.L., D.T., E.K., E.M.L., E.R., H.H., H.M., I.M., I.R., J.B., J.C.-F., J.D., J.F., J.M.H., J.R., J.Z., K.C., K.D., K.I., L.L., L.M., L.S., L.U.-R., L.W., M.B.G., M.J., N.L.-B., O.P., P.D., Q.G., R.S., S.K. and S.S. contributed driver methods and results. E.R., G.G. and G.T. implemented results integration. A.K., C.v.M., C.V., G.T., H.H., I.M., J.R., L.F. and M.M.N. contributed driver results integration. C.H., C.W.Y.C., E.K., E.R., G.G., J.K., J.M.H., J.S.P., M.M.N. and R.I.P. contributed single site recurrence analysis.

#### Candidate vetting and filtering

E.R., F.A., H.H., H.T.M., J.K., L.F. and M.M.N. contributed individual candidate filters. A.L., C.H., C.W., E.K., E.M.L., E.R., F.A., G.G., G.T., H.H., H.M., H.M.U., J.K., J.M.H., J.S.P., K.D., L.F., L.S., M.M.N., M.S.L., N.A.S. and R.J. performed candidate vetting.

#### Case-based analysis

E.R., F.A., G.G., H.H., I.M., J.M.H., J.R., J.S.P., K.P., M.M.N. and M.P.H. contributed case-based analysis. A.G.-P., A.H., A.L., C.H., D.C., D.T., E.K., E.R., F.A., G.G., G.T., H.H., H.K., I. M., J.C.-F, J.R., J.S.P., K.I., K.P., L.M., L.S., L.U.-R., L.W., M.A.R., M.B.G., M.M.N., M.S.L., N.A.S., N.L.-B., O.P., R.I.P., R.S., S.K. and Y.K. contributed results interpretation. A.K., J.S.P., K.A.B., K.-V.L., M.M.N., N.A.F., S.B.A., T.A.J. and T.T. contributed expression profiling (extended GENCODE set). A.K., C.V., D.C., H.M., H.T.M., J.R., J.S.P., K.I., L.W., M.A.R., M.M.N., M.S.L. and S.B.A. contributed mutation-to-expression correlation analysis. A.K., C.V., D.C., J.R., L.W., M.A.R., N.A.S. and Z.Z. contributed network or pathway analysis. R.S, Ciyue S., Chris S., and J.S.P. contributed structural RNA analysis.

#### Power analysis and driver mutations at known cancer genes

E.R. analysed SNV detection and driver discovery power. I.M. evaluated excess number of mutations at known drivers. F.A. and M.M.N. integrated additional evidence.

#### Leadership and organizational work

E.R., G.G. and J.S.P. contributed working group leadership. A.G.-P., D.A.W., E.K., E.R., G.G., G.T., I.M., J.R., J.S.P., L.F., L.M., M.B.G., N.L.-B., O.P., P.J.C., R.S. and S.K. contributed organization.

## References

1. Khurana, E. et al. Integrative annotation of variants from 1092 humans: application to cancer genomics. Science 342, 1235587 (2013).

2. Fredriksson, N. J., Ny, L., Nilsson, J. A. & Larsson, E. Systematic analysis of noncoding somatic mutations and gene expression alterations across 14 tumor types. Nat. Genet. 46, 1258–1263 (2014).

3. Nik-Zainal, S. et al. Landscape of somatic mutations in 560 breast cancer whole-genome sequences. Nature 534, 47–54 (2016).

4. Melton, C., Reuter, J. A., Spacek, D. V. & Snyder, M. Recurrent somatic mutations in regulatory regions of human cancer genomes. Nat. Genet. 47, 710–716 (2015).

5. Puente, X. S. et al. Non-coding recurrent mutations in chronic lymphocytic leukaemia. Nature 526, 519–524 (2015).

6. Northcott, P. A. et al. Enhancer hijacking activates GFI1 family oncogenes in medulloblastoma. Nature 511, 428–434 (2014).

7. Weinhold, N., Jacobsen, A., Schultz, N., Sander, C. & Lee, W. Genome-wide analysis of noncoding regulatory mutations in cancer. Nat. Genet. 46, 1160–1165 (2014).

8. Imielinski, M., Guo, G. & Meyerson, M. Insertions and Deletions Target Lineage-Defining Genes in Human Cancers. Cell 168, 460–472.e14 (2017).

9. Rheinbay, E. et al. Recurrent and functional regulatory mutations in breast cancer. Nature (2017). doi:10.1038/nature22992

10. Huang, F. W. et al. Highly recurrent TERT promoter mutations in human melanoma. Science 339, 957–959 (2013).

11. Horn, S. et al. TERT promoter mutations in familial and sporadic melanoma. Science 339, 959–961 (2013).

12. Flavahan, W. A. et al. Insulator dysfunction and oncogene activation in IDH mutant gliomas. Nature 529, 110–114 (2016).

13. Huarte, M. The emerging role of lncRNAs in cancer. Nat. Med. 21, 1253–1261 (2015).

14. Bhan, A., Soleimani, M. & Mandal, S. S. Long Noncoding RNA and Cancer: A New Paradigm. Cancer Res. 77, 3965–3981 (2017).

15. Lanzós, A. et al. Discovery of Cancer Driver Long Noncoding RNAs across 1112 Tumour Genomes: New Candidates and Distinguishing Features. Sci. Rep. 7, 41544 (2017).

16. Cancer Genome Atlas Research Network. Genomic and epigenomic landscapes of adult de novo acute myeloid leukemia. N. Engl. J. Med. 368, 2059–2074 (2013).

17. Campbell, P. J. et al. Pan-cancer analysis of whole genomes. (2017). doi:10.1101/162784

18. Perera, D. et al. Differential DNA repair underlies mutation hotspots at active promoters in cancer genomes. Nature 532, 259–263 (2016).

19. Sabarinathan, R., Mularoni, L., Deu-Pons, J., Gonzalez-Perez, A. & López-Bigas, N. Nucleotide excision repair is impaired by binding of transcription factors to DNA. Nature 532, 264–267 (2016).

20. Ellis, M. J. et al. Whole-genome analysis informs breast cancer response to aromatase inhibition. Nature 486, 353–360 (2012).

21. Brown, M. B. 400: A Method for Combining Non-Independent, One-Sided Tests of Significance. Biometrics 31, 987 (1975).

22. Fisher, R. A. Statistical Methods For Research Workers. (Genesis Publishing Pvt Ltd, 1925).

23. Benjamini, Y. & Hochberg, Y. Controlling the False Discovery Rate: A Practical and Powerful Approach to Multiple Testing. J. R. Stat. Soc. Series B Stat. Methodol. 57, 289–300 (1995).

24. Pasqualucci, L. et al. Hypermutation of multiple proto-oncogenes in B-cell diffuse large-cell lymphomas. Nature 412, 341–346 (2001).

25. Forbes, S. A. et al. COSMIC (the Catalogue of Somatic Mutations in Cancer): a resource to investigate acquired mutations in human cancer. Nucleic Acids Res. 38, D652–7 (2010).

26. Saeki, K. The B cell-specific major raft protein, Raftlin, is necessary for the integrity of lipid raft and BCR signal transduction. EMBO J. 22, 3015–3026 (2003).

27. ENCODE Project Consortium. An integrated encyclopedia of DNA elements in the human genome. Nature 489, 57–74 (2012).

28. Kotani, T., Akabane, S., Takeyasu, K., Ueda, T. & Takeuchi, N. Human G-proteins, ObgH1 and Mtg1, associate with the large mitochondrial ribosome subunit and are involved in translation and assembly of respiratory complexes. Nucleic Acids Res. 41, 3713–3722 (2013).

29. Kopan, R. & Ilagan, M. X. G. The canonical Notch signaling pathway: unfolding the activation mechanism. Cell 137, 216–233 (2009).

30. Dulak, A. M. et al. Exome and whole-genome sequencing of esophageal adenocarcinoma identifies recurrent driver events and mutational complexity. Nat. Genet. 45, 478–486 (2013).

31. Diaz-Lagares, A. et al. Epigenetic inactivation of the p53-induced long noncoding RNA TP53 target 1 in human cancer. Proc. Natl. Acad. Sci. U. S. A. 113, E7535–E7544 (2016).

32. Li, B.-S. et al. MicroRNA-25 promotes gastric cancer migration, invasion and proliferation by directly targeting transducer of ERBB2, 1 and correlates with poor survival. Oncogene 34, 2556–2565 (2015).

33. Hosoda, N. et al. Anti-proliferative protein Tob negatively regulates CPEB3 target by recruiting Caf1 deadenylase. EMBO J. 30, 1311–1323 (2011).

34. Morin, R. D. et al. Genetic Landscapes of Relapsed and Refractory Diffuse Large B-Cell Lymphomas. Clin. Cancer Res. 22, 2290–2300 (2016).

35. Chapuy, B. et al. Targetable genetic features of primary testicular and primary central nervous system lymphomas. Blood 127, 869–881 (2016).

36. Shadel, G. S. & Clayton, D. A. Mitochondrial DNA maintenance in vertebrates. Annu. Rev. Biochem. 66, 409–435 (1997).

37. Schmitt, M. E. & Clayton, D. A. Nuclear RNase MRP is required for correct processing of pre-5.8S rRNA in Saccharomyces cerevisiae. Mol. Cell. Biol. 13, 7935–7941 (1993).

38. Gill, T., Cai, T., Aulds, J., Wierzbicki, S. & Schmitt, M. E. RNase MRP cleaves the CLB2 mRNA to promote cell cycle progression: novel method of mRNA degradation. Mol. Cell. Biol. 24, 945–953 (2004).

39. Esakova, O. & Krasilnikov, A. S. Of proteins and RNA: the RNase P/MRP family. RNA 16, 1725–1747 (2010).

40. Sabarinathan R. E. al. RNAsnp: efficient detection of local RNA secondary structure changes induced by SNPs. - PubMed - NCBI. Available at: https://www.ncbi.nlm.nih.gov/pubmed/23315997. (accessed: 7th July 2017)

41. Mularoni, L., Sabarinathan, R., Deu-Pons, J., Gonzalez-Perez, A. & López-Bigas, N. OncodriveFML: a general framework to identify coding and non-coding regions with cancer driver mutations. Genome Biol. 17, 128 (2016).

42. Robbiani, D. F. et al. AID produces DNA double-strand breaks in non-Ig genes and mature B cell lymphomas with reciprocal chromosome translocations. Mol. Cell 36, 631–641 (2009).

43. Fujimoto, A. et al. Whole-genome mutational landscape and characterization of noncoding and structural mutations in liver cancer. Nat. Genet. 48, 500–509 (2016).

44. Lippert, M. J. et al. Role for topoisomerase 1 in transcription-associated mutagenesis in yeast. Proc. Natl. Acad. Sci. U. S. A. 108, 698–703 (2011).

45. Sankar, T. S., Wastuwidyaningtyas, B. D., Dong, Y., Lewis, S. A. & Wang, J. D. The nature of mutations induced by replication–transcription collisions. Nature 535, 178–181 (2016).

46. Jinks-Robertson, S. & Bhagwat, A. S. Transcription-associated mutagenesis. Annu. Rev. Genet. 48, 341–359 (2014).

47. Ke, H. et al. NEAT1 is Required for Survival of Breast Cancer Cells Through FUS and miR-548. Gene Regul. Syst. Bio. 10, 11–17 (2016).

48. Han, Y., Liu, Y., Nie, L., Gui, Y. & Cai, Z. Inducing Cell Proliferation Inhibition, Apoptosis, and Motility Reduction by Silencing Long Noncoding Ribonucleic Acid Metastasis-associated Lung Adenocarcinoma Transcript 1 in Urothelial Carcinoma of the Bladder. Urology 81, 209.e1–209.e7 (2013).

49. Dabney, J. & Meyer, M. Length and GC-biases during sequencing library amplification: a comparison of various polymerase-buffer systems with ancient and modern DNA sequencing libraries. Biotechniques 52, 87–94 (2012).

50. Lawrence, M. S. et al. Discovery and saturation analysis of cancer genes across 21 tumour types. Nature 505, 495–501 (2014).

51. Cibulskis, K. et al. Sensitive detection of somatic point mutations in impure and heterogeneous cancer samples. Nat. Biotechnol. 31, 213–219 (2013).

52. Carter, S. L. et al. Absolute quantification of somatic DNA alterations in human cancer. Nat. Biotechnol. 30, 413–421 (2012).

53. Martincorena, I. et al. Universal Patterns of Selection in Cancer and Somatic Tissues. Cell (2017). doi:10.1016/j.cell.2017.09.042

54. Lawrence, M. S. et al. Mutational heterogeneity in cancer and the search for new cancer-associated genes. Nature 499, 214–218 (2013).

55. Finucane, H. K. et al. Partitioning heritability by functional annotation using genome-wide association summary statistics. Nat. Genet. 47, 1228–1235 (2015).

56. Perera, D. et al. Differential DNA repair underlies mutation hotspots at active promoters in cancer genomes. Nature 532, 259–263 (2016).

57. Sabarinathan, R., Mularoni, L., Deu-Pons, J., Gonzalez-Perez, A. & López-Bigas, N. Nucleotide excision repair is impaired by binding of transcription factors to DNA. Nature 532, 264–267 (2016).

58. Flavahan, W. A. et al. Insulator dysfunction and oncogene activation in IDH mutant gliomas. Nature 529, 110–114 (2016).

59. Hnisz, D. et al. Activation of proto-oncogenes by disruption of chromosome neighborhoods. Science 351, 1454–1458 (2016).

## References

1. Ramos, A. H. et al. Oncotator: cancer variant annotation tool. Hum. Mutat. 36, E2423–9 (2015).

2. Kasar, S. et al. Whole-genome sequencing reveals activation-induced cytidine deaminase signatures during indolent chronic lymphocytic leukaemia evolution. Nat. Commun. 6, 8866 (2015).

3. Kim, J. et al. Somatic ERCC2 mutations are associated with a distinct genomic signature in urothelial tumors. Nat. Genet. 48, 600–606 (2016).

4. Polak, P. et al. A mutational signature reveals alterations underlying deficient homologous recombination repair in breast cancer. Nat. Genet. 49, 1476–1486 (2017).

5. Harrow, J. et al. GENCODE: the reference human genome annotation for The ENCODE Project. Genome Res. 22, 1760–1774 (2012).

6. Iyer, M. K. et al. The landscape of long noncoding RNAs in the human transcriptome. Nat. Genet. 47, 199–208 (2015).

7. Burge, S. W. et al. Rfam 11.0: 10 years of RNA families. Nucleic Acids Res. 41, D226–32 (2013).

8. Lowe, T. M. & Eddy, S. R. tRNAscan-SE: a program for improved detection of transfer RNA genes in genomic sequence. Nucleic Acids Res. 25, 955–964 (1997).

9. Lestrade, L. & Weber, M. J. snoRNA-LBME-db, a comprehensive database of human H/ACA and C/D box snoRNAs. Nucleic Acids Res. 34, D158–62 (2006).

10. Kozomara, A. & Griffiths-Jones, S. miRBase: annotating high confidence miRNAs using deep sequencing data. Nucleic Acids Res. 42, D68–D73 (2014).

11. Londin, E. et al. Analysis of 13 cell types reveals evidence for the expression of numerous novel primate- and tissue-specific microRNAs. Proc. Natl. Acad. Sci. U. S. A. 112, E1106–15 (2015).

12. Fingerman, I. M. et al. NCBI Epigenomics: a new public resource for exploring epigenomic data sets. Nucleic Acids Res. 39, D908–12 (2011).

13. Roadmap Epigenomics Consortium et al. Integrative analysis of 111 reference human epigenomes. Nature 518, 317–330 (2015).

14. Fu, Y. et al. FunSeq2: a framework for prioritizing noncoding regulatory variants in cancer. Genome Biol. 15, 480 (2014).

15. Alexandrov, L. B. et al. Signatures of mutational processes in human cancer. Nature 500, 415–421 (2013).

16. Polak, P. et al. Cell-of-origin chromatin organization shapes the mutational landscape of cancer. Nature 518, 360–364 (2015).

17. Benjamini, Y. & Hochberg, Y. Controlling the False Discovery Rate: A Practical and Powerful Approach to Multiple Testing. J. R. Stat. Soc. Series B Stat. Methodol. 57, 289–300 (1995).

18. Martincorena, I. et al. Universal Patterns of Selection in Cancer and Somatic Tissues. Cell (2017). doi:10.1016/j.cell.2017.09.042

19. Lanzos, A. et al. Discovery of Cancer Driver Long Noncoding RNAs across 1112 Tumour Genomes: New Candidates and Distinguishing Features. (2016). doi:10.1101/065805

20. Lochovsky, L., Zhang, J., Fu, Y., Khurana, E. & Gerstein, M. LARVA: an integrative framework for large-scale analysis of recurrent variants in noncoding annotations. Nucleic Acids Res. 43, 8123–8134 (2015).

21. Lawrence, M. S. et al. Discovery and saturation analysis of cancer genes across 21 tumour types. Nature 505, 495–501 (2014).

22. Nik-Zainal, S. et al. Landscape of somatic mutations in 560 breast cancer whole-genome sequences. Nature 534, 47–54 (2016).

23. Juul, M. et al. Non-coding cancer driver candidates identified with a sample- and position-specific model of the somatic mutation rate. Elife 6, (2017).

24. Mularoni, L., Sabarinathan, R., Deu-Pons, J., Gonzalez-Perez, A. & López-Bigas, N. OncodriveFML: a general framework to identify coding and non-coding regions with cancer driver mutations. Genome Biol. 17, 128 (2016).

25. Kircher, M. et al. A general framework for estimating the relative pathogenicity of human genetic variants. Nat. Genet. 46, 310–315 (2014).

26. Douville, C. et al. Assessing the Pathogenicity of Insertion and Deletion Variants with the Variant Effect Scoring Tool (VEST-Indel). Hum. Mutat. 37, 28–35 (2016).

27. Cibulskis, K. et al. Sensitive detection of somatic point mutations in impure and heterogeneous cancer samples. Nat. Biotechnol. 31, 213–219 (2013).

28. Anscombe, F. J. & Fisher, R. A. Statistical Methods for Research Workers. J. R. Stat. Soc. Ser. A 118, 486 (1955).

29. Brown, M. B. 400: A Method for Combining Non-Independent, One-Sided Tests of Significance. Biometrics 31, 987 (1975).

30. Poole, W., Gibbs, D. L., Shmulevich, I., Bernard, B. & Knijnenburg, T. A. Combining dependentP-values with an empirical adaptation of Brown’s method. Bioinformatics 32, i430–i436 (2016).

31. Poole, W., Gibbs, D. L., Shmulevich, I., Bernard, B. & Knijnenburg, T. Combining Dependent P-values with an Empirical Adaptation of Brown’s Method. (2015). doi:10.1101/029637

32. Derrien, T. et al. Fast computation and applications of genome mappability. PLoS One 7, e30377 (2012).

33. Sabarinathan, R., Mularoni, L., Deu-Pons, J., Gonzalez-Perez, A. & López-Bigas, N. Nucleotide excision repair is impaired by binding of transcription factors to DNA. Nature 532, 264–267 (2016).

34. Perera, D. et al. Differential DNA repair underlies mutation hotspots at active promoters in cancer genomes. Nature 532, 259–263 (2016).

35. Fredriksson, J., Eliot, K., Filges, S., Stahlberg, A. & Larsson, E. Recurrent promoter mutations in melanoma are defined by an extended context-specific mutational signature. (2016). doi:10.1101/069351

36. Pearson, C. E., Zorbas, H., Price, G. B. & Zannis-Hadjopoulos, M. Inverted repeats, stem-loops, and cruciforms: significance for initiation of DNA replication. J. Cell. Biochem. 63, 1–22 (1996).

37. Nik-Zainal, S. et al. Landscape of somatic mutations in 560 breast cancer whole-genome sequences. Nature 534, 47–54 (2016).

38. Ye, C., Ji, G., Li, L. & Liang, C. detectIR: a novel program for detecting perfect and imperfect inverted repeats using complex numbers and vector calculation. PLoS One 9, e113349 (2014).

39. Fonseca, N. A. et al. Pan-cancer study of heterogeneous RNA aberrations. bioRxiv 183889 (2017). doi:10.1101/183889

40. Dobin, A. et al. STAR: ultrafast universal RNA-seq aligner. Bioinformatics 29, 15–21 (2013).

41. Kim, D. et al. TopHat2: accurate alignment of transcriptomes in the presence of insertions, deletions and gene fusions. Genome Biol. 14, R36 (2013).

42. Hose, J. et al. Dosage compensation can buffer copy-number variation in wild yeast. Elife 4, (2015).

43. Lockstone, H. E. et al. Gene expression profiling in the adult Down syndrome brain. Genomics 90, 647–660 (2007).

44. Birchler, J. A. Reflections on studies of gene expression in aneuploids. Biochem. J 426, 119–123 (2010).

45. Heard, E. & Disteche, C. M. Dosage compensation in mammals: fine-tuning the expression of the X chromosome. Genes Dev. 20, 1848–1867 (2006).

46. Mermel, C. H. et al. GISTIC2.0 facilitates sensitive and confident localization of the targets of focal somatic copy-number alteration in human cancers. Genome Biol. 12, R41 (2011).

47. Thorvaldsdóttir, H., Robinson, J. T. & Mesirov, J. P. Integrative Genomics Viewer (IGV): high-performance genomics data visualization and exploration. Brief. Bioinform. 14, 178–192 (2013).

48. Carter, S. L. et al. Absolute quantification of somatic DNA alterations in human cancer. Nat. Biotechnol. 30, 413–421 (2012).

49. Rheinbay, E. et al. Recurrent and functional regulatory mutations in breast cancer. Nature 547, 55–60 (2017).

50. Martincorena, I. et al. Universal Patterns of Selection in Cancer and Somatic Tissues. Cell (2017). doi:10.1016/j.cell.2017.09.042

51. Zack, T. I. et al. Pan-cancer patterns of somatic copy number alteration. Nat. Genet. 45, 1134–1140 (2013).

52. Katz, Y., Wang, E. T., Airoldi, E. M. & Burge, C. B. Analysis and design of RNA sequencing experiments for identifying isoform regulation. Nat. Methods 7, 1009–1015 (2010).

53. Sabarinathan, R. et al. RNAsnp: efficient detection of local RNA secondary structure changes induced by SNPs. Hum. Mutat. 34, 546–556 (2013).

54. Nawrocki, E. P. et al. Rfam 12.0: updates to the RNA families database. Nucleic Acids Res. 43, D130–7 (2015).

55. Weinreb, C. et al. 3D RNA and Functional Interactions from Evolutionary Couplings. Cell 165, 963–975 (2016).

56. Perederina, A., Esakova, O., Quan, C., Khanova, E. & Krasilnikov, A. S. Eukaryotic ribonucleases P/MRP: the crystal structure of the P3 domain. EMBO J. 29, 761–769 (2010).

57. Khanova, E., Esakova, O., Perederina, A., Berezin, I. & Krasilnikov, A. S. Structural organizations of yeast RNase P and RNase MRP holoenzymes as revealed by UV-crosslinking studies of RNA–protein interactions. RNA 18, 720–728 (2012).

58. Esakova, O., Perederina, A., Berezin, I. & Krasilnikov, A. S. Conserved regions of ribonucleoprotein ribonuclease MRP are involved in interactions with its substrate. Nucleic Acids Res. 41, 7084–7091 (2013).

59. Welter, D. et al. The NHGRI GWAS Catalog, a curated resource of SNP-trait associations. Nucleic Acids Res. 42, D1001–6 (2014).

60. Sud, A., Kinnersley, B. & Houlston, R. S. Genome-wide association studies of cancer: current insights and future perspectives. Nat. Rev. Cancer 17, 692–704 (2017).

## References

1. Alexandrov, L. B. et al. Signatures of mutational processes in human cancer. Nature 500, 415–421 (2013).

2. Katainen, R. et al. CTCF/cohesin-binding sites are frequently mutated in cancer. Nat. Genet. 47, 818–821 (2015).

3. Weinhold, N., Jacobsen, A., Schultz, N., Sander, C. & Lee, W. Genome-wide analysis of noncoding regulatory mutations in cancer. Nat. Genet. 46, 1160–1165 (2014).

4. Nik-Zainal, S. et al. Landscape of somatic mutations in 560 breast cancer whole-genome sequences. Nature 534, 47–54 (2016).

5. Khurana, E. et al. Integrative annotation of variants from 1092 humans: application to cancer genomics. Science 342, 1235587 (2013).

6. Fujimoto, A. et al. Whole-genome mutational landscape and characterization of noncoding and structural mutations in liver cancer. Nat. Genet. 48, 500–509 (2016).

7. Yi, C. et al. MiR-663, a microRNA targeting p21(WAF1/CIP1), promotes the proliferation and tumorigenesis of nasopharyngeal carcinoma. Oncogene 31, 4421–4433 (2012).

8. Baer M. E. al. Structure and transcription of a human gene for H1 RNA, the RNA component of human RNase P. - PubMed - NCBI. Available at: https://www.ncbi.nlm.nih.gov/pubmed/2308839.(Accessed: 7th July 2017)

9. Iyer, M. K. et al. The landscape of long noncoding RNAs in the human transcriptome. Nat. Genet. 47, 199–208 (2015).

10. Wang, L. et al. Linc00963: a novel, long non-coding RNA involved in the transition of prostate cancer from androgen-dependence to androgen-independence. Int. J. Oncol. 44, 2041–2049 (2014).

11. Harrow, J. et al. GENCODE: the reference human genome annotation for The ENCODE Project. Genome Res. 22, 1760–1774 (2012).

12. Katz, Y., Wang, E. T., Airoldi, E. M. & Burge, C. B. Analysis and design of RNA sequencing experiments for identifying isoform regulation. Nat. Methods 7, 1009–1015 (2010).

13. Martincorena, I. et al. Universal Patterns of Selection in Cancer and Somatic Tissues. Cell 171, 1029–1041.e21 (2017).

